# Co-activation of selective nicotinic acetylcholine receptor subtypes is required for neuroprotection against Alzheimer’s disease

**DOI:** 10.1101/2024.07.08.602576

**Authors:** Rahmi Lee, Gayeon Kim, Ellison R. Black, Seonil Kim

## Abstract

Beta-amyloid peptide (Aβ)-induced suppression of hippocampal GABAergic interneuron activity drives hyperexcitability, amyloid pathology, and cognitive impairment in Alzheimer’s disease (AD), suggesting that enhancing hippocampal inhibition may be protective. However, hippocampal interneurons are highly diverse and differentially regulate inhibition and cognition, making it challenging to identify the affected subtypes and optimally restore hippocampal inhibition in AD. We have previously found that Aβ selectively binds to two of the three major hippocampal nicotinic acetylcholine receptor (nAChR) subtypes, α7- and α4β2-nAChRs, but not α3β4-nAChRs, and inhibits these two receptors in hippocampal inhibitory interneurons to decrease their activity, leading to hyperexcitation in excitatory neurons. Here, we further reveal that α7- and α4β2-nAChRs predominantly control the nicotinic cholinergic signaling and neuronal activity in hippocampal parvalbumin-positive (PV+) and somatostatin-positive (SST+) inhibitory interneurons, respectively. We also find that systemic co-stimulation of α7- and α4β2-nAChRs is required to reverse hippocampal hyperexcitability, dysfunction of fear learning-associated hippocampal oscillatory activity, and hippocampus-dependent memory loss and reduce Aβ pathology in AD model mice. This suggests that co-stimulation of PV+ and SST+ cells via activation of α7- and α4β2-nAChRs together is required to enhance hippocampal inhibition optimally, which reverses hippocampal dysfunction, reduces amyloid pathology, and ultimately prevents memory loss in AD.

**Significance Statement:** Development of a safe and effective unified strategy to counteract hyperexcitability holds a significant promise in advancing Alzheimer’s disease (AD) treatment. Studies suggest that beta-amyloid peptide (Aβ) is a major trigger of hyperexcitability. However, the mechanisms of how Aβ could participate in hyperexcitability are still not fully understood. Here, we reveal that Aβ inhibits distinct hippocampal nicotinic acetylcholine receptor (nAChR) subtypes in different hippocampal inhibitory interneuron subtypes, which leads to hyperexcitability, resulting in amyloid pathology and memory loss in AD. We further discover that systemic co-stimulation of these nAChRs is required for neuroprotection against AD at the cellular, circuit, network, and behavioral levels. Identifying this mechanism can thus provide a new therapeutic strategy in AD.

## Introduction

About 5.8 million Americans currently have Alzheimer’s disease (AD); that figure is projected to reach 14 million people by 2050 (Wong 2020). In 2020, the direct costs to American society of caring for those with AD and related dementias are a total of an estimated $305 billion, which makes AD the most expensive disease in America (Wong 2020). However, there is no known cause and cure for AD (Walsh and Selkoe 2020). Therefore, understanding the neural mechanisms of AD pathogenesis is of the utmost importance for better diagnostic and therapeutic development.

Both preclinical and clinical studies demonstrate that cortical and hippocampal hyperactivity emerges early in Alzheimer’s disease (AD) and is closely associated with cognitive decline (Targa Dias Anastacio et al. 2022; Yuan and Grutzendler 2016; Vossel et al. 2016; Mondadori et al. 2006). Elevated neuronal activity promotes the release of both β-amyloid (Aβ) (Cirrito et al. 2005; Yamamoto et al. 2015) and tau (Pooler et al. 2013; Wu et al. 2016), thereby accelerating AD pathology. Importantly, reducing hippocampal hyperactivity with antiepileptic drugs improves cognition in both AD mouse models (Sanchez et al. 2012; Shi et al. 2013; Zheng et al. 2022) and individuals with mild cognitive impairment (Bakker et al. 2012; Vossel et al. 2021).

These findings suggest that neuronal hyperexcitability is a key driver of AD pathology and cognitive dysfunction, making it an attractive target for both symptomatic and disease-modifying therapies. However, the underlying circuit and cell type-specific mechanisms linking hyperexcitability to AD pathology and cognitive impairment remain poorly understood.

The hippocampal memory system is particularly vulnerable to AD-related cellular pathology (Hyman and Trojanowski 1997). In the hippocampus, Aβ is shown to decrease the activity of inhibitory cells to increase excitation in pyramidal cells (Palop and Mucke 2010). This likely results in hippocampal hyperexcitability and network dysfunction, leading to cognitive decline in AD (Palop and Mucke 2010; Kazim et al. 2021; Targa Dias Anastacio et al. 2022). This suggests that the reduction of hippocampal inhibition by Aβ is a crucial trigger for hyperexcitability-induced cognitive impairment in AD. Hence, enhancing hippocampal inhibition is thought to be protective against AD (Palop and Mucke 2016). However, Aβ unlikely affects all subtypes of hippocampal interneurons equally (Ren et al. 2018; Caccavano et al. 2020; Xu et al. 2020). Therefore, identifying the affected interneuron subtypes in AD to enhance hippocampal inhibition optimally is conceptually and technically challenging.

Cholinergic regulation of GABAergic inhibitory network is generally stronger than direct actions on excitatory neurons in the hippocampus since nicotinic acetylcholine receptors (nAChRs) are more densely expressed in inhibitory interneurons than excitatory cells (Ji et al. 2001; Frazier, Rollins, et al. 1998; Hefft et al. 1999; Jones and Yakel 1997; McQuiston and Madison 1999; Ji and Dani 2000). The loss of cholinergic neurons and nAChR expression in the hippocampus is a prominent AD pathology (Nordberg 2001; Kadir et al. 2006; Ferreira-Vieira et al. 2016; Pensalfini et al. 2020; Alam and Nixon 2021; Jiang et al. 2016; Chen et al. 2009; Ginsberg et al. 2011). This suggests that cholinergic dysfunction is likely a primary cause for hippocampal inhibition deficits in AD. Although nAChRs are high-affinity targets for Aβ (Jurgensen and Ferreira 2010), contradictory results have been reported describing effects of Aβ on nAChRs (Liu et al. 2001; Pettit et al. 2001; Pym et al. 2005; George et al. 2021). Moreover, 10-15 subtypes of neuronal nAChRs have been reported in human brains (Akaike and Izumi 2018). In fact, FDA-approved current drugs for AD mostly attempt to delay the general breakdown of acetylcholine, which potentially stimulates all types of acetylcholine receptors. Thus, it is not surprising that these non-selective activators suffer from modest efficacy (Ferreira-Vieira et al. 2016; Thompson et al. 2004; Mangialasche et al. 2010). This suggests, in turn, that distinct nAChR subtypes are differentially affected in AD. Thus, it is challenging to comprehend how Aβ impacts nAChR functions in AD. Furthermore, the important question of how Aβ-induced subtype-specific disruptions in hippocampal interneuron functions are linked to distinct nAChR subtypes remains unanswered.

The three major nAChR subtypes in the hippocampus are composed of α7, α4β2, and α3β4 subunit combinations (Albuquerque et al. 2009; Lindstrom 2003; Clementi et al. 2000). Importantly, our previous work using Ca^2+^ imaging in cultured hippocampal neurons has shown that Aβ selectively inhibits α7- and α4β2-nAChRs together, but not α3β4-nAChRs, which reduces overall neuronal activity in interneurons, resulting in neuronal hyperexcitation in excitatory cells (Sun et al. 2019; Roberts et al. 2021). Additionally, our published data using co-immunoprecipitation have shown that Aβ can specifically bind to α7- and α4-containing nAChRs, but not α3-containing receptors (Roberts et al. 2021). We have also identified two amino acids that are critical for the interaction of Aβ with α7 and α4 subunits, but not the α3 subunit, providing the molecular mechanism underlying the selective interaction of Aβ with specific nAChRs (Roberts et al. 2021). In line with these findings, considerable evidence suggests that Aβ is shown to exert subtype-specific inhibition of α7- and/or α4β2-nAChR function without affecting α3β4-nAChRs (Pettit et al. 2001; Liu et al. 2001; Grassi et al. 2003; Wu et al. 2004; Pym et al. 2005; Magdesian et al. 2005; Olivero et al. 2014; Sun et al. 2019; Jurgensen and Ferreira 2010). α7 and α4 subtypes are also more significantly reduced in the brains of AD patients compared to α3-type receptors (Guan et al. 2000). This supports the idea that Aβ predominantly reduces neuronal activity in inhibitory interneurons via selective inhibition of α7- and α4β2-nAChRs, but not α3β4-nAChRs (Sun et al. 2019). We have further revealed that co-activation of α7- and α4β2-nAChRs is required to reverse the Aβ-induced adverse effects in hippocampal excitatory and inhibitory cells (Roberts et al. 2021; Sun et al. 2019). Moreover, non-selective stimulation of acetylcholine receptors exacerbates Aβ effects on cultured neurons (Roberts et al. 2021; Sun et al. 2019). Therefore, our published works have shown that selective co-activation of α7- and α4β2-nAChRs is required to reverse the Aβ effects on the activity of inhibitory interneurons in AD. Nonetheless, it is still unknown how the Aβ effects on distinct nAChR subtypes differentially affect inhibitory interneurons to disrupt hippocampal activity at the local circuits and network, resulting in amyloid pathology and cognitive decline in AD.

Parvalbumin-positive (PV+) and somatostatin-positive (SST+) cells, two major subtypes of inhibitory interneurons in the hippocampus, mainly provide somatic and dendritic inhibition onto pyramidal cells, respectively, to differentially regulate hippocampal network activity and cognitive processes (Pelkey et al. 2017). Here, we reveal that α7-, α4β2-, and α3β4-nAChRs regulate the cholinergic activity mainly in PV+, SST+, and excitatory cells, respectively. As PV+ and SST+ cells differentially provide inhibition onto excitatory neurons in the hippocampus, co-activation of these cells by stimulating α7- and α4β2-nAChRs together has significant impacts on hippocampal inhibition at the circuit levels. Indeed, we find that systemic co-stimulation of α7- and α4β2-nAChRs is required to reverse hippocampal hyperexcitability, dysfunction of fear learning-associated hippocampal oscillatory activity, and fear memory loss and reduce Aβ pathology in a widely used amyloid pathology mouse model, 5XFAD mice (Oakley et al. 2006). This suggests that co-stimulation of PV+ and SST+ cells via activation of α7- and α4β2-nAChRs together is required to enhance hippocampal inhibition optimally, which reverses hippocampal dysfunction, reduces amyloid pathology, and ultimately prevents memory loss in AD.

## Results

### Nicotinic cholinergic regulation of cultured hippocampal PV+ and SST+ interneurons is mainly mediated by α7- and α4β2-nAChRs, respectively

To determine whether nicotinic cholinergic regulation was stronger in GABAergic inhibitory cells than in excitatory neurons, we first compared nicotinic cholinergic activity between excitatory cells (EX) and inhibitory interneurons (IN) using Ca^2+^ imaging with nicotine uncaging in 12-14 days *in vitro* (DIV) cultured mouse hippocampal neurons as neuronal nAChRs are ligand-gated ion channels, and when activated, they induce membrane depolarization and increase somatic Ca^2+^ levels (Banala et al. 2018). We found that nicotinic cholinergic activity measured by Ca^2+^ transients in inhibitory interneurons was significantly higher than in excitatory neurons (EX, 1.000 ± 0.879 ΔF/F_0_ and IN, 1.575 ± 1.208 ΔF/F_0_, *p* = 0.0109) (**Fig. 1a and Table S1**). This finding is consistent with the previous reports that cholinergic regulation of inhibitory cells is stronger than excitatory neurons in the hippocampus (Ji et al. 2001; Frazier, Rollins, et al. 1998; Hefft et al. 1999; Jones and Yakel 1997; McQuiston and Madison 1999; Ji and Dani 2000).

**Figure 1.**
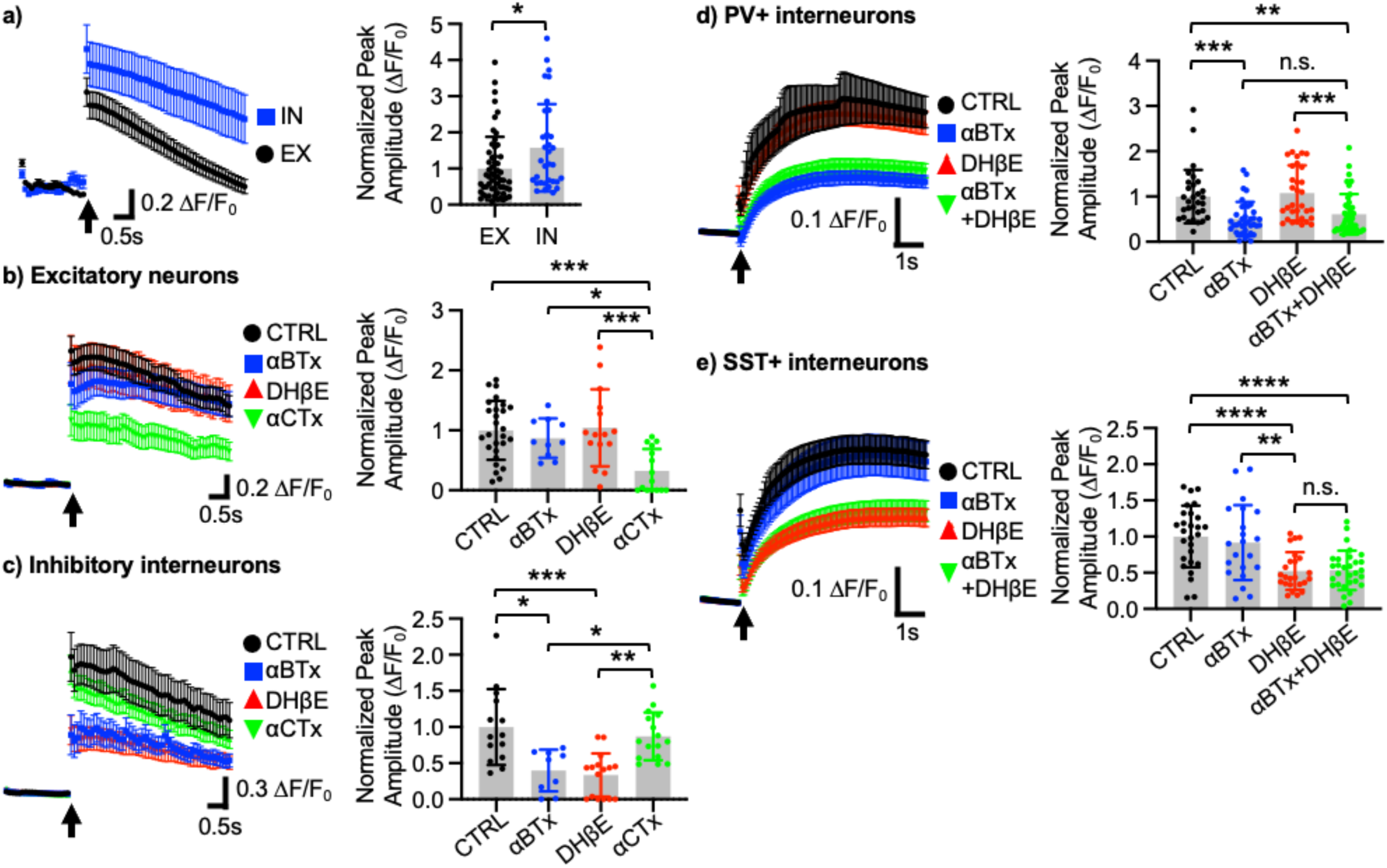
Nicotinic cholinergic regulation of cultured hippocampal PV+ and SST+ interneurons is mainly mediated by α7- and α4β2-nAChRs, respectively. **a)** Average traces of GCaMP6f signals in excitatory (EX) and inhibitory (IN) cells and summary data of normalized peak amplitude in each condition (n = number of cells excitatory cells = 57 and inhibitory cells = 33 from 4 pups in 2 independent cultures). \**p* < 0.05, unpaired two-tailed student’s t-test. Average traces of GCaMP6f signals and summary graphs of normalized peak amplitude in each condition in **b)** excitatory neurons (n = number of cells, CTRL = 27, αBTx = 10, DhβE = 15, and αCTx = 15 from 4 pups in 2 independent cultures) and **c)** inhibitory interneurons (n = number of cells, CTRL = 14, αBTx = 9, DhβE = 16, and αCTx = 16 from 4 pups in 2 independent cultures). Average traces of GCaMP7s signals and summary data of normalized peak amplitude in each condition in **d)** PV+ interneurons (n = number of cells, CTRL = 30, αBTx = 38, DhβE = 32, and αBTx + DhβE = 43 from 6 pups in 3 independent cultures) and **e)** SST+ interneurons (n = number of cells, CTRL = 27, αBTx = 21, DhβE = 24, and αBTx + DhβE = 30 from 6 pups in 3 independent cultures). \**p* < 0.05, \*\**p* < 0.01, \*\*\**p* < 0.001, and \*\*\*\**p* < 0.0001. one-way ANOVA, Tukey test. An arrow indicates photostimulation. n.s.: not significant.

We next treated 12-14 DIV cultured mouse hippocampal neurons with nAChR subtype-specific antagonists and conducted Ca^2+^ imaging with nicotine uncaging to identify which subtypes of nAChRs played a crucial role in nicotinic cholinergic regulation of hippocampal excitatory and inhibitory cells. In excitatory cells, when compared to controls (CTRL), acute treatment of 3 μM α-Conotoxin AuIB (αCTx), an α3β4-receptor inhibitor, significantly reduced nicotinic cholinergic activity, while 50 nM α-Bungarotoxin (αBTx), an α7-receptor inhibitor, or 1 μM Dihydro-β-erythroidine hydrobromide (DHβE), an α4β2-receptor inhibitor, had no effects (CTRL, 1.000 ± 0.492 ΔF/F_0_, αBTx, 0.870 ± 0.328 ΔF/F_0_, *p* = 0.8881, DHβE, 1.042 ± 0.644 ΔF/F_0_, *p* = 0.9932, and αCTx, 0.325 ± 0.362 ΔF/F_0_, *p* = 0.0003) (**Fig. 1b and Table S2**). Conversely, in inhibitory cells, when compared to controls (CTRL), αBTx or DHβE treatment significantly reduced nicotine-induced activity, but the α3β4 antagonist had no effect (CTRL, 1.000 ± 0.525 ΔF/F_0_, αBTx, 0.398 ± 0.290 ΔF/F_0_, *p* = 0.0025, DHβE, 0.335 ± 0.300 ΔF/F_0_, *p* < 0.0001, and αCTx, 0.871 ± 0.330 ΔF/F_0_, *p* = 0.7866) (**Fig. 1c and Table S2**). This data show that α7- and α4β2-nAChRs are mainly responsible for nicotinic cholinergic activity in hippocampal inhibitory cells, while α3β4-nAChRs are important for nicotinic cholinergic activity in hippocampal excitatory cells. These findings are consistent with the previous reports demonstrating that postsynaptic α7- and α4β2-nAChRs are mainly localized on GABAergic interneurons in the hippocampus and mediate nicotine currents (Jones and Yakel 1997; Frazier, Rollins, et al. 1998), while they are expressed in the presynaptic area of excitatory cells to regulate neurotransmitter release without neuronal activation such as action potentials (Radcliffe and Dani 1998; Alkondon et al. 1999; Alkondon and Albuquerque 2001; Maggi et al. 2001; Huang et al. 2010). This suggests that stimulation of α7- and α4β2-nAChRs has more profound effects on neuronal activation in inhibitory cells than excitatory neurons in the hippocampus.

We next analyzed subtype-specific roles of nAChRs in cultured hippocampal PV+ and SST+ interneurons. Since α3β4-nAChRs are shown to mainly regulate excitatory neurons, we focused on the role of α7- and α4β2-nAChRs in these inhibitory cells. In PV+ interneurons, when compared to controls (CTRL), acute treatment of αBTx significantly reduced nicotinic cholinergic activity, while DHβE had no effects (CTRL, 1.000 ± 0.590 ΔF/F_0_, αBTx, 0.507 ± 0.374 ΔF/F_0_, *p* = 0.0006, and DHβE, 1.076 ± 0.613 ΔF/F_0_, *p* = 0.9340) (**Fig. 1d and Table S3**). Importantly, co-treatment of αBTx and DHβE had no additional effect (αBTx + DHβE, 0.607 ± 0.448 ΔF/F_0_) (**Fig. 1d and Table S3**). Conversely, in SST+ interneurons, when compared to controls (CTRL), acute treatment of DHβE significantly reduced nicotinic cholinergic activity, while αBTx had no effects (CTRL, 1.000 ± 0.426 ΔF/F_0_, αBTx, 0.917 ± 0.519 ΔF/F_0_, *p* = 0.8707, and DHβE, 0.524 ± 0.262 ΔF/F_0_, *p* < 0.0001) (**Fig. 1e and Table S3**). Importantly, co-treatment of αBTx and DHβE had no additional effect (αBTx + DHβE, 0.533 ± 0.272 ΔF/F_0_) (**Fig. 1e and Table S3**). These findings show that α7- and α4β2-nAChRs play important roles in nicotinic cholinergic regulation of PV+ and SST+ cells, respectively.

### The Aβ-induced decrease in the neuronal activity of PV+ and SST+ interneurons is reversed by stimulation of α7- and α4β2-nAChRs, respectively

We next examined whether Aβ affected nicotinic cholinergic activity of PV+ and SST+ interneurons using nicotine uncaging as described in **Fig. 1**. We found that 250 nM soluble Aβ42 oligomers (oAβ42) treatment significantly reduced nicotine-induced activity in both PV+ and SST+ interneurons compared to neurons treated with 250 nM scrambled Aβ42 (sAβ42) (PV+ interneurons, sAβ42, 1.000 ± 0.772 ΔF/F_0_ and oAβ42, 0.448 ± 0.233 ΔF/F_0_, *p* = 0.0038 and SST+ interneurons, sAβ42, 1.000 ± 0.391 ΔF/F_0_ and oAβ42, 0.600 ± 0.237 ΔF/F_0_, *p* = 0.0002) (**Fig. 2a-b and Table S4**). We further determined how Aβ affected neuronal activity in PV+ and SST+ interneurons. As somatic Ca^2+^ levels in neurons indicate neuronal activity (Gleichmann and Mattson 2011), we measured spontaneous somatic Ca^2+^ activity without tetrodotoxin (TTX) in 12-14 DIV cultured hippocampal neurons as carried out previously (Roberts et al. 2021; Sun et al. 2019; Sztukowski et al. 2018; Zaytseva et al. 2023). 250 nM oAβ42 treatment significantly decreased neuronal activity in both PV+ and SST+ interneurons (PV+ interneurons, sAβ42, 1.000 ± 0.379 ΔF/F_min_ and oAβ42, 0.589 ± 0.223 ΔF/F_min_, *p* = 0.0371, and SST+ interneurons, sAβ42, 1.000 ± 0.499 ΔF/F_min_ and oAβ42, 0.604 ± 0.295 ΔF/F_min_, *p* = 0.0044) (**Fig. 2c-d and Table S5 and S6**). To determine subtype-specific roles of nAChRs in the Aβ effect on PV+ and SST+ cells’ activity, we treated these cells with 250 nM oAβ42 or 250 nM sAβ42 and subtype-specific nAChR agonists - 1 μM PNU-282987 (PNU), an α7 agonist, 2 μM RJR-2403 Oxalate (RJR), an α4β2 agonist, or 1 μM NS-3861 (NS), an α3β4 agonist, as carried out previously (Roberts et al. 2021). In PV+ interneurons, when compared to controls, activation of α7-nAChRs significantly reversed the Aβ effects, while stimulation of other nAChR subtype had no effects (oAβ42 + PNU, 1.408 ± 0.806 ΔF/F_min_, *p* < 0.0001, oAβ42 + RJR, 0.528 ± 0.299 ΔF/F_min_, *p* > 0.9999, and oAβ42 + NS, 0.550 ± 0.384 ΔF/F_min_, *p* > 0.9999) (**Fig. 2c and Table S5**). In SST+ interneurons, when compared to controls, stimulation of α4β2-nAChRs reversed the Aβ effects, but other agonists were unable to reverse the Aβ effects (oAβ42 + PNU, 0.557 ± 0.271 ΔF/F_min_, *p* = 0.9900, oAβ42 + RJR, 0.994 ± 0.358 ΔF/F_min_, *p* = 0.0103, and oAβ42 + NS, 0.561 ± 0.249 ΔF/F_min_, *p* > 0.9999) (**Fig. 2d and Table S6**). We discover that the Aβ-induced decrease in the neuronal activity of PV+ and SST+ interneurons is reversed by stimulation of α7- and α4β2-nAChRs, respectively. Additionally, our findings show that α3β4-nAChRs are important for nicotinic cholinergic activity in excitatory cells. This suggests that stimulation of α7- and α4β2-nAChRs is a novel and ideal way to stimulate PV+ and SST+ interneurons’ activity rather than excitatory neurons’ activity.

**Figure 2.**
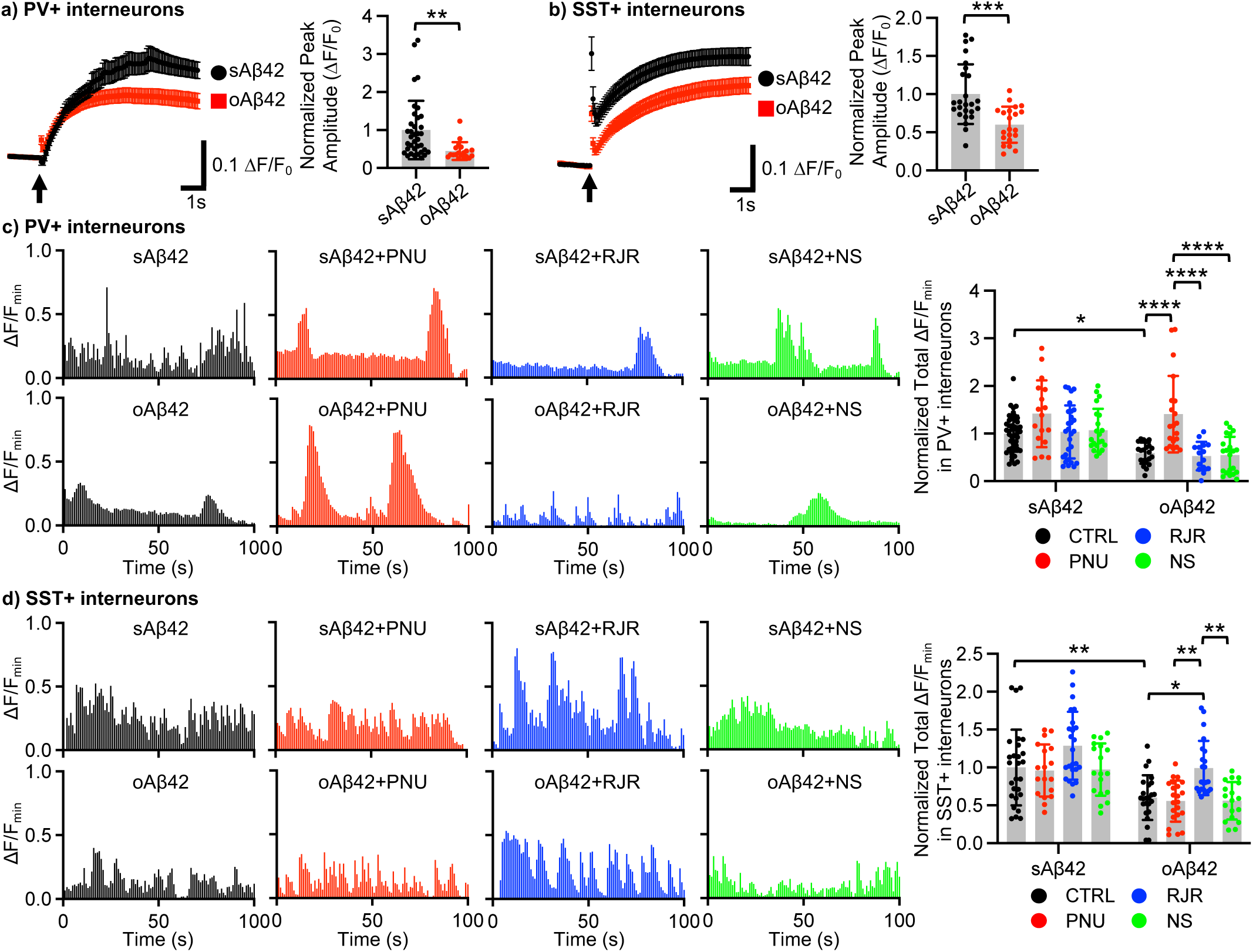
The Aβ-induced decrease in the neuronal activity of PV+ and SST+ interneurons is reversed by stimulation of α7- and α4β2-nAChRs, respectively. Average traces of GCaMP7s signals and summary data of normalized peak amplitude in each condition in **a)** PV+ interneurons (n = number of cells, sAβ42 = 36 and oAβ42 = 19 from 6 pups in 3 independent cultures) and **b)** SST+ interneurons (n = number of cells, sAβ42 = 25 and oAβ42 = 21 from 6 pups in 3 independent cultures). \*\**p* < 0.01 and \*\*\**p* < 0.001, unpaired two-tailed student’s t-test. An arrow indicates photostimulation. Representative traces of GCaMP7s fluorescence intensity and summary graphs of normalized total Ca^2+^ activity in each condition in **c)** PV+ interneurons (n = number of cells, sAβ42 (CTRL) = 47, sAβ42 + PNU = 17, sAβ42 + RJR = 28, sAβ42 + NS = 21, oAβ42 (CTRL) = 21, oAβ42 + PNU = 20, oAβ42 + RJR = 15, and oAβ42 + NS = 21 from 6 pups in 3 independent cultures) and **d)** SST+ interneurons (n = number of cells, sAβ42 (CTRL) = 27, sAβ42 + PNU = 18, sAβ42 + RJR = 24, sAβ42 + NS = 18, oAβ42 (CTRL) = 24, oAβ42 + PNU = 22, oAβ42 + RJR = 22, and oAβ42 +NS = 18 from 6 pups in 3 independent cultures). \**p* < 0.05, \*\**p* < 0.01, and \*\*\*\**p* < 0.0001, Two-way ANOVA, Tukey test.

### Co-stimulation of PV+ and SST+ interneurons by activating α7- and α4β2-nAChRs together is required to reverse hippocampal hyperexcitability in 5XFAD mice

Previous studies suggest that a reduction of GABAergic inhibitory interneurons’ activity in the hippocampus by Aβ is one of the main causes of neuronal hyperexcitability in AD (Palop and Mucke 2010; Kazim et al. 2021; Targa Dias Anastacio et al. 2022; Gallego-Rudolf et al. 2024a). However, cell-type specificity is absent in these studies, thus it is essential to determine cell-type specific changes of neuronal activity in the AD hippocampus. We measured the expression of the activity-regulated gene, c-Fos (Greenberg and Ziff 1984), to determine if the activity of PV+ and SST+ cells was reduced while pyramidal neurons were hyperexcitable in the hippocampus in AD pathology animals. We then examined whether co-stimulation of PV+ and SST+ cells via activating α7- and α4β2-nAChRs together is necessary for reducing hyperexcitability in hippocampal excitatory cells in the AD hippocampus in mice. Widely used amyloid pathology model mice, 5XFAD, were used because they show hippocampal hyperexcitability as early as 2.5 months of age (Li et al. 2021). Hippocampal sections of 5-month-old 5XFAD and wild-type (WT) female and male were stained with anti-PV, anti-SST, and anti-c-Fos antibodies, and DAPI was used to identify nuclei in pyramidal cells (**Fig. 3a)**. We then counted hippocampal c-Fos-positive (c-Fos+) cells in pyramidal, PV+, and SST+ cells per 5.0 × 10^6^ μm^2^ and compared them in each condition. First, we found a significant elevation of the number of c-Fos+ pyramidal cells in the hippocampus of both female and male 5XFAD mice when compared to WT animals (Female; WT + Saline, 69.845 ± 27.943 and 5XFAD + Saline, 150.818 ± 56.931, *p* < 0.0001, Male; WT + Saline, 65.166 ± 28.820 and 5XFAD + Saline, 143.918 ± 47.784, *p* < 0.0001) (**Fig. 3b, Fig. S1, and Table S7**), indicating hyperexcitability. To test whether co-activation α7- and α4β2-nAChRs was required to reverse hippocampal hyperexcitation in 5XFAD mice, we used subtype-specific nAChR agonists, PNU and RJR. We first treated PNU to animals for examining whether stimulating α7-nAChRs was sufficient to reverse hippocampal hyperexcitability. PNU treatment was unable to reverse hyperexcitation in the female 5XFAD hippocampus while it had a slight but significant effect on the activation of hippocampal pyramidal cells in male 5XFAD mice (Female; 5XFAD + PNU, 121.873 ± 76.511, *p* = 0.6233 and Male; 5XFAD + PNU, 110.495 ± 34.132, *p* = 0.0013) although these neurons were still hyperexcitable when compared to cells in the male WT hippocampus (*p* = 0.0104) (**Fig. 3b, Fig. S1, and Table S7**). RJR treatment for activation of α4β2-nAChRs was incapable of reversing hippocampal hyperexcitability in 5XFAD mice (Female; 5XFAD + RJR, 127.829 ± 59.897, *p* = 0.2577 and Male; 5XFAD + RJR, 131.236 ± 53.291, *p* = 0.3558 (**Fig. 3b, Fig. S1, and Table S7**).

**Figure 3.**
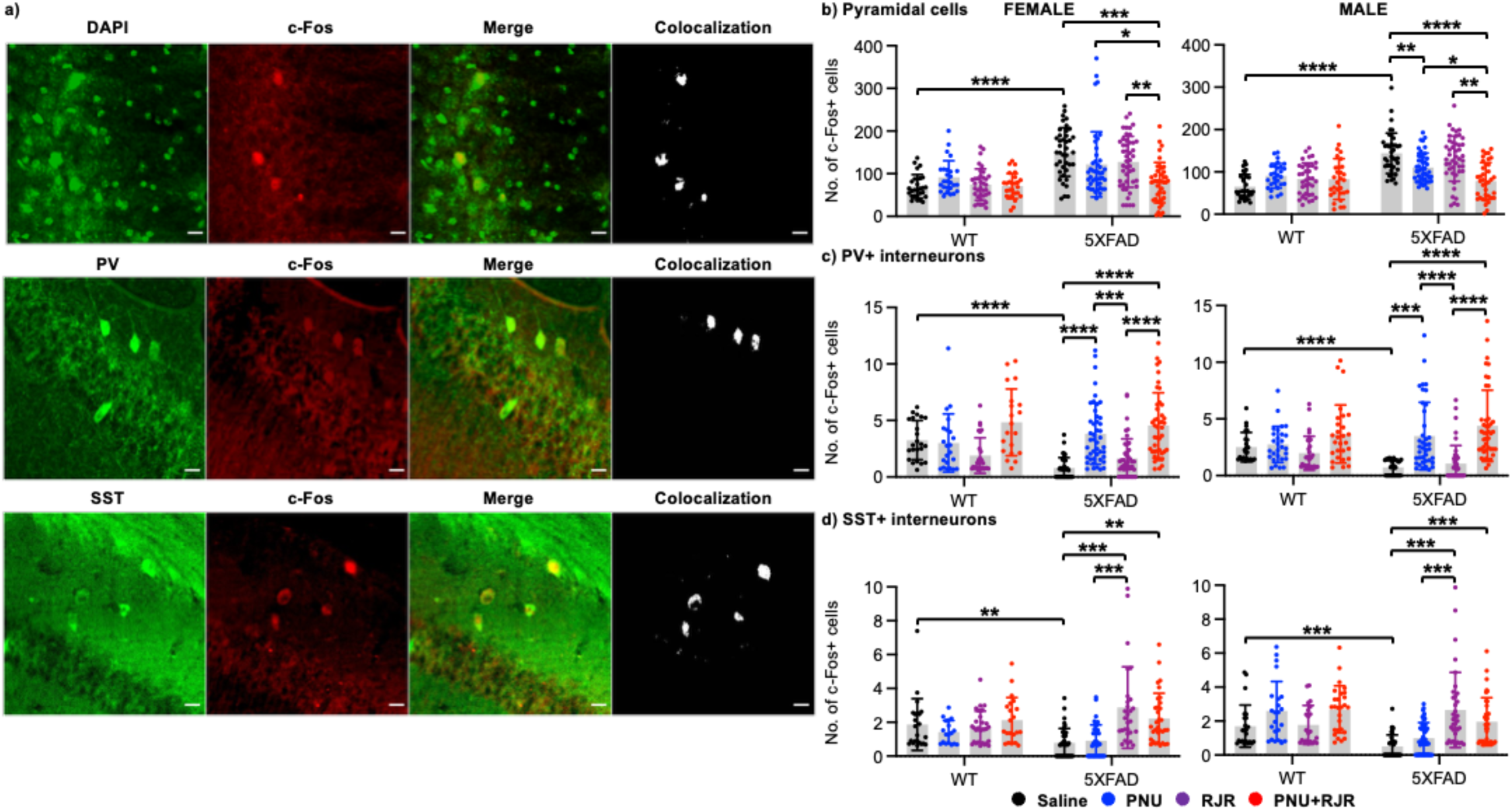
Co-stimulation of PV+ and SST+ interneurons by activating α7- and α4β2-nAChRs together is required to reverse hippocampal hyperexcitability in 5XFAD mice. **a)** Representative images of DAPI, PV, or SST (green), c-Fos (red), merge, and colocalization in the hippocampus. A scale bar represents 10 μm. **b)** Summary data of an average of the total number of c-Fos+ pyramidal neurons in each condition (n = number of sections [number of animals]; Female, WT + Saline = 28 [7], WT + PNU = 26 [7], WT + RJR = 36 [8], WT + PNU + RJR = 26 [6], 5XFAD + saline = 40 [9], 5XFAD + PNU = 44 [11], 5XFAD + RJR = 46 [10], and 5XFAD + PNU + RJR = 42 [10], Male, WT + Saline = 30 [6], WT + PNU = 28 [6], WT + RJR = 36 [7], WT + PNU + RJR = 30 [6], 5XFAD + saline = 40 [9], 5XFAD + PNU = 44 [11], 5XFAD + RJR = 45 [11], and 5XFAD + PNU + RJR = 37 [10]). **c)** Summary data of an average of the total number of c-Fos+ PV+ interneurons in each condition (n = number of sections [number of animals]; Female, WT + Saline = 23 [7], WT + PNU = 22 [7], WT + RJR = 28 [8], WT + PNU + RJR = 20 [6], 5XFAD + saline = 40 [9], 5XFAD + PNU = 45 [11], 5XFAD + RJR = 45 [10], and 5XFAD + PNU + RJR = 44 [10], Male, WT + Saline = 23 [6], WT + PNU = 28 [6], WT + RJR = 31 [7], WT + PNU + RJR = 28 [6], 5XFAD + saline = 35 [9], 5XFAD + PNU = 39 [11], 5XFAD + RJR = 46 [11], and 5XFAD + PNU + RJR = 44 [10]). **d)** Summary data of an average of the total number of c-Fos+ SST+ interneurons in each condition (n = number of sections [number of animals]; Female, WT + Saline = 23 [7], WT + PNU = 18 [7], WT + RJR = 30 [8], WT + PNU + RJR = 21 [6], 5XFAD + saline = 40 [9], 5XFAD + PNU = 43 [11], 5XFAD + RJR = 28 [10], and 5XFAD + PNU + RJR = 33 [10], Male, WT + Saline = 24 [6], WT + PNU = 23 [6], WT + RJR = 22 [7], WT + PNU + RJR = 27 [6], 5XFAD + saline = 39 [9], 5XFAD + PNU = 47 [11], 5XFAD + RJR = 35 [11], and 5XFAD + PNU + RJR = 36 [10]). \**p* < 0.05, \*\**p* < 0.01, \*\*\**p* < 0.001, and \*\*\*\**p* < 0.0001. Mixed-effects models / repeated-measures ANOVA, Tukey test.

As single receptor activation was unable to reverse hyperexcitation, we next treated PNU and RJR together and found that co-activation of α7- and α4β2-nAChRs was required to reduce hippocampal hyperexcitation in 5XFAD mice (Female; 5XFAD + PNU + RJR, 79.067 ± 47.185, *p* = 0.0001 and Male; 5XFAD + PNU + RJR, 78.509 ± 43.314, *p* < 0.0001) (**Fig. 3b, Fig. S1, and Table S7**). Importantly, agonist treatments had no major effect on the activity of hippocampal pyramidal cells in WT mice (**Fig. 3b, Fig. S1, and Table S7**). Notably, in the male 5XFAD hippocampus, PNU + RJR treatment had a stronger effect on neuronal hyperexcitation than PNU alone (*p* = 0.0175) (**Table S7**). This thus suggests that Aβ induces neuronal hyperactivation in hippocampal pyramidal neurons in 5XFAD mice, and co-stimulation of α7- and α4β2-nAChRs is required to reverse hippocampal hyperexcitability in 5XFAD mice.

We next examined whether hippocampal disinhibition was linked to hyperexcitation in 5XFAD mice. The number of in c-Fos+ PV+ cells was significantly decreased in the 5XFAD hippocampus (Female; WT + Saline, 3.256 ± 1.730 and 5XFAD + Saline, 0.788 ± 0.929, *p* < 0.0001, Male; WT + Saline, 2.513 ± 1.294 and 5XFAD + Saline, 0.733 ± 0.624, *p* < 0.0001) (**Fig. 3c, Fig. S2, and Table S8**). PNU treatment significantly increased the hippocampal PV+ cells’ activity in 5XFAD mice (Female; 5XFAD + PNU, 3.827 ± 2.703, *p* < 0.0001 and Male; 5XFAD + PNU, 3.487 ± 2.980, *p* = 0.0005) (**Fig. 3c, Fig. S2, and Table S8**). Conversely, RJR had no effect on PV+ cells’ activity in the 5XFAD hippocampus (Female; 5XFAD + RJR, 1.627 ± 1.735, *p* = 0.0735 and Male; 5XFAD + RJR, 1.067 ± 1.580, *p* = 0.8721) (**Fig. 3c, S2, and Table S8**). Co-activation of α7- and α4β2-nAChRs markedly elevated the activity of hippocampal PV+ cells to the same degree as PNU treatment itself (Female; 5XFAD + PNU + RJR, 4.554 ± 2.889, *p* < 0.0001 and Male; 5XFAD + PNU + RJR, 4.391 ± 3.143, *p* < 0.0001) (**Fig. 3c, Fig. S2, and Table S8**). Agonist treatments had no major effect on PV+ cells’ activity in the WT hippocampus (**Fig. 3c, Fig. S2, and Table S8**). This suggests that Aβ reduces the activity of hippocampal PV+ cells, but activation of α7-nAChRs by PNU selectively simulates these cells in 5XFAD mice, which is consistent with our findings in cultured neurons (**Fig. 2a and 2c**).

Finally, we examined the contribution of SST+ cells to hyperexcitability. The hippocampal SST+ cells’ activity was markedly reduced in 5XFAD mice (Female; WT + Saline, 1.868 ± 1.527 and 5XFAD + Saline, 0.781 ± 0.854, *p* = 0.0037, Male; WT + Saline, 1.699 ± 1.238 and 5XFAD + Saline, 0.511 ± 0.669, *p* =0.0001) (**Fig. 3d, Fig. S3, and Table S9**). Notably, PNU was unable to affect SST+ cells’ activity in the 5XFAD hippocampus (Female; 5XFAD + PNU, 0.911 ± 0.934, *p* = 0.6740 and Male; 5XFAD + PNU, 1.001 ± 0.903, *p* = 0.2145) (**Fig. 3d, Fig. S3, and Table S9**). However, RJR treatment markedly elevated the activity of hippocampal SST+ cells in 5XFAD mice (Female; 5XFAD + RJR, 2.876 ± 2.408, *p* = 0.0006 and Male; 5XFAD + RJR, 2.650 ± 2.216, *p* < 0.0001) (**Fig. 3d, Fig. S3, and Table S9**). Importantly, co-activation of α7- and α4β2-nAChRs significantly elevated the hippocampal SST+ cells’ activity to the same degree as RJR treatment itself in 5XFAD mice (Female; 5XFAD + PNU + RJR, 2.219 ± 1.498, *p* = 0.0002 and Male; 5XFAD + PNU + RJR, 1.967 ± 1.413, *p* = 0.0004) (**Fig. 3d, Fig. S3, and Table S9**). Notably, agonist treatments had no effect on hippocampal SST+ cells’ activity in WT mice (**Fig. 3d, Fig. S3, and Table S9**). This suggests that Aβ reduces the hippocampal SST+ cells’ activity, but stimulation of α4β2-nAChRs by RJR selectively activates these cells in 5XFAD mice, which is consistent with our findings in cultured neurons (**Fig. 2b and 2d**). These findings all together suggest that Aβ significantly reduces both PV+ and SST+ interneurons’ activity to induce hyperexcitation in pyramidal cells in 5XFAD hippocampus, and co-stimulation of PV+ and SST+ cells via activating α7- and α4β2-nAChRs together is required to reduce disinhibition-induced hippocampal hyperexcitability in 5XFAD mice.

### Deficits in hippocampal network activity in 5XFAD mice following fear learning

Neural network activity in the hippocampus plays important roles in memory processes (Seidenbecher et al. 2003; Narayanan et al. 2007; Markram et al. 1997; Colgin 2016; Duzel et al. 2010). Studies have shown that altered hippocampal network activity can impair learning and memory processes, which contributes to the development of AD (Filippini et al. 2009; Mondadori et al. 2006; Bateman et al. 2012; Reiman et al. 2012; Herrmann and Demiralp 2005). In people at high risk of developing AD, abnormal activation and/or deactivation of specific network activities during hippocampal memory processes are detected decades before the onset of clinical disease (Filippini et al. 2009; Mondadori et al. 2006; Bateman et al. 2012; Reiman et al. 2012; Herrmann and Demiralp 2005; Gallego-Rudolf et al. 2024b). This suggests that monitoring hippocampal network activity may be a useful tool for early detection and diagnosis of AD. Especially, hippocampal inhibitory interneurons regulate network oscillations of local field potentials (LFPs), which are thought to play a mechanistic role in various aspects of hippocampus-dependent memory processes (Seidenbecher et al. 2003; Narayanan et al. 2007; Markram et al. 1997; Colgin 2016; Duzel et al. 2010; Pelkey et al. 2017). In mice, it has been reported that theta (4-8 Hz) synchrony is increased between the hippocampal CA1 and lateral amygdala during the consolidation and reconsolidation of fear memory (Seidenbecher et al. 2003; Narayanan et al. 2007). Research also suggests that gamma rhythms (30-100 Hz) are two functionally distinct oscillatory activities, slow (30-50 Hz) and fast (50-100 Hz) gamma (Bieri et al. 2014; Griffiths et al. 2019). Slow gamma oscillations are likely involved in storing memory (Nakazawa et al. 2002; Steffenach et al. 2002; Treves and Rolls 1992). Fast gamma rhythms promote the transmission of current sensory information to the hippocampus during new memory encoding (Colgin 2016; Mably and Colgin 2018). It has been shown that spatial learning increases theta, slow gamma, and fast gamma powers in the hippocampal CA1 area (Rossato et al. 2019; Zheng et al. 2016; Trimper et al. 2014). We thus first recorded hippocampal LFPs before and following contextual fear conditioning (CFC) to determine if hippocampal network activity was disrupted in 5XFAD mice after fear learning. Prior to learning, basal LFPs were recorded for an hour (PRE) (**Fig. 4a**). Especially, the excitatory/inhibitory (E/I) balance within the hippocampus is critical for neural network stability and is precisely regulated during sleep to facilitate memory consolidation (Watson and Buzsaki 2015). Consolidation of hippocampus-dependent fear memory is known to occur between 1 and 3 hours after training (Kwapis and Wood 2014; Bourtchouladze et al. 1998). Therefore, test animals were returned to their home cage for 90 minutes after fear learning, and then we recorded hippocampal LFPs for additional one hour (POST) (**Fig. 4a**). We then compared hippocampal theta, slow gamma, and fast gamma powers between 5XFAD mice and WT littermates. We found that theta (Female; WT PRE, 1.000 ± 1.199 and WT POST, 1.696 ± 1.552, *p* = 0.0029, Male; WT PRE, 1.000 ± 1.270 and WT POST, 1.926 ± 2.849, *p* = 0.0001), slow gamma (Female; WT PRE, 1.000 ± 1.223 and WT POST, 1.624 ± 1.420, *p* = 0.0031, Male; WT PRE, 1.000 ± 1.124 and WT POST, 2.049 ± 2.413, *p* < 0.0001), and fast gamma powers (Female; WT PRE, 1.000 ± 1.086 and WT POST, 1.601 ± 1.372, *p* = 0.0009, Male; WT PRE, 1.000 ± 1.051 and WT POST, 2.467 ± 2.486, *p* < 0.0001) were significantly elevated following CFC in WT female and male mice (**Fig. 4e-g and Table S10**). Conversely, 5XFAD mice showed significant reduction in powers of theta (Female; 5XFAD PRE, 2.195 ± 6.108 and 5XFAD POST, 0.225 ± 0.210, *p* = 0.0015, Male; 5XFAD PRE, 2.168 ± 3.446 and 5XFAD POST, 0.539 ± 1.019, *p* < 0.0001), slow gamma (Female; 5XFAD PRE, 3.145 ± 7.767 and 5XFAD POST, 0.480 ± 0.342, *p* = 0.0007, Male; 5XFAD PRE, 2.095 ± 3.080 and 5XFAD POST, 0.690 ± 1.940, *p* < 0.0001), and fast gamma bands (Female; 5XFAD PRE, 3.059 ± 7.262 and 5XFAD POST, 0.493 ± 0.284, *p* = 0.0005, Male; 5XFAD PRE, 2.055 ± 2.724 and 5XFAD POST, 0.770 ± 0.855, *p* < 0.0001) (**Fig. 4e-g and Table S10**). Interestingly, we found significant higher basal powers in 5XFAD mice than WT animals in all frequency bands (**Fig. 4e-g and Table S10**), which is likely related to Aβ-induced early hippocampal hyperexcitation. These findings suggest that the learning-induced increase in hippocampal oscillations is impaired in 5XFAD mice.

**Figure 4.**
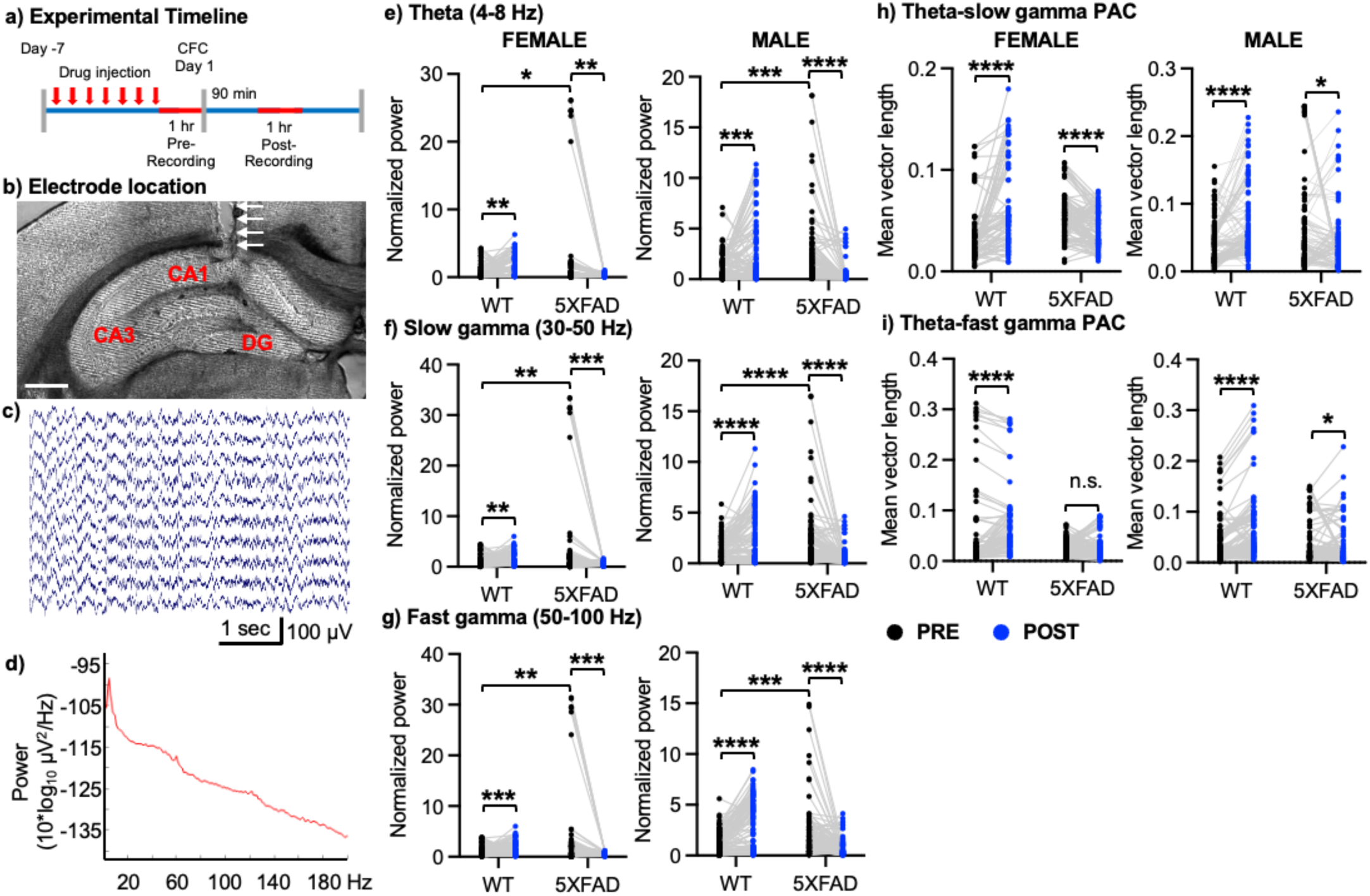
Deficits in hippocampal network activity in 5XFAD mice following fear learning. **a)** Experimental timeline. **b)** A representative image of a brain section after electrolytic lesioning indicates electrode placement (arrows) in the hippocampus CA1 area. A scale bar represents 1 mm. **c)** Representative section of LFP traces in CA1 of a freely behaving mouse. **d)** Representative power spectral density. Summary graphs of normalized powers of **e)** theta, **f)** slow gamma and **g)** fast gamma frequencies in female and male mice in each condition. Summary of **h)** Theta-slow gamma PAC and **i)** Theta-fast gamma PAC in each condition. n = number of channels [number of animals]; Female, WT = 74 [7] and 5XFAD = 101 [9], Male, WT =125 [12] and 5XFAD = 106 [10]). \**p* < 0.05, \*\**p* < 0.01, \*\*\**p* < 0.001, and \*\*\*\**p* < 0.0001. n.s. – not significant. Paired two-tailed student’s t-test was used to compare between PRE and POST data. Two-way ANOVA, Šídák’s test was used to compare between WT PRE and 5XFAD PRE data.

In healthy rodents and humans, the hippocampus exhibits prominent gamma oscillations that emerge at specific phases of theta oscillations known as theta-gamma coupling (Nakazono et al. 2018). The strength of theta-gamma coupling in the hippocampus is correlated with the task load of memory in humans (Axmacher et al. 2010). This coupling is impaired in hippocampal slices in AD model mice (Goutagny et al. 2013). Appropriate phase-amplitude coupling (PAC) and entrainment to hippocampal oscillations are thought to be essential for cognition and have been directly linked to memory performance in humans (Axmacher et al. 2010; Daume et al. 2024; Lega et al. 2016). We thus calculated the mean vector length (MVL) as an entrainment strength. We found that the MVL in both theta-slow gamma and theta-fast gamma was significantly higher following learning than before learning in WT female and male mice (Theta-slow gamma, Female; WT PRE, 0.036 ± 0.027 and WT POST, 0.067 ± 0.044, *p* < 0.0001, Male; WT PRE, 0.041 ± 0.032 and WT POST, 0.070 ± 0.053, *p* < 0.0001. Theta-fast gamma, Female; WT PRE, 0.059 ± 0.087 and WT POST, 0.076 ± 0.070, *p* = 0.0004, Male; WT PRE, 0.036 ± 0.042 and WT POST, 0.061 ± 0.061, *p* < 0.0001) (**Fig. 4h-I and Table S11**). In contrast, the theta-slow gamma MVL in 5XFAD mice was significantly lower following learning when compared to the basal condition (Female; 5XFAD PRE, 0.055 ± 0.021 and 5XFAD POST, 0.042 ± 0.018, *p* < 0.0001, Male; 5XFAD PRE, 0.060 ± 0.067 and 5XFAD POST, 0.043 ± 0.047, *p* = 0.0157) (**Fig. 4h-I and Table S11**). Moreover, fear learning was unable to elevate 5XFAD animals’ theta-fast gamma PAC in the hippocampus (Female; 5XFAD PRE, 0.028 ± 0.015 and 5XFAD POST, 0.028 ± 0.022, *p* = 0.9556, Male; 5XFAD PRE, 0.041 ± 0.046 and 5XFAD POST, 0.030 ± 0.039, *p* = 0.0180) (**Fig. 4h-I and Table S11**). This data suggest that fear learning strengthens PAC entrainment in WT mice, consistent with the previous report (Radiske et al. 2020). However, fear leaning is unable to increase theta-gamma coupling in 5XFAD mice. These findings thus show deficits in hippocampal network activity in 5XFAD mice following fear learning.

### Co-activation of α7- and α4β2-nAChRs is required to restore normal hippocampal network activity in 5XFAD mice

We next compared hippocampal oscillatory activity following fear learning between WT and 5XFAD mice. When compared to WT mice, 5XFAD mice exhibited a significant decrease in the powers of theta (Female; WT, 1.000 ± 0.915 and 5XFAD, 0.151 ± 0.148, *p* < 0.0001, Male; WT, 1.000 ± 1.479 and 5XFAD, 0.280 ± 0.529, *p* < 0.0001), slow gamma (Female; WT, 1.000 ± 1.874 and 5XFAD, 0.268 ± 10.167, *p* = 0.0015, Male; WT, 1.000 ± 1.079 and 5XFAD, 0.304 ± 0.414, *p* = 0.0007), and fast gamma bands (Female; WT, 1.000 ± 0.853 and 5XFAD, 0.297 ± 0.162, *p* = 0.0024, Male; WT, 1.000 ± 1.008 and 5XFAD, 0.312 ± 0.347, *p* < 0.0001) (**Fig. 5a-c and Table S12-S14**). This indicates impaired fear memory processing in 5XFAD mice. Acetylcholine modulates hippocampal oscillations to control memory processes (Wilson and Fadel 2017; Haam and Yakel 2017; Headley and Pare 2017). We thus examined if stimulating α7- and α4β2-nAChRs together was required to reverse hippocampal network deficits in 5XFAD mice after fear conditioning. We found that activation of single receptors by themselves using PNU or RJR had no effect on the powers of theta (Female; 5XFAD + PNU, 0.138 ± 0.234, *p* = 0.9997, and 5XFAD + RJR, 0.127 ± 0.174, *p* = 0.9984, Male; 5XFAD + PNU, 0.123 ± 0.137, *p* = 0.8638, and 5XFAD + RJR, 0.090 ± 0.107, *p* = 0.7720), slow gamma (Female; 5XFAD + PNU, 0.305 ± 0.419, *p* = 0.9988, and 5XFAD + RJR, 0.214 ± 0.297, *p* = 0.8501, Male; 5XFAD + PNU, 0.272 ± 0.157, *p* = 0.9995, and 5XFAD + RJR, 0.142 ± 0.122, *p* = 0.9405), and fast gamma bands (Female; 5XFAD + PNU, 0.269 ± 0.392, *p* = 0.9995, and 5XFAD + RJR, 0.198 ± 0.240, *p* = 0.9849, Male; 5XFAD + PNU, 0.263 ± 0.195, *p* = 0.9962, and 5XFAD + RJR, 0.142 ± 0.139, *p* = 0.8634) (**Fig. 5a-c and Table S12-S14**). This data show that stimulation of either α7- or α4β2-nAChRs is insufficient to improve hippocampal network activity in 5XFAD mice. However, co-stimulation of α7- and α4β2-nAChRs significantly elevated the powers of theta (Female; 5XFAD + PNU+RJR, 0.518 ± 0.834, *p* = 0.0057, Male; 5XFAD + PNU+RJR, 0.684 ± 0.986, *p* = 0.0460), slow gamma (Female; 5XFAD + PNU + RJR, 1.058 ± 1.633, *p* = 0.0006, Male; 5XFAD + PNU + RJR, 1.300 ± 2.375, *p* = 0.0001), and fast gamma bands (Female; 5XFAD + PNU + RJR, 1.253 ± 1.919, *p* < 0.0001, Male; 5XFAD + PNU + RJR, 1.261 ± 1.763, *p* < 0.0001) (**Fig. 5a-c and Table S12-S14**). Interestingly, agonist treatment significantly reduced theta power only in WT mice (Female; WT + PNU, 0.490 ± 0.950, *p* = 0.0077, WT + RJR, 0.566 ± 0.600, *p* = 0.0138, WT + PNU + RJR, 0.457 ± 1.209, *p* < 0.0001, Male; WT + PNU, 0.422 ± 0.365, *p* = 0.0031, WT + RJR, 0.489 ± 0.256, *p* = 0.0173, WT + PNU + RJR, 0.609 ± 1.648, *p* = 0.0382) while drugs had no major effect on slow and fast gamma powers in WT animals (**Fig. 5a-c and Table S12-S14**). These findings suggest that a memory consolidation-mediated increase in hippocampal oscillations is disrupted in 5XFAD mice, which is reversed only by co-activation of α7-and α4β2-nAChRs.

**Figure 5.**
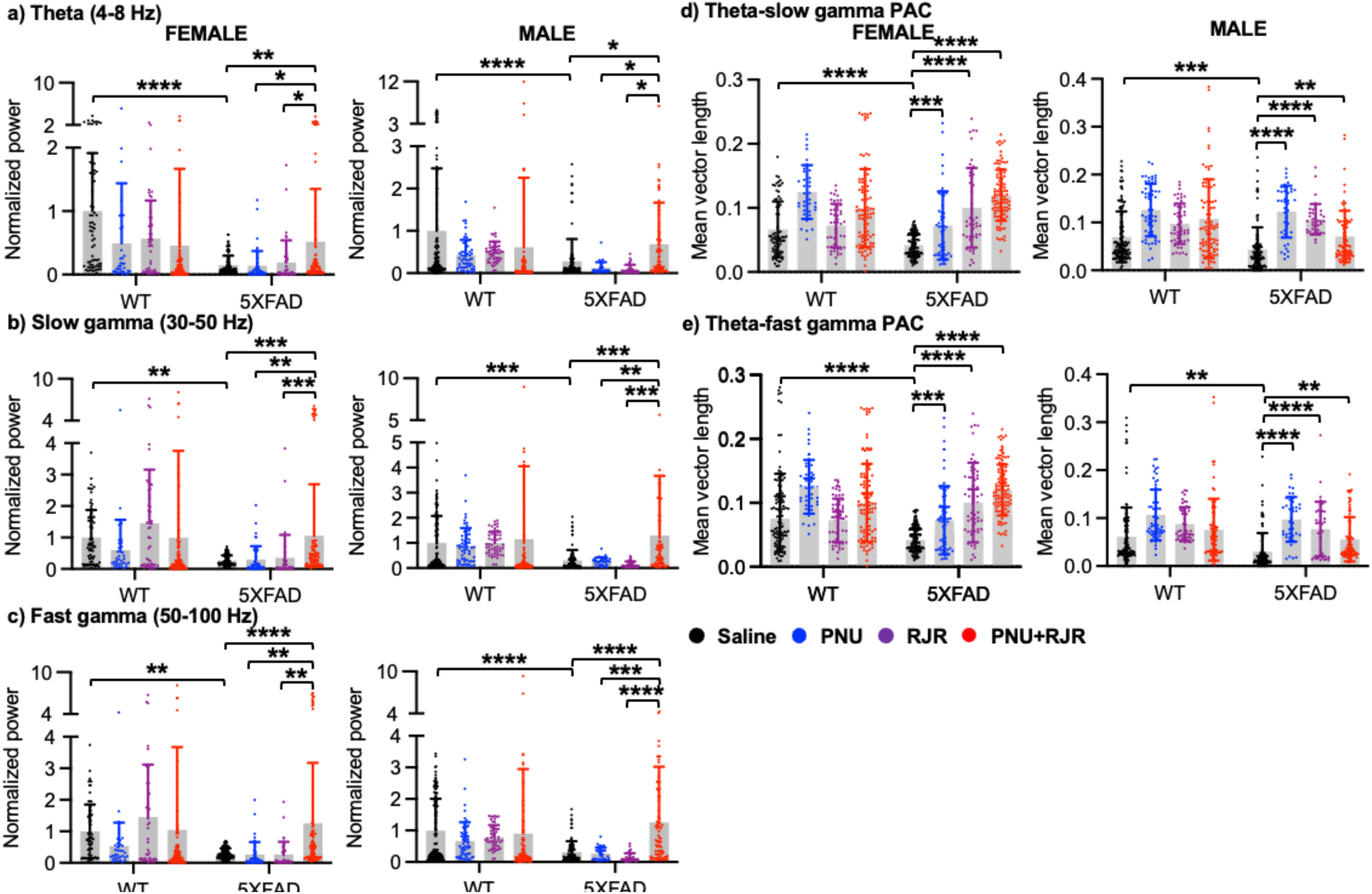
Co-activation of α7- and α4β2-nAChRs is required to restore normal hippocampal network activity in 5XFAD mice. Summary graphs of normalized powers of **a)** theta, **b)** slow gamma and **c)** fast gamma frequencies in female and male mice in each condition. Summary of **d)** Theta-slow gamma PAC and **e)** Theta-fast gamma PAC in each condition. n = number of channels [number of animals]; Female, WT + Saline = 74 [7], WT + PNU = 35 [4], WT + RJR = 48 [4], WT + PNU + RJR = 97 [9], 5XFAD + saline = 101 [9], 5XFAD + PNU = 51 [4], 5XFAD + RJR = 41 [4], and 5XFAD + PNU + RJR = 93 [8], Male, WT + Saline = 125 [12], WT + PNU = 57 [5], WT + RJR = 51 [4], WT + PNU + RJR = 86 [8], 5XFAD + saline = 106 [10], 5XFAD + PNU = 36 [4], 5XFAD + RJR = 38 [4], and 5XFAD + PNU + RJR = 79 [7]. \**p* < 0.05, \*\**p* < 0.01, \*\*\**p* < 0.001, and \*\*\*\**p* < 0.0001. Mixed-effects models / repeated-measures ANOVA, Tukey test.

We next analyzed the strength of theta-gamma coupling in the hippocampus. We found that the MVL in both theta-slow gamma and theta-fast gamma was significantly lower in 5XFAD mice following learning than WT animals (Theta-slow gamma, Female; WT, 0.067 ± 0.044 and 5XFAD, 0.042 ± 0.018, *p* = 0.0004, Male; WT, 0.070 ± 0.053 and 5XFAD, 0.043 ± 0.047, *p* = 0.0003. Theta-fast gamma, Female; WT, 0.076 ± 0.070 and 5XFAD, 0.028 ± 0.022, *p* < 0.0001, Male; WT, 0.061 ± 0.061 and 5XFAD, 0.030 ± 0.039, *p* < 0.0001) (**Fig. 5d-e and Table S15 and S16**). We found that activation of single receptors or two receptors together was sufficient to increase MVL in 5XFAD mice (Theta-slow gamma, Female; 5XFAD + PNU, 0.072 ± 0.053, *p* = 0.0005, 5XFAD + RJR, 0.100 ± 0.062, *p* < 0.0001, and 5XFAD + PNU + RJR, 0.120 ± 0.040, *p* < 0.0001, Male; 5XFAD + PNU, 0.122 ± 0.054, *p* < 0.0001, 5XFAD + RJR, 0.107 ± 0.032, *p* < 0.0001, and 5XFAD + PNU + RJR, 0.071 ± 0.053, *p* = 0.0055. Theta-fast gamma, Female; 5XFAD + PNU, 0.055 ± 0.039, *p* = 0.0008, 5XFAD + RJR, 0.086 ± 0.036, *p* < 0.0001, and 5XFAD + PNU + RJR, 0.078 ± 0.041, *p* < 0.0001, Male; 5XFAD + PNU, 0.097 ± 0.047, *p* < 0.0001, 5XFAD + RJR, 0.077 ± 0.057, *p* < 0.0001, and 5XFAD + PNU + RJR, 0.056 ± 0.046, *p* = 0.0052) (**Fig. 5d-e and Table S15 and S16**). This data show that activation of α7- and/or α4β2-nAChRs is sufficient to strengthen fear learning-related PAC entrainment in 5XFAD mice. In sum, 5XFAD mice showed significant deficits in hippocampal network activity following fear conditioning. While stimulation of α7-and/or α4β2-nAChRs is sufficient to increase PAC in 5XFAD mice, co-activation of α7- and α4β2-nAChRs, perhaps stimulating PV+ and SST+ cells together, is essential to restore normal powers of AD-related hippocampal oscillations following fear learning.

### Co-activation of α7- and α4β2-nAChRs is required to improve hippocampus-dependent memory in 5XFAD mice

A novel object location (NOL) test is known to assess hippocampus-dependent spatial recognition memory by measuring an animal’s ability to detect a familiar object’s displacement to a novel location (Denninger et al. 2018). We thus conducted NOL as described previously (Denninger et al. 2018) with 5-month-old 5XFAD and WT female and male mice. The preference index (PI) for the relocated object was compared between the training and test sessions. The PI for the relocated object was significantly higher during the test session than during the training session in saline-treated WT control mice (Female, WT + Saline training, 0.349 ± 0.198 and WT + Saline test, 0.530 ± 0.149, *p* < 0.0072, and Male, WT + Saline training, 0.404 ± 0.184 and WT+ Saline test, 0.575 ± 0.256, *p* < 0.0114) (**Fig. 6a and Table S17**). This significant increase in preference for the relocated object indicates that WT mice recognized the change in object location and preferentially investigated the object in its novel location, consistent with intact spatial memory. In contrast, 5XFAD mice did not show a significant increase in PI during the test session compared with the training session (Female, 5XFAD + Saline training, 0.585 ± 0.078 and 5XFAD + Saline test, 0.552 ± 0.191, *p* = 0.0823, and Male, 5XFAD + Saline training, 0.506 ± 0.115 and 5XFAD + Saline test, 0.589 ± 0.191, *p* = 0.4290) (**Fig. 6a and Table S17**). Thus, 5XFAD mice failed to exhibit a significant preference for the relocated object, suggesting impaired spatial recognition memory, consistent with the previous reports (Terstege et al. 2023; Zhang et al. 2023; Creighton et al. 2019). Given that co-activation of α7- and α4β2-nAChRs is required to reverse fear learning-associated hippocampal network activity (**Fig. 5**), we determined the effects of activating these receptors on spatial recognition memory in 5XFAD mice. We intraperitoneally injected PNU or RJR into 5XFAD and WT mice to see if activation of either α7- or α4β2-nAChRs reversed impaired spatial recognition memory in AD model mice. Single administration of either PNU or RJR failed to increase the PI during the test session in 5XFAD mice (Female, 5XFAD + PNU training, 0.567 ± 0.108 and 5XFAD + PNU test, 0.546 ± 0.223, *p* = 0.7310, and Male, 5XFAD + PNU training, 0.438 ± 0.245 and 5XFAD + PNU test, 0.283 ± 0.232, *p* = 0.1186. Female, 5XFAD + RJR training, 0.523 ± 0.132 and 5XFAD + RJR test, 0.577 ± 0.188, *p* = 0.5159, and Male, 5XFAD + RJR training, 0.552 ± 0.114 and 5XFAD + RJR test, 0.582 ± 0.171, *p* = 0.7547) (**Fig. 6a and Table S17**). This shows that stimulation of either α7- or α4β2-nAChRs is insufficient to improve spatial memory in 5XFAD mice. We next intraperitoneally injected PNU and RJR together into 5XFAD and WT littermates and examined whether co-activation of α7- and α4β2-nAChRs can increase the PI during the test session in 5XFAD mice. Notably, co-activation of α7- and α4β2-nAChRs significantly enhanced spatial recognition memory in 5XFAD male and female mice (Female, 5XFAD + PUN + RJR training, 0.393 ± 0.142 and 5XFAD + PUN + RJR test, 0.698 ± 0.137, *p* = 0.0022, and Male, 5XFAD + PUN + RJR training, 0.517 ± 0.109 and 5XFAD + PUN + RJR test, 0.731 ± 0.018, *p* = 0.0093) (**Fig. 6a and Table S17**). As this NOL test was used to assess short-term memory (Vogel-Ciernia and Wood 2014), we carried out contextual fear conditioning (CFC) to measure hippocampus-dependent long-term memory (Phillips and LeDoux 1992). Conditioned fear responses have been demonstrated to be impaired in patients with mild to moderate AD (Hamann et al. 2002; Hoefer et al. 2008). Fear memory loss in 5XFAD mice starts at the age of 5 months (Kimura and Ohno 2009). We thus carried out CFC as described previously (Kim et al. 2016; Shou et al. 2018) with 5-month-old 5XFAD and WT female and male mice. In consistent with the previous finding (Kimura and Ohno 2009), 5XFAD female and male mice intraperitoneally given saline displayed significantly reduced freezing in contextual fear conditioning compared to WT controls, indicating impaired hippocampus-dependent fear memory (Female, WT + Saline, 36.785 ± 10.866% and 5XFAD + Saline, 14.188 ± 12.202%, *p* < 0.0001, and Male, WT + Saline, 38.938 ± 13.066% and 5XFAD + Saline, 16.177 ± 8.761%, *p* < 0.0001) (**Fig. 6b and Table S18**). We next injected PNU or RJR to examine the effects of stimulating nAChRs on contextual fear memory in 5XFAD mice. Neither PNU nor RJR single injection reversed fear memory loss in 5XFAD mice (Female, 5XFAD + PNU, 15.751 ± 10.332%, *p* > 0.9999, and 5XFAD + RJR, 14.836 ± 10.400%, *p* > 0.9999, and Male, 5XFAD + PNU, 15.558 ± 6.577%, *p* > 0.9999, and 5XFAD + RJR, 14.648 ± 6.018%, *p* > 0.9999) (**Fig. 6b and Table S18**). This shows that stimulation of either α7- or α4β2-nAChRs is insufficient to improve fear memory in 5XFAD mice. Finally, we found that co-activation of α7- and α4β2-nAChRs significantly enhanced contextual fear memory in 5XFAD male and female mice (Female, 5XFAD + PNU + RJR, 43.664 ± 13.580%, *p* < 0.0001, and Male, 5XFAD + PNU + RJR, 43.434 ± 21.002%, *p* = 0.0101) (**Fig. 6b and Table S18**). Notably, normal spatial and fear memory was found in WT mice in the presence of agonists (**Fig. 6a-b and Table S17-18**). These findings suggest that co-activation of α7- and α4β2-nAChRs (potentially stimulates hippocampal PV+ and SST+ cells together) is required to improve hippocampus-dependent spatial and fear memory in 5XFAD mice.

**Figure 6.**
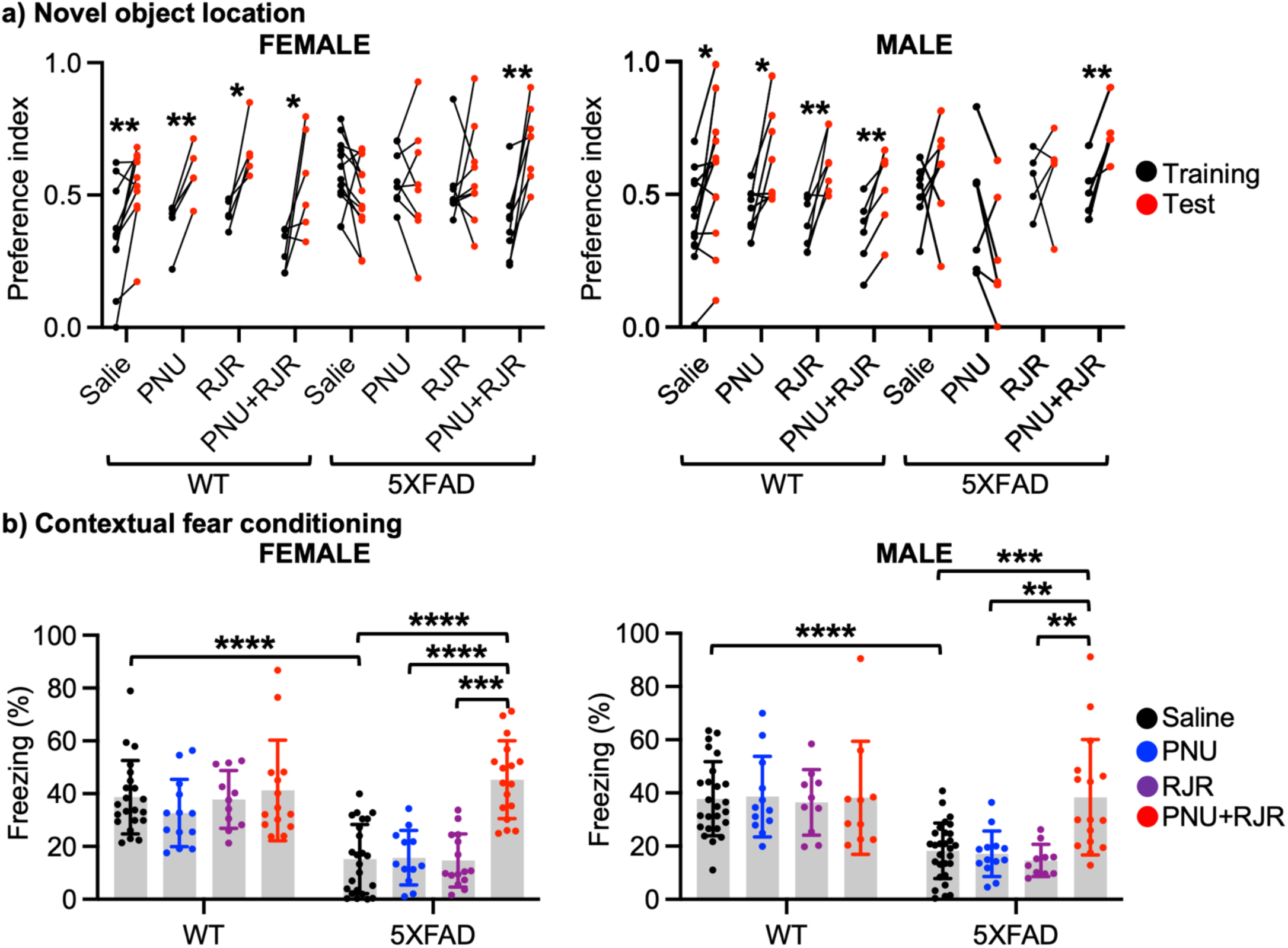
Co-activation of α7- and α4β2-nAChRs is required to improve hippocampus-dependent memory in 5XFAD mice. **a)** Summary data of spatial recognition memory of WT and 5XFAD female (n = number of animals, WT + Saline = 10, WT + PNU = 5, WT + RJR = 5, WT + PNU + RJR = 6, 5XFAD + Saline = 12, 5XFAD + PNU = 8, 5XFAD + RJR = 9, 5XFAD + PNU+RJR = 8) and male mice (n = number of animals, WT + Saline = 12, WT + PNU = 8, WT + RJR = 7, WT + PNU + RJR = 6, 5XFAD + Saline = 7, 5XFAD + PNU = 6, 5XFAD + RJR = 5, 5XFAD + PNU + RJR = 5) in each condition. \**p* < 0.05 and \*\**p* < 0.01, Paired two-tailed Student’s t-test. **b)** Summary data of freezing behavior of WT and 5XFAD female (n = number of animals, WT + Saline = 21, WT + PNU = 13, WT + RJR = 11, WT + PNU+RJR = 14, 5XFAD + Saline = 23, 5XFAD + PNU = 12, 5XFAD + RJR = 15, 5XFAD + PNU + RJR = 16) and male mice (n = number of animals, WT + Saline = 24, WT + PNU = 12, WT + RJR = 9, WT + PNU + RJR = 9, 5XFAD + Saline = 26, 5XFAD + PNU = 12, 5XFAD + RJR = 10, 5XFAD + PNU + RJR = 13) in each condition. \**p* < 0.05 and \*\*\*\**p* < 0.0001, Two-way ANOVA, Tukey test.

### Co-activation of α7- and α4β2-nAChRs is required to reduce the Aβ buildup in the 5XFAD hippocampus

Amyloid plaques are aggregation of Aβ in the extracellular space of the brain, central to the AD pathology. It has been shown that *in vivo* extracellular Aβ levels are dynamically and directly influenced by neuronal activity (Cirrito et al. 2005). Importantly, neuronal hyperexcitability increases Aβ secretion and accumulation (Cirrito et al. 2005; Yamamoto et al. 2015). This suggests that Aβ is a major trigger of hyperexcitability, which in turn establishes a vicious cycle of AD progress. Therefore, it is possible that Aβ-induced hippocampal hyperexcitation promotes the *in vivo* rapid growth of amyloid plaques, which can be prevented by reducing neuronal activity through enhancing hippocampal inhibition. To test this idea, we examined if enhancing PV+ and SST+ activity together by co-stimulating α7- and α4β2-nAChRs was required to reduce the Aβ buildup in the 5XFAD hippocampus. Notably, longitudinal *in vivo* imaging studies in AD pathology model mice have demonstrated that amyloid plaques form and grow quickly over the course of a day and one week (Yan et al. 2009; Meyer-Luehmann et al. 2008). We thus intraperitoneally injected PNU and RJR on their own or together into 5-month-old 5XFAD mice for 7 days. Saline was given to mice as a control. We then visualized amyloid plaques and measured the total number of amyloid plaques in the hippocampus. We found that stimulation of each receptor had no effect on amyloid pathology in the hippocampus of both female and male 5XFAD mice when compared to the saline-injected controls (Female, Saline, 50.62 ± 13.21, PNU, 49.71 ± 11.17, *p* = 0.9978, and RJR, 47.71 ± 10.28, *p* = 0.9359, and Male, Saline, 38.95 ± 13.53, PNU, 38.66 ± 8.85, *p* > 0.9999, and RJR, 47.88 ± 5.97, *p* = 0.2469) (**Fig. 7 and Table S19**). However, we revealed that co-stimulation of α7- and α4β2-nAChRs significantly reduced the total number of amyloid plaques in the hippocampus of both female and male 5XFAD mice when compared to the saline-injected controls (Female, PNU + RJR, 32.19 ± 7.19, *p* = 0.0013, and Male, PNU+RJR, 24.82 ± 9.89, *p* = 0.0042) (**Fig. 7 and Table S19**). This data show that co-activation of α7- and α4β2-nAChRs is required to reduce the rapid growth of amyloid plaques in 5XFAD mice.

**Figure 7.**
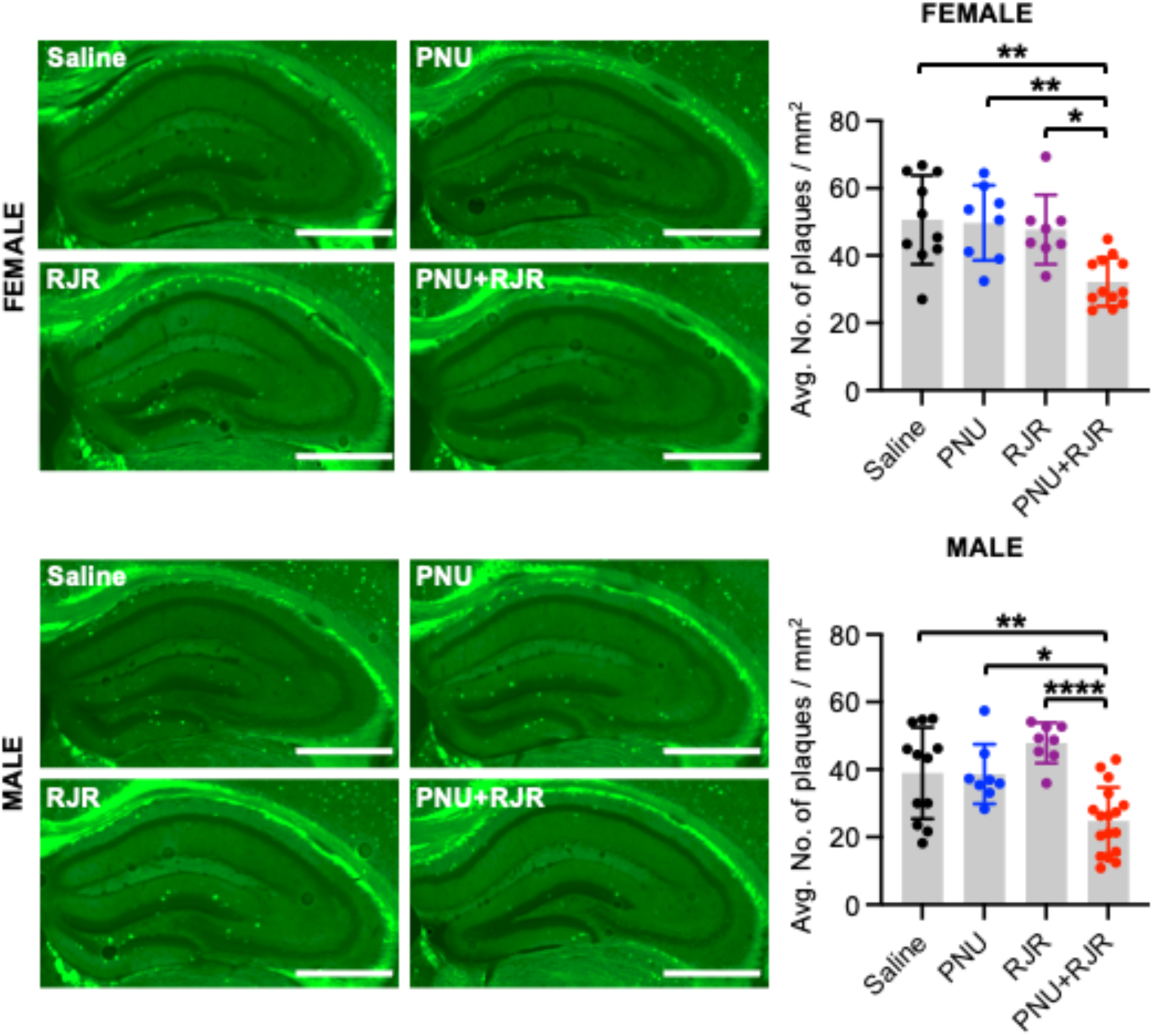
Co-activation of α7- and α4β2-nAChRs prevents the *in vivo* growth of amyloid plaques. Representative images of Thioflavin S-positive amyloid plaques (Green puncta) in the hippocampus in 5-month-old 5XFAD female (n = number of animals, Saline = 10, PNU = 8, RJR = 8, and PNU + RJR = 12) and male (n = number of animals, Saline = 12, PNU = 8, RJR = 8, and PNU + RJR = 17) mice in each condition. A scale bar represents 1 mm. Summary graphs of the total number of amyloid plaques per mm^2^ in each condition. \**p* < 0.05, \*\**p* < 0.01, and \*\*\*\**p* < 0.0001, One-way ANOVA, Tukey test.

## Discussion

A key role of toxic Aβ in AD (Cline et al. 2018; Guo et al. 2021; Jack et al. 2018) is supported by the success of recent clinical studies on anti-Aβ treatments resulting in the U.S. Food and Drug Administration (FDA) approval (Budd Haeberlein et al. 2022; van Dyck et al. 2023). However, their limited efficacy and significant side effects (Cadiz et al. 2024; Ebell et al. 2024; Fedele 2023; Kurkinen 2023; Loomis et al. 2024; Solopova et al. 2023), along with Aβ’s known physiological roles (Fedele 2023), suggest that targeting downstream effects of Aβ toxicity may be a safer and more effective strategy.

A large body of research has suggested that Aβ-induced inhibitory dysfunction in the hippocampus can be a crucial trigger for disruptions in neural oscillations and consequent cognitive function in AD (Verret et al. 2012; Park et al. 2020; Iaccarino et al. 2016; Chung et al. 2020; Kurudenkandy et al. 2014; Villette et al. 2010; Martinez-Losa et al. 2018; Gallego-Rudolf et al. 2024b). However, the molecular mechanisms underlying Aβ-induced hippocampal interneuron dysfunctions in AD have not been fully understood. Cholinergic dysfunction may be a prime suspect for Aβ-induced inhibitory dysfunction because cholinergic signaling has greater effects on hippocampal inhibitory cells than excitatory neurons (Ji et al. 2001; Frazier, Rollins, et al. 1998; Hefft et al. 1999; Jones and Yakel 1997; McQuiston and Madison 1999; Ji and Dani 2000), and there are diverse lines of evidence that molecular interactions between Aβ and nAChRs impair receptor functions in the early stage of AD (Jurgensen and Ferreira 2010; Lambert et al. 1998; McLean et al. 1999; Auld et al. 1998). This suggests that Aβ inhibits nAChR activities in inhibitory interneurons, which reduces hippocampal inhibition and oscillations to trigger cognitive decline in the early stage of AD. However, hippocampal inhibitory cells are highly diverse, and distinct interneuron subtypes differentially contributes to hippocampal oscillations and memory formation (Pelkey et al. 2017). In addition, various neuronal nAChRs have been found in the human brain (Albuquerque et al. 2009; Lindstrom 2003; Clementi et al. 2000). Distinct nAChR subtypes differentially affect hippocampal inhibition and oscillations (Lu et al. 2013; Alkondon and Albuquerque 2001; Griguoli and Cherubini 2012; Song et al. 2005; Gu et al. 2020). Therefore, it is challenging to elucidate the nAChR-mediated mechanisms by which Aβ affects hippocampal neural network and memory processing in AD at the interneuron subtype levels. Our published works have discovered that Aβ selectively binds to two of the three major hippocampal nAChR subtypes, α7- and α4β2-nAChRs, but not α3β4-nAChRs, and inhibits these two receptors in cultured hippocampal inhibitory interneurons to decrease their activity, leading to hyperexcitation in excitatory neurons (Roberts et al. 2021; Sun et al. 2019). We have also revealed that co-activation of α7-and α4β2-nAChRs is required to reverse the Aβ-induced adverse effects in hippocampal neurons (Roberts et al. 2021; Sun et al. 2019). In the current study, we further identify that α7-, α4β2-, and α3β4-nAChRs regulate the cholinergic activity mainly in PV+, SST+, and excitatory cells, respectively. Additionally, we discover that co-activation of α7- and α4β2-nAChRs is necessary to reverse hippocampal hyperexcitability, network dysfunction, and spatial a d fear memory loss and reduce amyloid pathology in 5XFAD mice. In sum, our study provides the mechanistic link between hippocampal disinhibition-induced hyperexcitability via altered cholinergic activity, network dysfunction, pathology, and memory loss in AD.

Based on our findings, we propose the following model. α7-, α4β2-, and α3β4-nAChRs predominantly control the nicotinic cholinergic signaling in PV+ interneurons that provide somatic inhibition, SST+ interneurons that provides dendritic inhibition, and pyramidal excitatory neurons, respectively, in the hippocampus, which is supported by the previous findings (**Fig. 8a**) (Alkondon et al. 1998; Alkondon and Albuquerque 2001; Alkondon et al. 2003; Buhler and Dunwiddie 2001; 2002; Frazier, Rollins, et al. 1998; Frazier, Buhler, et al. 1998; Jones and Yakel 1997; Lukas et al. 1999; McClure-Begley et al. 2009; McQuiston and Madison 1999; Sharma and Vijayaraghavan 2003; Sudweeks and Yakel 2000; Gu et al. 2020). These distinct cholinergic regulations of inhibitory activity contribute to inhibition onto pyramidal cells, which controls hippocampal network activity and memory processes (**Fig. 8a**). Aβ selectively inhibits α7- and α4β2-nAChRs in PV+ and SST+ cells, respectively, but not α3β4-nAChRs in excitatory neurons, which reduces inhibition to induce hyperexcitability and network dysfunction in the hippocampus, leading to amyloid pathology and memory loss in AD (**Fig. 8b**). Stimulation of PV+ or SST+ cells by the activation of each nAChR subtype-specific agonists, PNU-282987 (PNU), an α7 agonist, or RJR-2403 Oxalate (RJR), an α4β2 agonist, is insufficient to reverse the adverse effects of Aβ (**Fig. 8c-d**). Therefore, co-stimulation of hippocampal PV+ and SST+ inhibitory interneurons via activation of α7- and α4β2-nAChRs together is required to be protective against AD (**Fig. 8e**). Given that cholinergic deficiency is associated with AD, strategies aiming to restore normal cholinergic function have been developed as therapeutic drugs for AD, including several acetylcholinesterase inhibitors (Frolich 2002). Therefore, all cholinergic synapses can be intensified by these drugs. However, some subtypes of nAChRs that may not be affected by Aβ (e.g. α3β4) can also be activated when attempts are made to use such drugs in the treatment of AD, resulting in unexpected side effects. Moreover, acetylcholinesterase inhibitors have been shown to be only partially effective in reversing experimental deficits in hippocampal-dependent memory in rodents (Yuede et al. 2007). Consistently, we have revealed a non-selective cholinergic agonist exacerbates Aβ adverse effects on hippocampal neurons (Sun et al. 2019; Roberts et al. 2021). Furthermore, overstimulation of the M1 type metabotropic AChRs adversely affect neuronal activity in the prefrontal cortex, related to working memory (Vijayraghavan et al. 2018). Therefore, acetylcholinesterase inhibitors are not very effective in slowing AD progression due to non-selective stimulation of acetylcholine receptors (Raina et al. 2008; Sun et al. 2008). There are also discrepancies involving the use of nicotine treatment to stimulate nAChRs to alter cognitive function. For example, nicotine agonists have been found to improve performance in a variety of memory tasks in rodents and non-human primate studies (Levin and Simon 1998), while several other studies have failed to find significant enhancement of learning and memory by nicotine treatment (Vicens et al. 2003). Nonetheless, nAChR agonists have consistently suggested promising approaches in the treatment of AD (Buccafusco 2004). However, clinical trials thus far have been challenged by adverse effects or minimal improvement (Hoskin et al. 2019). Additionally, some studies demonstrate that stimulating one type of nAChRs by using a subtype specific agonist enhances cognitive performance, but other studies find no beneficial effect. For instance, selective α7 nAChR agonists have been reported to improve cognition in a variety of animal models (Bitner et al. 2007; Cincotta et al. 2008; Redrobe et al. 2009), while another study has found almost no beneficial effect on learning and memory in mice (Vicens et al. 2011). An α4β2-nAChR agonist alone can improve working memory only in young rats but not older animals (Levin and Christopher 2002). It is thus not yet clear whether single activation of specific nAChR subtypes provide optimal efficacy in AD (Buccafusco 2004). Research also shows that co-activation of α7- and α4β2-nAChRs has substantially greater neuroprotective effects against a ketamine-based model of schizophrenia-like cognitive impairments in rats, when compared to single receptor stimulation (Potasiewicz et al. 2018). Moreover, pharmacological inhibition of each α7- or α4β2-nAChR subtype has no effect on working memory in nonhuman primates (Upright and Baxter 2021). Additionally, both α7-nAChR agonists and positive allosteric modulators have largely failed to demonstrate efficacy in placebo-controlled trials in schizophrenia (Kantrowitz et al. 2020). These studies suggest that proper cholinergic modulation in hippocampal functions requires both α7- and α4β2-nAChRs. Our published works (Roberts et al. 2021; Sun et al. 2019) and current study further support this idea. Therefore, the idea that selective co-activation of nAChRs in the hippocampus can reverse Aβ effects on AD pathology is a new concept, which may lead to innovative and novel therapeutic strategies. In fact, several subtype-specific agonists have been developed for clinical trials, but co-activation of nAChRs has not been applied for clinical trials yet (Buccafusco 2004). The current work thus suggests activation of α7- and α4β2-nAChRs as an innovative and selective way to stimulate hippocampal PV+ and SST+ cells in the hippocampal local circuits, which has potentials for a novel therapeutic strategy in AD.

**Figure 8.**
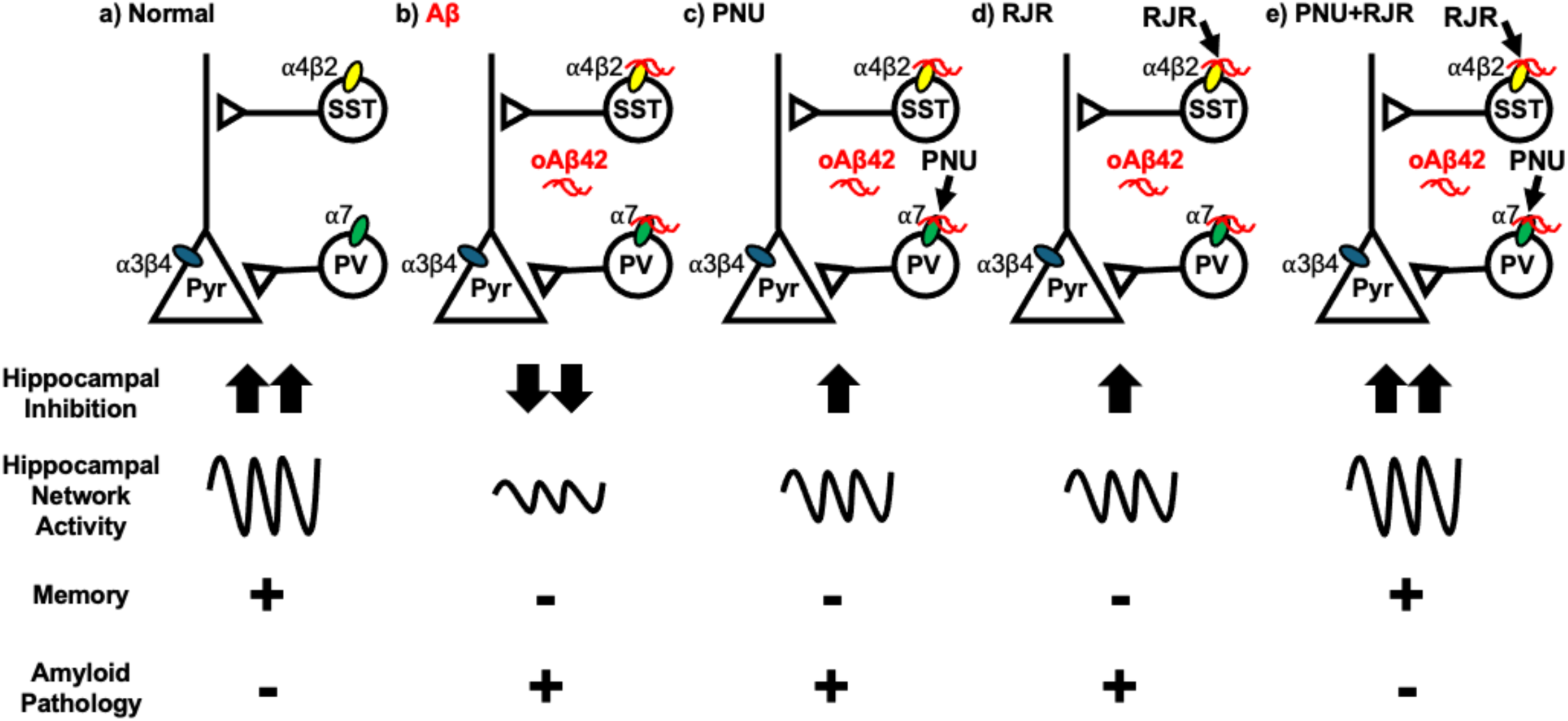
A schematic model of nAChR-mediated mechanisms of Aβ-induced dysfunction in hippocampal inhibition, network activities, amyloid pathology, and memory in AD. **a)** α7-, α4β2-, and α3β4-nAChRs predominantly control the nicotinic cholinergic signaling in parvalbumin-positive (PV+), somatostatin-positive (SST+), and pyramidal cells (Pyr), respectively, in the hippocampus. The cholinergic regulation of inhibitory activity contributes to inhibition onto pyramidal cells, which controls hippocampal network activity and memory processes. **b)** Aβ selectively inhibits α7- and α4β2-nAChRs in PV+ and SST+ cells, respectively, but not α3β4-nAChRs in excitatory neurons, which reduces inhibition to induce hyperexcitability and network dysfunction in the hippocampus, leading to memory loss in AD. Additionally, Aβ-induced hippocampal hyperexcitability increases Aβ secretion to enhance the amyloid pathology. Stimulation of PV+ or SST+ cells by the activation of each nAChR subtype-specific agonists, **c)** PNU-282987 (PNU), an α7 agonist, or **d)** RJR-2403 Oxalate (RJR), an α4β2 agonist, is insufficient to reverse the adverse effects of Aβ. **e)** Co-stimulation of hippocampal PV+ and SST+ inhibitory interneurons via activation of α7- and α4β2-nAChRs together is required to be protective against AD.

Aβ is suggested to have profound effects on NMDA and AMPA receptors (NMDARs and AMPARs) via the direct interaction. However, there is insufficient data supporting this idea. First, only immunocytochemistry data supports that Aβ can bind to NMDARs (Taniguchi et al. 2022). There are no confirmed studies demonstrating direct physical binding of Aβ oligomers to AMPAR subunits. Conversely, multiple experimental data using immunocytochemistry (Wang et al. 2000), co-immunoprecipitation (Wang et al. 2000; Roberts et al. 2021), and a specific and quantitative time-resolved FRET (TR-FRET)-based binding assay (Cecon et al. 2019) demonstrate that Aβ binds to nAChRs, particularly the α7- and α4β2-subtypes. In fact, we provide computational and biochemical evidence showing that Aβ directly associates with α7- and α4β2-nAChRs (Roberts et al. 2021). Second, it is known that Aβ affects excitatory and inhibitory neurons differently (Palop and Mucke 2016). Since both excitatory and inhibitory cells express AMPARs and NMDARs, it is unknown how Aβ modifies these receptors to influence excitatory and inhibitory neurons differently. Importantly, we demonstrate that AMPAR dysfunction is downstream of Aβ-induced imbalance of excitation and inhibition by blocking α7- and α4β2-nAChRs in hippocampal neurons (Roberts et al. 2021). According to these findings, the effects of Aβ on AMPARs and NMDARs are unlikely to be the main pathogenic mechanism in AD.

Theta oscillations in the hippocampus are generated by cholinergic input from the medial septum (MS) and is highly sensitive to nAChRs. Thus, they can be more responsive to agonist stimulation than other frequencies (Cobb et al. 1999; Dannenberg et al. 2017). In contrast, the local inhibitory interneuron networks play important roles in both slow and fast gamma oscillations, which makes them less sensitive to cholinergic inputs (Dannenberg et al. 2017). In fact, we find that PNU or RJR treatment reduces theta power but have no major effect on slow and fast gamma powers in WT mice (**Fig. 5a**). This reduction may have been observed due to excessive nAChR activation. Importantly, we do not see this same reduction of theta rhythm in 5XFAD mice because the baseline cholinergic signaling is already impaired, and we are able to see that co-activation of α7 and α4β2 nAChRs is able to restore to the optimal cholinergic tone. Therefore, agonist treatment likely enhances normal cholinergic activity in the WT hippocampus, resulting in this selective reduction in theta power, whereas local interneuron networks remain stable and resilient apart from this pharmacological activation.

Aβ can inhibit glial nAChRs on astrocytes and microglia rather than neuronal receptors to contribute to AD pathology. Therefore, stimulation of nAChR on glial cells may induce the protective effects. Although roles of astrocytic and microglial nAChRs in AD have not been fully understood to date (Sadigh-Eteghad et al. 2016), it has been shown that stimulation of astrocytic and microglial α7-nAChRs enhances Aβ phagocytosis in astrocytes and microglia to reduce Aβ levels (Takata et al. 2010; Yu et al. 2012). Hippocampal Aβ in AD model mice can thus inhibit glial α7-nAChRs to reduce phagocytic activity in these cells, resulting in an increase in Aβ levels to promote disease progress. nAChR agonist treatment would stimulate glial α7-nAChRs to reduce Aβ levels in the AD hippocampus by enhancing phagocytosis, leading to the protective effects. Thus, reduction of Aβ buildup in the 5XFAD hippocampus by PNU and RJR treatment in **Fig. 7** can be directly mediated by activating glial nAChRs rather than a decrease in neuronal hyperexcitability. In addition, activation of astrocytic α7- and α4β2-nAChRs promotes the expression of glial cell-derived neurotrophic factor (GDNF) (Liu et al. 2015; Oikawa et al. 2005; Takarada et al. 2012) that has been shown the protective roles in neurodegenerative disorders characterized mainly by damage of cholinergic neurons, such as AD (Straten et al. 2009). Thus, Aβ can inhibit astrocytic α7- and α4β2-nAChRs to reduce GDNF levels, resulting in the pathology, while activation of these receptors may induce the protective effects by increasing GDNF levels. This is important future research to differentiate between the potential contributions of neuronal and glial nAChRs to AD pathology.

One important question is why co-activation of PV+ and SST+ cells is required for neuroprotection against AD. PV+ interneurons provide perisomatic inhibition to control pyramidal cell output, whereas SST+ interneurons regulate dendritic excitation and input integration (Pelkey et al. 2017). Hippocampal network function is known to depend on coordinated activity of these cells. For example, it has been shown that hippocampal interneuron classes exhibit subtype-specific yet precisely coordinated firing patterns during network oscillations, indicating that different interneurons contribute complementary forms of inhibition to regulate pyramidal neuron activity and maintain circuit integrity (Varga et al. 2012). Additionally, recent work further demonstrates that PV+ and SST+ interneuron families cooperatively regulate hippocampal pyramidal neuron activity to establish accurate spatial representations (Valero et al. 2025). Disruption of either inhibitory pathway can thus compromise hippocampal network stability, highlighting the importance of restoring both PV+ and SST+ interneuron function to correct pathological hyperexcitability (Valero et al. 2025). Together, these studies establish that PV+ and SST+ interneurons regulate distinct but interdependent domains of hippocampal excitation, with PV+ cells constraining excessive somatic firing and SST+ cells limiting pathological dendritic excitation. Therefore, simultaneous restoration of PV+ and SST+ interneuron activity may be necessary to fully re-establish inhibitory control, suppress pathological hippocampal hyperexcitability, and provide neuroprotection.

Finally, the current work provides an opportunity to revisit the cholinergic hypothesis. Unfortunately, attempts to restore normal cholinergic activity using acetylcholinesterase inhibitors have only been moderately successful. Most of the previous research has focused on single nAChR subtype stimulation or non-selective activation in several brain disorders (Bitner et al. 2007; Cincotta et al. 2008; Redrobe et al. 2009; Vicens et al. 2011; Buccafusco 2004; Potasiewicz et al. 2018; Upright and Baxter 2021; Kantrowitz et al. 2020). It is also difficult to identify optimal combinations of nAChR agonists, thus co-activation of selective nAChRs has not been applied for both preclinical studies and clinical trials yet in AD (Buccafusco 2004). Identifying co-activation of α7- and α4β2-nAChRs as a protective approach on hippocampal functions and memory in AD will thus provide a new perspective on the cholinergic hypothesis.

## Materials and Methods

### Animals

C57Bl6J (Jax 000664), PV-Cre (Jax 017320), SST-Cre (Jax 018973), B6SJLF1/J (Ja× 100012), and 5XFAD (Jax 034840) mice were purchased from Jackson laboratory and bred in the animal facility at Colorado State University (CSU). Clinical studies are observational and limited in the mechanistic information that can be tested in living patients. In contrast, animal studies permit the manipulation of variables and the investigation of effects on an entire organism. The animal model, however, is disadvantaged by species differences. AD is a strictly human disease, and we are aware that rodents do not develop true AD (Elder et al. 2010). Nonetheless, multiple transgenic mouse models develop the pathological hallmarks of the disease as they age, like those found in humans (Elder et al. 2010; Hsiao et al. 1996). We thus employed a widely used amyloid pathology 5XFAD mouse models of AD. 5XFAD mice express human amyloid precursor protein (APP) and human presenilin 1 with the Familial Alzheimer’s Disease (FAD) mutations (Oakley et al. 2006; Oblak et al. 2021). Interestingly, biphasic changes in hippocampal circuit excitability are documented along the course of AD, with hyperactivity in the disease’s preclinical stages and hypoactivity in its later stages (Alexander et al. 2002; Busche and Konnerth 2016; Dickerson et al. 2005; Harris et al. 2020; Toniolo et al. 2020). Importantly, 5XFAD mice show hippocampal hyperexcitability as early as 2.5 months of age (Li et al. 2021). Additionally, at the age of 5 months, Aβ levels are significantly elevated in 5XFAD mice, which is associated with neuronal dysfunction and memory loss (Seo et al. 2014; Wang et al. 2019; Kimura and Ohno 2009; Oakley et al. 2006; Devi and Ohno 2010; Xiao et al. 2015). Moreover, deep proteome profiling of the hippocampus shows no major difference in nAChR expression between 5XFAD mice and wild-type (WT) littermate controls at the age of 5 months (Kim et al. 2019). However, cholinergic neuron loss is found after the age of 9 months in 5XFAD mice (Yan et al. 2018; Devi and Ohno 2015). These findings suggest that in 5-month-old 5XFAD mice, Aβ-induced nAChR dysfunction occurs at the functional levels as opposed to the receptor expression levels. Hippocampal network dysfunction also reported in this model at the age of 3-6 months (Iaccarino et al. 2016; Martorell et al. 2019). 5-month-old 5XFAD mice are thus the ideal amyloid pathology animal model for the current study. To collect brain tissues from animals, mice were deeply anesthetized and euthanized by CO_2_ asphyxiation. Animals were housed under a 12:12 hour light/dark cycle. CSU’s Institutional Animal Care and Use Committee (IACUC) reviewed and approved the animal care and protocol (3408). Results were reported following the ARRIVE (Animal Research: Reporting of In Vivo Experiments) guidelines (Kilkenny et al. 2010).

### Primary hippocampal neuronal culture

Postnatal day 0 (P0) male and female C57Bl6J, PV-Cre, or SST-Cre pups were used to produce mouse hippocampal neuron cultures as shown previously (Sathler et al. 2021; Sztukowski et al. 2018; Sathler et al. 2022). Hippocampi were isolated from P0 mouse brain tissues and digested with 10 U/ml papain (Worthington Biochemical Corp., LK003176). Mouse hippocampal neurons were plated on following poly lysine-coated dishes for glass bottom dishes (500,000 cells) for Ca^2+^ imaging. Neurons were grown in Neurobasal Medium without phenol red (Thermo Fisher Scientific, 12348017) with B27 supplement (Thermo Fisher Scientific, 17504044), 0.5 mM Glutamax (Thermo Fisher Scientific, 35050061), and 1% penicillin/streptomycin (Thermo Fisher Scientific, 15070063). The previous study evaluates maturation, aging, and death of mouse cortical cultured neurons for 60 days *in vitro* (DIV), which demonstrates that synaptogenesis is prominent during the first 15 days and then synaptic markers remained stable through 60 DIV (Lesuisse and Martin 2002). In particular, the levels of glutamate receptors increase to a maximum by 10-15 DIV and then remain unchanged through 60 DIV. This indicates that 12-14 DIV neurons that we used here are mature cells, and their maturity is likely comparable to that of older neurons. Additionally, neuronal cultures are shown to contain excitatory and inhibitory cells, cholinergic neurons, as well as glia (Roberts et al. 2021; Sun et al. 2019; Kaech and Banker 2006).

### Reagents

Soluble Aβ42 oligomers (oAβ42) were prepared as previously described (Stine et al. 2003; Sun et al. 2019). 1 mg of lyophilized human Aβ42 (Anaspec) was dissolved in 1ml of 1,1,1,3,3,3-hexafluoro-2-propanol (HFIP) (Sigma, St. Louis, MO) to prevent aggregation, portioned into 10 μg aliquots, air-dried and stored at −80 °C. For use in experiments, an aliquot was thawed at room temperature and then dissolved in DMSO to make a 100 μM solution. The solution was incubated for 16 hours at 4°C and then diluted to a final concentration for use in experiments. Scrambled Aβ42 oligomers (sAβ42) were prepared the same way for controls. The following nAChR antagonists were used in this study: α-Bungarotoxin (αBTx) (Alomone labs), an α7 antagonist, Dihydro-β-erythroidine hydrobromide (DHβE) (Tocris Bioscience), an α4β2 antagonist, and α-Conotoxin AuIB (Alomone labs), an α3β4 antagonist. The following nAChR agonists were used in this study: PNU-282987 (PNU) (Tocris Bioscience), an α7 agonist, RJR-2403 Oxalate (RJR) (Tocris Bioscience), an α4β2 agonist, and NS-3861 (NS) (Tocris Bioscience), an α3β4 agonist. Tetrodotoxin (TTX) (Abcam) was used to prevent action potential-dependent spontaneous network activity.

### GCaMP Ca^2+^ imaging with nicotine uncaging

For Ca^2+^ imaging with nicotine uncaging, a genetically encoded Ca^2+^ indicator, GCaMP, was used. As the majority of cells in hippocampal cultures are excitatory neurons (Benson and Cohen 1996), we expressed GCaMP6f (Addgene #40755) (Chen et al. 2013) under the control of the CMV promoter by transfection for imaging excitatory cells, and GCaMP6f under the control of the GABAergic neuron-specific enhancer of the mouse *Dlx* (mDlx) gene (Addgene #83899) (Dimidschstein et al. 2016) was transfected for imaging inhibitory interneurons as shown previously (Sun et al. 2019). Glass-bottom dishes were mounted on a temperature-controlled stage on an Olympus IX73 microscope and maintained at 37°C and 5% CO_2_ using a Tokai-Hit heating stage and digital temperature and humidity controller. For nicotine uncaging, 1 μM photoactivatable nicotine (PA-Nic) (Tocris Bioscience) was added to the culture media, and epi-illumination photolysis (390 nm, 0.12 mW/mm^2^, 1 sec) was used as shown previously (Banala et al. 2018). TTX (1 μM) was added in culture media to prevent action potential-dependent spontaneous network activity. A baseline average (F_0_) of 20 frames (50 ms exposure) were captured in the soma prior to nicotine uncaging, and 50 more frames (50 ms exposure) were obtained after photostimulation. The fractional change in fluorescence intensity relative to baseline (ΔF/F_0_) was calculated. For **Fig. 1a**, the average peak amplitude in excitatory cells (EX) was used to normalize the peak amplitude in each cell and was compared to the inhibitory cells’ average (IN). For **Fig. 1b and 1c**, the images were captured right after each antagonist was added to the media. The average peak amplitude in the control group without drug treatment (CTRL) was used to normalize the peak amplitude in each cell, and the control group’s average peak amplitude was compared to the experimental groups’ average. For **Fig. 1d and 1e**, PV-Cre and SST-Cre mice were used to culture hippocampal neurons. We infected 4 DIV neurons prepared from PV-Cre or SST-Cre mice with adeno-associated virus (AAV) expressing Cre-dependent GCaMP7s (Addgene# 104495-AAV1) - pGP-AAV-CAG-FLEX-jGCaMP7s-WPRE was a gift from Douglas Kim & GENIE Project (Addgene plasmid #104495; http://n2t.net/addgene:104495; RRID:Addgene_104495) (Dana et al. 2019) to determine nicotinic cholinergic activity in PV+ and SST+ cells using nicotine uncaging. For GCaMP7s, the images were captured right after each antagonist was added to the media. A baseline average (F_0_) of 20 frames (5 ms exposure) were captured prior to nicotine uncaging, and 100 more frames (5 ms exposure) were obtained after photolysis. The average peak amplitude in the control group (CTRL) was used to normalize the peak amplitude in each cell, and the control group’s average peak amplitude was compared to the experimental groups’ average. For Fig. 2, PV-Cre and SST-Cre mice were also used to culture hippocampal neurons. For **Fig 2a and 2b**, Ca^2+^ imaging with nicotine uncaging was conducted as described in **Fig. 1d and 1e**. The images were captured right after 250 nM sAβ42 or 250 nM oAβ42 was added to the media. The average peak amplitude in sAβ42-treated cells was used to normalize the peak amplitude in each cell, and the sAβ42 group’s average peak amplitude was compared to the oAβ42-treated cells’ average.

### GCaMP Ca^2+^ imaging

We measured spontaneous Ca^2+^ activity in cultured hippocampal excitatory neurons because It has been shown that networks of neurons in culture can produce spontaneous synchronized activity (Cohen et al. 2008). In fact, network activity emerges at 3-7 DIV independent of either ongoing excitatory or inhibitory synaptic activity and matures over the following several weeks in cultures (Cohen et al. 2008). Therefore, the somatic Ca^2+^ signals we observed are from the spontaneous network activity in cultured cells. To do this, we used AAV to express Cre-dependent GCaMP7s (Addgene# 104495-AAV1) (Dana et al. 2019) in cells prepared from PV-Cre or SST-Cre mice. We then measured Ca^2+^ activity in the soma of 14 DIV cultured hippocampal excitatory neurons with a modification of the previously described method (Kim, Titcombe, et al. 2015; Kim, Violette, et al. 2015; Kim and Ziff 2014; Roberts et al. 2021; Sun et al. 2019; Sztukowski et al. 2018; Zaytseva et al. 2023). For GCaMP7s, the images were captured right after 250 nM sAβ42 or 250 nM oAβ42 was added to the media in the presence or absence of each agonist. A 10 ms exposure time and a total of 100 images were obtained with a one-second interval. F_min_ was determined as the minimum fluorescence value during the imaging. Total Ca^2+^ activity was obtained by 100 values of ΔF/F_min_ = (F_t_ – F_min_) / F_min_ in each image, and values of ΔF/F_min_ < 0.1 were rejected due to potential photobleaching. The average total Ca^2+^ activity in the control group (CTRL) was used to normalize total Ca^2+^ activity in each cell. The control group’s average total Ca^2+^ activity was compared to the experimental groups’ average as described previously (Kim, Titcombe, et al. 2015; Kim, Violette, et al. 2015; Kim and Ziff 2014; Sztukowski et al. 2018; Roberts et al. 2021; Sun et al. 2019).

### *In vivo* activation of nAChRs

For *in vivo* activation of nAChRs, we used PNU-282987 (PNU), an α7-nAChR agonist, and RJR-2403 Oxalate (RJR), an α4β2-nAChR agonist. These agonists have a number of advantages that can be used to test our hypothesis; 1) they are blood-brain barrier (BBB) permeable (Hijioka et al. 2012; Vicens et al. 2011; Redrobe et al. 2009; Levin and Christopher 2002; Bencherif et al. 1996), 2) they do not require endogenous acetylcholine to stimulate receptors since they are direct agonists rather than positive allosteric modulators (Hijioka et al. 2012; Niimi et al. 2013; Duris et al. 2011; Kawamura et al. 2022; Hajos et al. 2005; Vicens et al. 2011; Redrobe et al. 2009), 3) repeated injection of these agonists is sufficient to show receptor activation in the rodent brains without causing receptor desensitization (Smith and Dar 2007; Lippiello et al. 1996; Sapronov et al. 2006; Duris et al. 2011; Kawamura et al. 2022), and 4) many α7-nAChR agonists including PNU are known to act as serotonin 5-HT_3_ receptor antagonists due to high structural homology between the receptors. However, the previous study has demonstrated that PNU is unable to affect hippocampal network activity in α7-nAChR knockout mice, suggesting that effects of PNU on hippocampal oscillatory activity are exclusively mediated via α7-nAChRs rather than 5-HT_3_ receptors (Stoiljkovic et al. 2016). Therefore, effects of PNU via 5-HT_3_ receptor antagonism are unlikely. PNU and RJR were thus prepared at concentrations of 1 mg/ml and intraperitoneally injected into 5-month-old 5XFAD and WT female and male mice at a dose of 5 mg/kg by themselves or together once per day for 7 days, a condition that is sufficient to stimulate each nAChR subtype *in vivo* (Hijioka et al. 2012; Vicens et al. 2011; Redrobe et al. 2009; Levin and Christopher 2002). Saline was given to mice as a control.

### Immunohistochemistry

c-Fos labeling was used to measure cell-type specific neuronal activity as shown previously (Tran et al. 2021; Roh et al. 2024) because this provides activity patterns across multiple cell types simultaneously. We conducted immunohistochemistry with 5-month-old 5XFAD mice and WT littermates. Brain tissues were isolated and sectioned at 40 μm by using a vibratome. 10 consecutive sections containing the hippocampus (from Bregma: −1.64mm) in each mouse were stained with anti-PV, anti-SST, and anti-c-Fos antibodies. Nuclei were also labeled with DAPI as shown previously (Tran et al. 2021; Roh et al. 2024). We imaged these sections using Olympus inverted microscope IX73 with the CellSens software. The colocalization analysis function in the CellSens program was utilized to identify c-Fos-positive (c-Fos+) signals on DAPI-positive (DAPI+) cells but PV and SST-negative cells as the activity of pyramidal cells. Similarly, we measured interneurons’ activity by identifying c-Fos+ signals on PV+ or SST+ cells. The total number of c-Fos+ cells on each neuronal subtype in the hippocampus was counted using the particle analysis function in ImageJ (threshold = 100 μm^2^). In addition, we measured the size of the hippocampus using ImageJ and normalize the total number of c-Fos+ cells on each neuronal subtype per 5.0 × 10^6^ μm^2^. As no overt hippocampal neuron loss is found by 12-month-old 5XFAD mice (Jawhar et al. 2012), cell-type specific neuronal activity was measured as an average of the total number of c-Fos+ cells on each cell subtype/5.0 × 10^6^ μm^2^ from each section. The average counts per section across all experimental conditions were then compared because multiple immunostainings were performed on the same anatomical sections, analyses were conducted at the section level to assess within-section differences in marker-defined cell populations.

### Stereotaxic surgery

5-month-old mice were anaesthetized with an intraperitoneal injection of a mixture of ketamine (100 mg/kg) and xylazine (10 mg/kg) and placed in a stereotactic frame. The target craniotomy site for local field potential (LFP) recordings was marked on the skull, and electrodes were implanted bilaterally into the dorsal hippocampal CA1 area (Bregma coordinates: AP: −1.95 mm, ML: ± 1.12 mm, DV: −1.20 mm). One small screw for the ground anchored at the anterior edges of the surgical site and a wire for a reference in the posterior part of the craniotomy site were bound with dental cement (GC America) to secure the implant in place. The electrodes were constructed by twisting 16 Formvar-insulated NiCh wires (75 μm, California Fine Wire) and attached to a Mill-Max connector. After surgery, animals’ health was closely monitored as described in our approved IACUC protocol.

### Hippocampal LFP recording

We conducted contextual fear conditioning one week after the surgery as a learning and memory paradigm. Basal LFPs were recorded before learning for one hour (**Fig. 4a**). According to a large body of studies, consolidation of hippocampus-dependent fear memory occurs between 1 and 3 hours after training (Kwapis and Wood 2014; Bourtchouladze et al. 1998). Therefore, after fear learning, test animals were returned to their home cage for 90 minutes, and then we recorded hippocampal LFPs for one hour, a time for fear memory consolidation (**Fig. 4a**). LFP data were collected through electrodes which connect to a 16-channel head-stage (Intan Technologies) that digitizes the raw analog LFP signals via an onboard analog-to-digital converter. We used the Open Ephys Acquisition Board (Siegle et al. 2017) to acquire the digitized signal at 2 kHz. The electrode locations were verified histologically at the end of recordings (**Fig. 4b**). If the electrode was placed outside the CA1 area, the animal was excluded in analysis.

### LFP data analysis

LFP data were analyzed in MATLAB (The Mathworks Inc.). The data were formatted for importation into the Brainstorm platform (Tadel et al. 2011). Recorded traces were downsampled to 2 kHz and then bandpass filtered between 0.1 and 500 Hz (**Fig. 4c**). We applied a notch filter at 60 Hz to reduce powerline artifacts. Power spectral density was calculated using Welch’s method in the Brainstorm platform (Tadel et al. 2011), which involves segmenting the signals and applies a Fast Fourier Transform (FFT) algorithm to each segment before averaging the power spectra (**Fig. 4d**). The power spectra were then processed by averaging frequency bins into larger predefined bands, including theta (4-8 Hz), slow gamma (30-50 Hz), and fast gamma (50-100 Hz). The power of each frequency band was normalized to WT controls and then compared before and after fear conditioning in WT and 5XFAD mice. We also compared the powers in each condition between WT and 5XFAD mice following learning. LFP power was quantified for each electrode and frequency band because multiple electrodes were obtained from the same animal. For theta-gamma phase amplitude coupling (PAC) analysis, we measured the high-frequency bands modulated by theta bands in epochs with a matched, filtered LFP amplitude envelope. Data were filtered into theta, slow gamma, and fast gamma using a zero-phase FIR filter implemented with the eegfilt function in the EEGLAB toolbox for MATLAB (Delorme and Makeig 2004). The analytic signal for each was computed using the Hilbert Transform (MATLAB function hilbert.m), resulting in instantaneous theta phase and gamma band amplitude. The peaks of the gamma-band amplitudes were defined as local maxima at or exceeding the mean + 1 standard deviation (SD) of the baseline gamma-band amplitude from the Hilbert Transform envelopes. The instantaneous theta phase for each peak was extracted. The mean vector length (MVL) was calculated as an entrainment strength. We compared the MVL before and after fear conditioning. The MVL in each condition was also compared in WT and 5XFAD mice following fear learning.

### Novel object location (NOL) test

Because the NOL test primarily evaluates spatial learning, which relies heavily on hippocampal activity, we carried out the NOL test to examine hippocampus-dependent memory as described previously (Denninger et al. 2018). Animals were spent 10 minutes in a round social arena (Diameter: 25cm) for habituation on Day 1. The following day (Day 2), animals were habituated in the same chamber again for 5 minutes. Two different objects (object 1 and 2) were placed in the chamber for training, and a mouse was free to explore for 10 minutes and went back to the home cage for 10 minutes. The object 2 was then moved to a new location, and the animal was free to explore for additional 10 minutes. The behavior was recorded with a video camera, and the amount of time spent in each object will be determined by an automated tracking program (Anymaze) that measured how long a mouse’s head was directed toward the object in a 30-degree head orientation and came within 2 cm of it. Object preference was quantified as a preference index (PI), calculated as the time with the object 2 / the time with the object 1 + the time with the object 2 as shown previously (Denninger et al. 2018). Because individual mice may exhibit baseline preferences for one object during training (Tropea et al. 2022), PI was calculated separately for each mouse during the training and testing sessions, using the same designated object for both sessions. Training and testing PIs were then compared within animals. This within-subject analysis was used to determine whether relocation of the object produced a significant change in object preference relative to each animal’s baseline preference. Normal spatial memory was operationally defined as an increase in the PI for the object relocated during testing relative to the PI measured during the training session.

### Contextual fear conditioning (CFC)

Contextual fear conditioning (Habitest Modular System, Coulbourn Instrument) was carried out as described previously (Kim et al. 2016; Shou et al. 2018). In Day 1 (encoding), the test mouse was placed in a novel rectangular chamber with a grid floor. After a 3-min baseline period, the test animal was given one shock (a 2 sec, 0.5 mA shock) and stayed in the chamber for an additional 1 min after the shock before being returned to the home cage for overnight (consolidation). A contextual memory test was conducted the next day (Day 2 - retrieval) in the same conditioning chamber for 3 min. Fear memory was determined by measuring the percentage of the freezing response (immobility excluding respiration and heartbeat) using an automated tracking program (FreezeFrame).

### Amyloid plaque analysis

10 hippocampal sections (from Bregma: −1.64mm) in each mouse were obtained and stained with 1% Thioflavin S. Amyloid plaques were identified as Thioflavin S-positive puncta. The total number of amyloid plaques in the hippocampus was counted using the particle analysis function in ImageJ (threshold = 10 μm^2^). We also measured the size of the hippocampus using ImageJ and normalized these measurements per 1 mm^2^. Amyloid plaques were measured as an average of these measurements per 1 mm^2^ from 10 sections (total 20 hemisphere/animal). The average counts per animal across all experimental conditions were then compared between the conditions.

### Statistical analysis

The Franklin A. Graybill Statistical Laboratory at CSU was consulted for statistical analysis in the current study, including sample size determination, randomization, experiment conception and design, data analysis, and interpretation. We used the GraphPad Prism 11 software to determine statistical significance (set at *p* < 0.05). Grouped results of single comparisons were tested for normality with the Shapiro-Wilk normality or Kolmogorov-Smirnov test and analyzed using an unpaired or paired two-tailed Student’s t-test when data are normally distributed. Differences between multiple groups were assessed by N-way analysis of variance (ANOVA) with the Tukey test or Šídák’s test. Statistical analyses for LFP and immunostaining data were performed using mixed-effects models / repeated-measures ANOVA with Tukey test, and animal identity included as a random effect. Electrodes and sections in LFP and c-Fos analysis, respectively, were treated as repeated measures nested within animals, and animal identity was included as a random effect. This approach accounts for the nested structure of the data, avoids pseudoreplication, and accommodates unequal electrode counts across animals. The graphs were presented as mean ± Standard Deviation (SD).

## Acknowledgments

We thank members of the Kim laboratory for their generous support. We appreciate Dr. Eunhye Park for the technical guidance and Dr. Don Rojas to support data analysis. This work is supported by the NIH/NCATS Colorado CTSA Grant (UL1 TR002535), the Boettcher Foundation’s Webb-Waring Biomedical Research Program, BrightFocus Foundation, and the NIH grant (1R03AG072102), and Colorado State University College of Veterinary Medicine and Biomedical Sciences.

## Author contributions

Conceptualization: S.K. Methodology: S.K. Validation: R.L. G.K. and S.K. Formal analysis: R.L. G.K. and S.K. Investigation: R.L. G.K. E.R.B. and S.K. Writing - Original Draft: R.L and S.K. Writing - Review & Editing: R.L and

S.K. Visualization: R.L. G.K. and S.K. Supervision: S.K. Project administration: S.K. Funding acquisition: E.R.B. and S.K.

