## Supplementary figures for "Co-activation of selective nicotinic acetylcholine receptor subtypes is required for neuroprotection against Alzheimer’s disease"

**Supplementary Fig 1.**

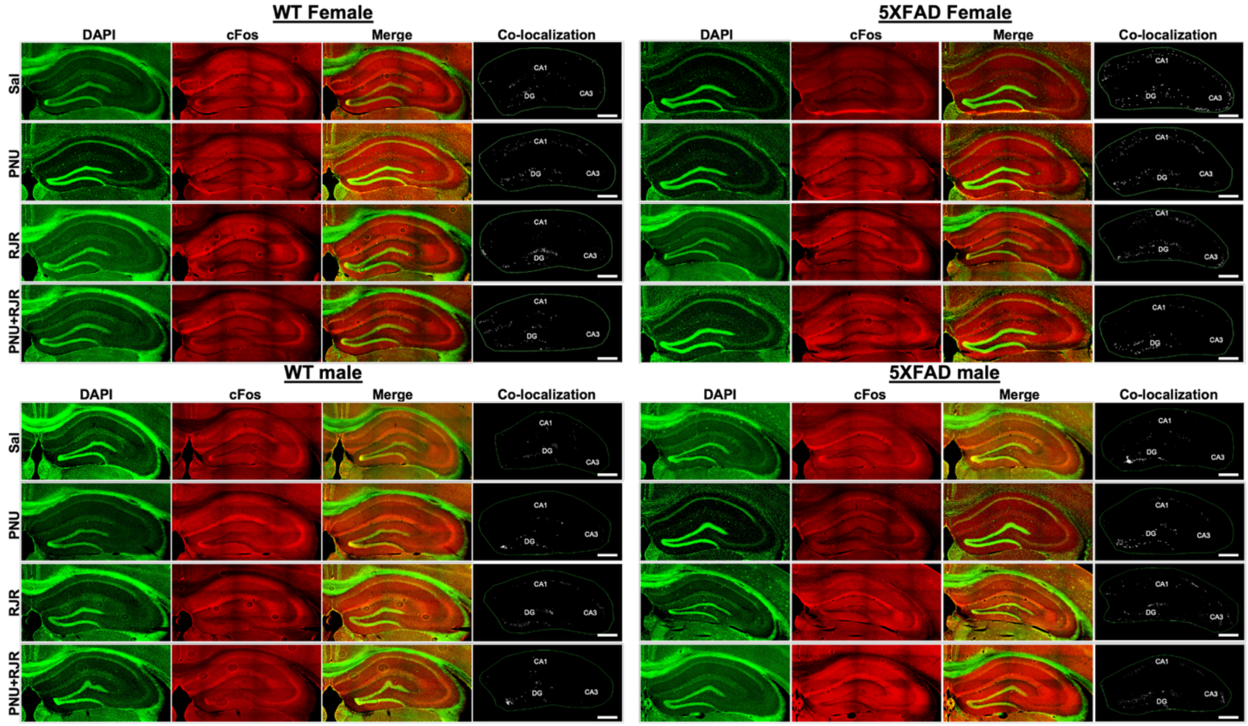


Representative hippocampal sections from the indicated experimental groups in Fig. 3b showing DAPI staining (green; left), c-Fos immunoreactivity (red; middle-left), and merged images (middle-right). Co-localization of c-Fos immunoreactivity with DAPI-labeled nuclei (c-Fos+/DAPI+ cells) identifies c-Fos-positive activated cells, including pyramidal neurons within the hippocampus. Corresponding hippocampal maps (right) show the distribution of c-Fos+/DAPI+ cells within the CA1, dentate gyrus (DG), and CA3 subregions. Outlines show hippocampal region of interest (ROI). Scale bars indicate 500 μm.

**Supplementary Fig 2.**

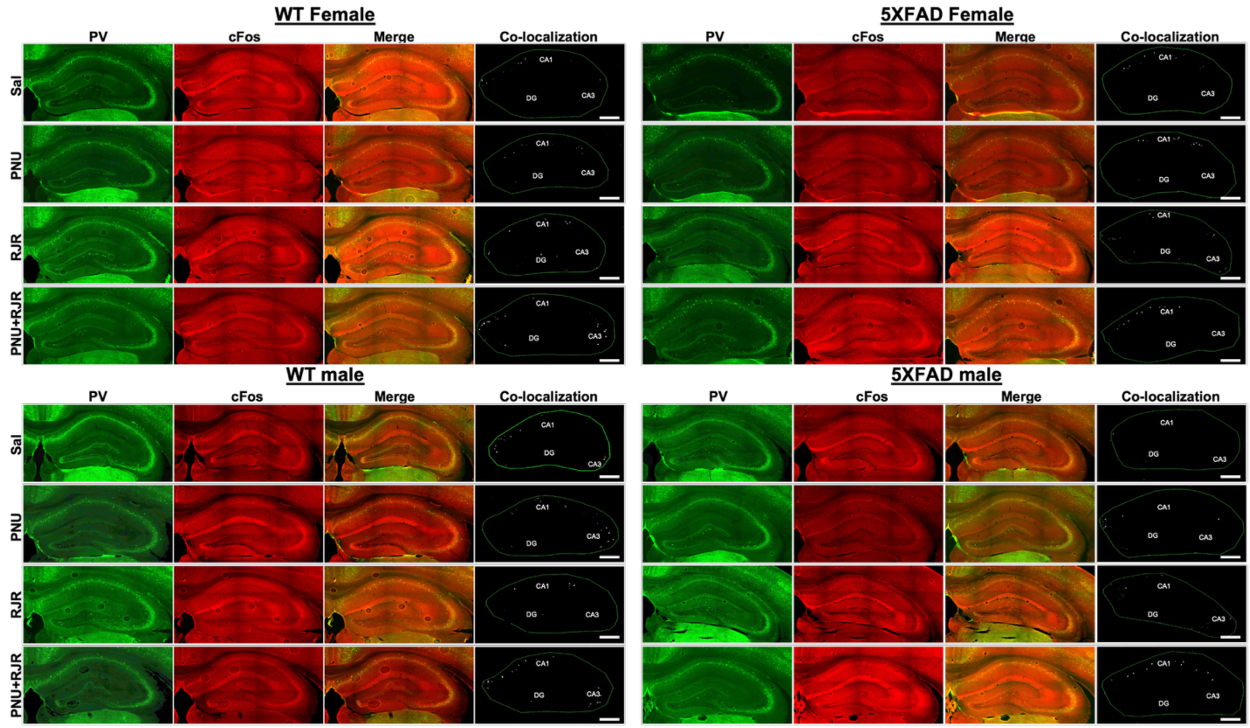


Representative hippocampal sections from the indicated experimental groups in Fig. 3c showing PV staining (green; left), c-Fos immunoreactivity (red; middle-left), and merged images (middle-right). Co-localization of c-Fos immunoreactivity with PV+ interneurons (c-Fos+/PV+ cells) identifies c-Fos-positive activated PV+ cells within the hippocampus. Corresponding hippocampal maps (right) show the distribution of c-Fos+/PV+ cells within the CA1, dentate gyrus (DG), and CA3 subregions. Outlines show hippocampal region of interest (ROI). Scale bars indicate 500 μm.

**Supplementary Fig 3.**

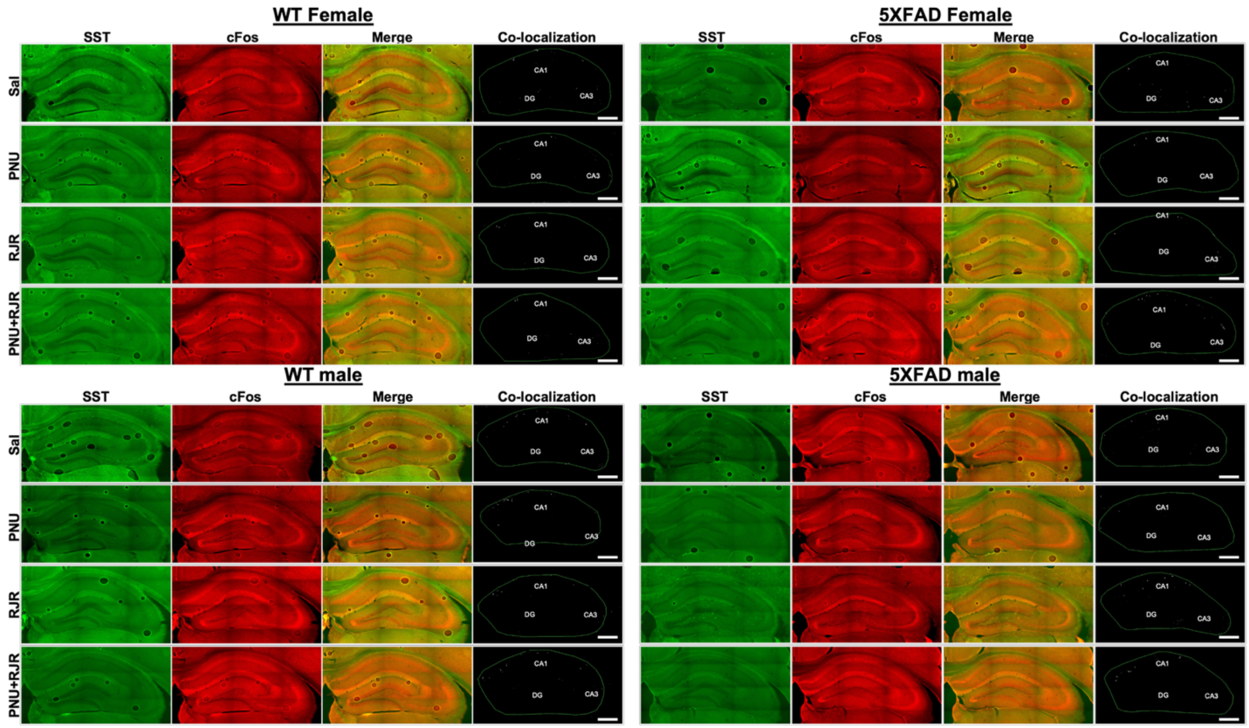


Representative hippocampal sections from the indicated experimental groups in Fig. 3d showing SST staining (green; left), c-Fos immunoreactivity (red; middle-left), and merged images (middle-right). Co-localization of c-Fos immunoreactivity with SST+ interneurons (c-Fos+/SST+ cells) identifies c-Fos-positive activated SST+ cells within the hippocampus. Corresponding hippocampal maps (right) show the distribution of c-Fos+/SST+ cells within the CA1, dentate gyrus (DG), and CA3 subregions. Outlines show hippocampal region of interest (ROI). Scale bars indicate 500 μm.
