## Supplementary Tables for "Co-activation of selective nicotinic acetylcholine receptor subtypes is required for neuroprotection against Alzheimer’s disease"

**Supplementary Table 1. Statistical analysis of Ca^2+^ imaging with nicotine uncaging in Fig. 1a.**

| **Normalized Peak Amplitude (ΔF/F_0_)** | |
| --- | --- |
| EX  IN | 1.000 ± 0.879 ΔF/F_0_  1.575 ± 1.208 ΔF/F_0_ |
| Unpaired t test | |
| P value | 0.0109 |
| t, df | t=2.601, df=88 |

**Supplementary Table 2. Statistical analysis of Ca^2+^ imaging with nicotine uncaging in Fig. 1b and 1c.**

| **Excitatory cells** | | | |
| --- | --- | --- | --- |
| **Normalized Peak Amplitude (ΔF/F_0_)** | | | |
|  | CTRL  αBTx  DhβE  αCTx | 1.000 ± 0.492 ΔF/F_0_  0.870 ± 0.328 ΔF/F_0_  1.042 ± 0.644 ΔF/F_0_  0.325 ± 0.362 ΔF/F_0_ | |
| ANOVA summary |  |  | |
| F | 7.453 |  |  |
| P value | 0.0002 |  |  |
| P value summary | *** |  |  |
| Significant diff. among means (P < 0.05)? | Yes |  |  |
| R squared | 0.2619 |  | |
| Tukey's multiple comparisons test | Mean diff. | 95.00% CI of diff. | Adjusted P Value |
| CTRL vs. αBTx | 0.1299 | -0.3451 to 0.6048 | 0.8881 |
| CTRL vs. DHβE | -0.04198 | -0.4552 to 0.3712 | 0.9932 |
| CTRL vs. αCTx | 0.6746 | 0.2615 to 1.088 | 0.0003 |
| αBTx vs. DHβE | -0.1719 | -0.6957 to 0.3519 | 0.8224 |
| αBTx vs. αCTx | 0.5448 | 0.02098 to 1.069 | 0.0385 |
| DHβE vs. αCTx | 0.7166 | 0.2481 to 1.185 | 0.0008 |
| **Inhibitory interneurons** | | | |
| **Normalized Peak Amplitude (ΔF/F_0_)** | | | |
|  | CTRL  αBTx  DhβE  αCTx | 1.000 ± 0.525 ΔF/F_0_  0.398 ± 0.290 ΔF/F_0_  0.335 ± 0.300 ΔF/F_0_  0.871 ± 0.330 ΔF/F_0_ | |
| ANOVA summary |  |  | |
| F | 10.91 |  |  |
| P value | <0.0001 |  |  |
| P value summary | **** |  |  |
| Significant diff. among means (P < 0.05)? | Yes |  |  |
| R squared | 0.3908 |  | |
| Tukey's multiple comparisons test | Mean diff. | 95.00% CI of diff. | Adjusted P Value |
| CTRL vs. αBTx | 0.6021 | 0.1752 to 1.029 | 0.0025 |
| CTRL vs. DHβE | 0.6648 | 0.2991 to 1.031 | <0.0001 |
| CTRL vs. αCTx | 0.1287 | -0.2371 to 0.4944 | 0.7866 |
| αBTx vs. DHβE | 0.06266 | -0.3538 to 0.4791 | 0.9782 |
| αBTx vs. αCTx | -0.4735 | -0.8899 to -0.05707 | 0.0200 |
| DHβE vs. αCTx | -0.5361 | -0.8895 to -0.1828 | 0.0010 |

**Supplementary Table 3. Statistical analysis of Ca^2+^ imaging with nicotine uncaging in Fig. 1d and 1e.**

| **PV+ interneurons** | | | |
| --- | --- | --- | --- |
| **Normalized Peak Amplitude (ΔF/F_0_)** | | | |
|  | CTRL  αBTx  DhβE  αBTx + DhβE | 1.000 ± 0.590 ΔF/F_0_  0.507 ± 0.374 ΔF/F_0_  1.076 ± 0.613 ΔF/F_0_  0.607 ± 0.448 ΔF/F_0_ | |
| ANOVA summary |  |  | |
| F | 10.97 |  |  |
| P value | <0.0001 |  |  |
| P value summary | **** |  |  |
| Significant diff. among means (P < 0.05)? | Yes |  |  |
| R squared | 0.1914 |  | |
| Tukey's multiple comparisons test | Mean diff. | 95.00% CI of diff. | Adjusted P Value |
| CTRL vs. αBTx | 0.4929 | 0.1728 to 0.8130 | 0.0006 |
| CTRL vs. DHβE | -0.07598 | -0.4090 to 0.2571 | 0.9340 |
| CTRL vs. αBTx + DhβE | 0.3934 | 0.08163 to 0.7052 | 0.0071 |
| αBTx vs. DHβE | -0.5689 | -0.8833 to -0.2544 | <0.0001 |
| αBTx vs. αBTx + DhβE | -0.09949 | -0.3913 to 0.1923 | 0.8118 |
| DHβE vs. αBTx + DhβE | 0.4694 | 0.1634 to 0.7754 | 0.0006 |
| **SST+ interneurons** | | | |
| **Normalized Peak Amplitude (ΔF/F_0_)** | | | |
|  | CTRL  αBTx  DhβE  αBTx + DhβE | 1.000 ± 0.426 ΔF/F_0_  0.917 ± 0.519 ΔF/F_0_  0.524 ± 0.262 ΔF/F_0_  0.533 ± 0.272 ΔF/F_0_ | |
| ANOVA summary |  |  | |
| F | 11.56 |  |  |
| P value | <0.0001 |  |  |
| P value summary | **** |  |  |
| Significant diff. among means (P < 0.05)? | Yes |  |  |
| R squared | 0.2614 |  | |
| Tukey's multiple comparisons test | Mean diff. | 95.00% CI of diff. | Adjusted P Value |
| CTRL vs. αBTx | 0.08337 | -0.2022 to 0.3689 | 0.8708 |
| CTRL vs. DHβE | 0.4764 | 0.2011 to 0.7518 | <0.0001 |
| CTRL vs. αBTx + DhβE | 0.4675 | 0.2071 to 0.7278 | <0.0001 |
| αBTx vs. DHβE | 0.3931 | 0.09978 to 0.6863 | 0.0038 |
| αBTx vs. αBTx + DhβE | 0.3841 | 0.1048 to 0.6634 | 0.0028 |
| DHβE vs. αBTx + DhβE | -0.008950 | -0.2777 to 0.2598 | 0.9998 |

**Supplementary Table 4. Statistical analysis of Ca^2+^ imaging with nicotine uncaging in Fig. 2a.**

| **PV+ interneurons** | |
| --- | --- |
| **Normalized Peak Amplitude (ΔF/F_0_)** | |
| sAβ42  oAβ42 | 1.000 ± 0.772 ΔF/F_0_  0.448 ± 0.233 ΔF/F_0_ |
| Unpaired t test | |
| P value | 0.0038 |
| t, df | t=3.032, df=53 |
| **SST+ interneurons** | |
| **Normalized Peak Amplitude (ΔF/F_0_)** | |
| EX  IN | 1.000 ± 0.391 ΔF/F_0_  0.600 ± 0.237 ΔF/F_0_ |
| Unpaired t test | |
| P value | 0.0002 |
| t, df | t=4.101, df=44 |

**Supplementary Table 5. Statistical analysis of Ca^2+^ imaging in Fig. 2c.**

| **PV+ interneurons** | | | | | |
| --- | --- | --- | --- | --- | --- |
| **Normalized Total ΔF/F_min_** | | | | | |
|  | | sAβ42  sAβ42 + PNU  sAβ42 + RJR  sAβ42 + NS  oAβ42  oAβ42 + PNU  oAβ42 + RJR  oAβ42 + NS | | 1.000 ± 0.379 ΔF/F_min_  1.417 ± 0.702 ΔF/F_min_  1.036 ± 0.558 ΔF/F_min_  1.070 ± 0.451 ΔF/F_min_  0.589 ± 0.223 ΔF/F_min_  1.408 ± 0.806 ΔF/F_min_  0.528 ± 0.299 ΔF/F_min_  0.550 ± 0.384 ΔF/F_min_ | |
| ANOVA table | SS (Type III) | DF | MS | F (DFn, DFd) | P value |
| Interaction | 1.647 | 3 | 0.5491 | F (3, 182) = 2.246 | P=0.0845 |
| Row Factor | 5.593 | 1 | 5.593 | F (1, 182) = 22.88 | P<0.0001 |
| Column Factor | 11.07 | 3 | 3.691 | F (3, 182) = 15.10 | P<0.0001 |
| Residual | 44.50 | 182 | 0.2445 |  | |
| Tukey's multiple comparisons test | | Mean diff. | | 95.00% CI of diff. | Adjusted P Value |
| sAβ42 vs. sAβ42 + PNU | | -0.4166 | | -0.8458 to 0.01249 | 0.0639 |
| sAβ42 vs. sAβ42 + RJR | | -0.03628 | | -0.3983 to 0.3257 | >0.9999 |
| sAβ42 vs. sAβ42 + NS | | -0.07011 | | -0.4681 to 0.3279 | 0.9994 |
| sAβ42 vs. oAβ42 | | 0.4115 | | 0.01348 to 0.8094 | 0.0371 |
| sAβ42 vs. oAβ42 + PNU | | -0.4079 | | -0.8127 to -0.003091 | 0.0468 |
| sAβ42 vs. oAβ42 + RJR | | 0.4722 | | 0.02258 to 0.9219 | 0.0320 |
| sAβ42 vs. oAβ42 + NS | | 0.4500 | | 0.05205 to 0.8480 | 0.0148 |
| sAβ42 + PNU vs. sAβ42 + RJR | | 0.3804 | | -0.08584 to 0.8466 | 0.2013 |
| sAβ42 + PNU vs. sAβ42 + NS | | 0.3465 | | -0.1482 to 0.8412 | 0.3884 |
| sAβ42 + PNU vs. oAβ42 | | 0.8281 | | 0.3334 to 1.323 | <0.0001 |
| sAβ42 + PNU vs. oAβ42 + PNU | | 0.008748 | | -0.4914 to 0.5089 | >0.9999 |
| sAβ42 + PNU vs. oAβ42 + RJR | | 0.8889 | | 0.3517 to 1.426 | <0.0001 |
| sAβ42 + PNU vs. oAβ42 + NS | | 0.8667 | | 0.3720 to 1.361 | <0.0001 |
| sAβ42 + RJR vs. sAβ42 +NS | | -0.03383 | | -0.4715 to 0.4039 | >0.9999 |
| sAβ42 + RJR vs. oAβ42 | | 0.4477 | | 0.01004 to 0.8854 | 0.0409 |
| sAβ42 + RJR vs. oAβ42 + PNU | | -0.3716 | | -0.8155 to 0.07230 | 0.1750 |
| sAβ42 + RJR vs. oAβ42 + RJR | | 0.5085 | | 0.02335 to 0.9937 | 0.0326 |
| sAβ42 + RJR vs. oAβ42 + NS | | 0.4863 | | 0.04862 to 0.9240 | 0.0180 |
| sAβ42 + NS vs. oAβ42 | | 0.4816 | | 0.01364 to 0.9495 | 0.0387 |
| sAβ42 + NS vs. oAβ42 + PNU | | -0.3378 | | -0.8115 to 0.1360 | 0.3648 |
| sAβ42 + NS vs. oAβ42 + RJR | | 0.5423 | | 0.02975 to 1.055 | 0.0298 |
| sAβ42 + NS vs. oAβ42 + NS | | 0.5201 | | 0.05222 to 0.9881 | 0.0179 |
| oAβ42 vs. oAβ42 + PNU | | -0.8194 | | -1.293 to -0.3456 | <0.0001 |
| oAβ42 vs. oAβ42 + RJR | | 0.06077 | | -0.4518 to 0.5734 | >0.9999 |
| oAβ42 vs. oAβ42 + NS | | 0.03858 | | -0.4293 to 0.5065 | >0.9999 |
| oAβ42 + PNU vs. oAβ42 + RJR | | 0.8801 | | 0.3622 to 1.398 | <0.0001 |
| oAβ42 + PNU vs. oAβ42 + NS | | 0.8579 | | 0.3842 to 1.332 | <0.0001 |
| oAβ42 + RJR vs. oAβ42 + NS | | -0.02219 | | -0.5348 to 0.4904 | >0.9999 |

**Supplementary Table 6. Statistical analysis of Ca^2+^ imaging in Fig. 2d.**

| **SST+ interneurons** | | | | | |
| --- | --- | --- | --- | --- | --- |
| **Normalized Total ΔF/F_min_** | | | | | |
|  | | sAβ42  sAβ42 + PNU  sAβ42 + RJR  sAβ42 + NS  oAβ42  oAβ42 + PNU  oAβ42 + RJR  oAβ42 + NS | | 1.000 ± 0.499 ΔF/F_min_  0.960 ± 0.344 ΔF/F_min_  1.286 ± 0.448 ΔF/F_min_  0.973 ± 0.345 ΔF/F_min_  0.604 ± 0.295 ΔF/F_min_  0.557 ± 0.271 ΔF/F_min_  0.994 ± 0.358 ΔF/F_min_  0.561 ± 0.249 ΔF/F_min_ | |
| ANOVA table | SS (Type III) | DF | MS | F (DFn, DFd) | P value |
| Interaction | 0.1032 | 3 | 0.03441 | F (3, 165) = 0.2529 | P=0.8592 |
| Row Factor | 5.984 | 1 | 5.984 | F (1, 165) = 43.98 | P<0.0001 |
| Column Factor | 4.458 | 3 | 1.486 | F (3, 165) = 10.92 | P<0.0001 |
| Residual | 22.45 | 165 | 0.1360 |  | |
| Tukey's multiple comparisons test | | Mean diff. | | 95.00% CI of diff. | Adjusted *P* Value |
| sAβ42 vs. sAβ42 + PNU | | 0.04027 | | -0.3043 to 0.3849 | >0.9999 |
| sAβ42 vs. sAβ42 + RJR | | -0.2864 | | -0.6041 to 0.03128 | 0.1104 |
| sAβ42 vs. sAβ42 + NS | | 0.02724 | | -0.3173 to 0.3718 | >0.9999 |
| sAβ42 vs. oAβ42 | | 0.3965 | | 0.07877 to 0.7141 | 0.0044 |
| sAβ42 vs. oAβ42 + PNU | | 0.4431 | | 0.1179 to 0.7684 | 0.0012 |
| sAβ42 vs. oAβ42 + RJR | | 0.006344 | | -0.3189 to 0.3316 | >0.9999 |
| sAβ42 vs. oAβ42 + NS | | 0.4390 | | 0.09440 to 0.7836 | 0.0033 |
| sAβ42 + PNU vs. sAβ42 + RJR | | -0.3267 | | -0.6798 to 0.02641 | 0.0922 |
| sAβ42 + PNU vs. sAβ42 + NS | | -0.01303 | | -0.3905 to 0.3644 | >0.9999 |
| sAβ42 + PNU vs. oAβ42 | | 0.3562 | | 0.003097 to 0.7093 | 0.0464 |
| sAβ42 + PNU vs. oAβ42 + PNU | | 0.4029 | | 0.04296 to 0.7628 | 0.0167 |
| sAβ42 + PNU vs. oAβ42 + RJR | | -0.03392 | | -0.3938 to 0.3260 | >0.9999 |
| sAβ42 + PNU vs. oAβ42 + NS | | 0.3987 | | 0.02124 to 0.7762 | 0.0303 |
| sAβ42 + RJR vs. sAβ42 +NS | | 0.3137 | | -0.03944 to 0.6667 | 0.1218 |
| sAβ42 + RJR vs. oAβ42 | | 0.6829 | | 0.3560 to 1.010 | <0.0001 |
| sAβ42 + RJR vs. oAβ42 + PNU | | 0.7295 | | 0.3953 to 1.064 | <0.0001 |
| sAβ42 + RJR vs. oAβ42 + RJR | | 0.2928 | | -0.04149 to 0.6270 | 0.1333 |
| sAβ42 + RJR vs. oAβ42 + NS | | 0.7254 | | 0.3723 to 1.078 | <0.0001 |
| sAβ42 + NS vs. oAβ42 | | 0.3692 | | 0.01613 to 0.7223 | 0.0334 |
| sAβ42 + NS vs. oAβ42 + PNU | | 0.4159 | | 0.05599 to 0.7758 | 0.0116 |
| sAβ42 + NS vs. oAβ42 + RJR | | -0.02089 | | -0.3808 to 0.3390 | >0.9999 |
| sAβ42 + NS vs. oAβ42 + NS | | 0.4117 | | 0.03427 to 0.7892 | 0.0220 |
| oAβ42 vs. oAβ42 + PNU | | 0.04667 | | -0.2876 to 0.3809 | 0.9999 |
| oAβ42 vs. oAβ42 + RJR | | -0.3901 | | -0.7244 to -0.05587 | 0.0103 |
| oAβ42 vs. oAβ42 + NS | | 0.04252 | | -0.3106 to 0.3956 | >0.9999 |
| oAβ42 + PNU vs. oAβ42 + RJR | | -0.4368 | | -0.7782 to -0.09535 | 0.0031 |
| oAβ42 + PNU vs. oAβ42 + NS | | -0.004149 | | -0.3641 to 0.3558 | >0.9999 |
| oAβ42 + RJR vs. oAβ42 + NS | | 0.4326 | | 0.07273 to 0.7925 | 0.0072 |

**Supplementary Table 7. Statistical analysis of c-Fos counting in hippocampal pyramidal cells in Fig. 3b.**

| **Female** | | | |
| --- | --- | --- | --- |
|  | WT + Saline  WT + PNU  WT + RJR  WT + PNU + RJR  5XFAD + Saline  5XFAD + PNU  5XFAD + RJR  5XFAD + PNU + RJR | 69.845 ± 27.943  91.705 ± 38.469  74.321 ± 36.753  71.036 ± 29.632  150.818 ± 56.931  121.873 ± 76.511  127.829 ± 59.897  79.067 ± 47.185 | |
| Fixed effects (type III) | P value | F (DFn, DFd) | |
| Genotype | <0.0001 | F (1, 280) = 47.12 | |
| Section | 0.0024 | F (2.069, 193.1) = 6.102 | |
| Genotype x Section | 0.0028 | F (2.069, 193.1) = 5.939 | |
| Tukey's multiple comparisons test | Mean diff. | 95.00% CI of diff. | Adjusted P Value |
| WT |  |  |  |
| Saline vs. PNU | -23.58 | -45.28 to -1.888 | 0.0295 |
| Saline vs. RJR | 3.554 | -16.01 to 23.11 | 0.9590 |
| Saline vs. PNU+RJR | -1.488 | -23.19 to 20.21 | 0.9975 |
| PNU vs. RJR | 24.46 | -1.677 to 50.60 | 0.0725 |
| PNU vs. PNU+RJR | 15.69 | -15.33 to 46.71 | 0.5072 |
| RJR vs. PNU+RJR | -3.890 | -17.41 to 9.632 | 0.8577 |
| 5XFAD |  |  |  |
| Saline vs. PNU | 25.63 | -31.71 to 82.97 | 0.6233 |
| Saline vs. RJR | 20.98 | -9.221 to 51.18 | 0.2577 |
| Saline vs. PNU+RJR | 80.71 | 36.29 to 125.1 | 0.0001 |
| PNU vs. RJR | -5.344 | -57.42 to 46.73 | 0.9926 |
| PNU vs. PNU+RJR | 29.44 | 2.300 to 56.59 | 0.0292 |
| RJR vs. PNU+RJR | 54.08 | 17.14 to 91.01 | 0.0019 |
| Saline |  |  |  |
| WT vs. 5XFAD | -80.97 | -101.8 to -60.10 | <0.0001 |
| PNU |  |  |  |
| WT vs. 5XFAD | -30.17 | -57.68 to -2.656 | 0.0321 |
| RJR |  |  |  |
| WT vs. 5XFAD | -53.51 | -74.91 to -32.10 | <0.0001 |
| PNU+RJR |  |  |  |
| WT vs. 5XFAD | -8.031 | -26.63 to 10.57 | 0.3918 |
| **Male** | | | |
|  | WT + Saline  WT + PNU  WT + RJR  WT + PNU + RJR  5XFAD + Saline  5XFAD + PNU  5XFAD + RJR  5XFAD + PNU + RJR | 65.166 ± 28.820  90.577 ± 29.214  83.423 ± 37.648  82.915 ± 48.144  143.918 ± 47.784  110.495 ± 34.132  131.236 ± 53.291  78.509 ± 43.314 | |
| Fixed effects (type III) | P value | F (DFn, DFd) | |
| Genotype | <0.0001 | F (1, 88) = 47.12 | |
| Section | 0.0013 | F (2.723, 176.1) = 5.796 | |
| Genotype x Section | <0.0001 | F (2.723, 176.1) = 12.81 | |
| Tukey's multiple comparisons test | Mean diff. | 95.00% CI of diff. | Adjusted P Value |
| WT |  |  |  |
| Saline vs. PNU | -23.28 | -41.89 to -4.670 | 0.0101 |
| Saline vs. RJR | -15.24 | -39.74 to 9.274 | 0.3452 |
| Saline vs. PNU+RJR | -17.75 | -49.61 to 14.12 | 0.4404 |
| PNU vs. RJR | 9.457 | -17.82 to 36.73 | 0.7790 |
| PNU vs. PNU+RJR | 3.777 | -28.83 to 36.39 | 0.9887 |
| RJR vs. PNU+RJR | -2.513 | -28.81 to 23.78 | 0.9937 |
| 5XFAD |  |  |  |
| Saline vs. PNU | 33.18 | 11.61 to 54.74 | 0.0013 |
| Saline vs. RJR | 22.93 | -14.28 to 60.14 | 0.3558 |
| Saline vs. PNU+RJR | 72.86 | 42.96 to 102.8 | <0.0001 |
| PNU vs. RJR | -21.87 | -48.29 to 4.555 | 0.1352 |
| PNU vs. PNU+RJR | 28.26 | 3.998 to 52.52 | 0.0175 |
| RJR vs. PNU+RJR | 45.18 | 12.21 to 78.14 | 0.0040 |
| Saline |  |  |  |
| WT vs. 5XFAD | -78.75 | -97.14 to -60.37 | <0.0001 |
| PNU |  |  |  |
| WT vs. 5XFAD | -19.92 | -35.00 to -4.842 | 0.0104 |
| RJR |  |  |  |
| WT vs. 5XFAD | -47.81 | -67.97 to -27.66 | <0.0001 |
| PNU+RJR |  |  |  |
| WT vs. 5XFAD | 4.406 | -18.23 to 27.04 | 0.6983 |

**Supplementary Table 8. Statistical analysis of c-Fos counting in hippocampal PV+ interneurons in Fig. 3c.**

| **Female** | | | |
| --- | --- | --- | --- |
|  | WT + Saline  WT + PNU  WT + RJR  WT + PNU + RJR  5XFAD + Saline  5XFAD + PNU  5XFAD + RJR  5XFAD + PNU + RJR | 3.256 ± 1.730  3.008 ± 2.573  1.898 ± 1.580  4.833 ± 2.944  0.788 ± 0.929  3.827 ± 2.703  1.627 ± 1.735  4.554 ± 2.889 | |
| Fixed effects (type III) | P value | F (DFn, DFd) | |
| Genotype | 0.0672 | F (1, 88) = 3.436 | |
| Section | <0.0001 | F (2.312, 131.8) = 23.36 | |
| Genotype x Section | 0.0017 | F (2.312, 131.8) = 6.135 | |
| Tukey's multiple comparisons test | Mean diff. | 95.00% CI of diff. | Adjusted P Value |
| WT |  |  |  |
| Saline vs. PNU | 0.01388 | -1.889 to 1.917 | >0.9999 |
| Saline vs. RJR | 1.315 | -0.5405 to 3.170 | 0.2165 |
| Saline vs. PNU+RJR | -0.9393 | -3.511 to 1.632 | 0.7117 |
| PNU vs. RJR | 0.3299 | -0.9866 to 1.646 | 0.8867 |
| PNU vs. PNU+RJR | -2.642 | -5.111 to -0.1730 | 0.0343 |
| RJR vs. PNU+RJR | -3.339 | -4.656 to -2.021 | <0.0001 |
| 5XFAD |  |  |  |
| Saline vs. PNU | -3.151 | -4.533 to -1.769 | <0.0001 |
| Saline vs. RJR | -0.9984 | -2.065 to 0.06850 | 0.0735 |
| Saline vs. PNU+RJR | -4.092 | -5.662 to -2.522 | <0.0001 |
| PNU vs. RJR | 2.241 | 0.9280 to 3.554 | 0.0003 |
| PNU vs. PNU+RJR | -0.6088 | -2.352 to 1.134 | 0.7857 |
| RJR vs. PNU+RJR | -2.875 | -4.177 to -1.572 | <0.0001 |
| Saline |  |  |  |
| WT vs. 5XFAD | 2.468 | 1.672 to 3.264 | <0.0001 |
| PNU |  |  |  |
| WT vs. 5XFAD | -0.8187 | -2.191 to 0.5531 | 0.2354 |
| RJR |  |  |  |
| WT vs. 5XFAD | 0.2716 | -0.5180 to 1.061 | 0.4942 |
| PNU+RJR |  |  |  |
| WT vs. 5XFAD | 0.2792 | -1.321 to 1.880 | 0.7256 |
| **Male** | | | |
|  | WT + Saline  WT + PNU  WT + RJR  WT + PNU + RJR  5XFAD + Saline  5XFAD + PNU  5XFAD + RJR  5XFAD + PNU + RJR | 2.513 ± 1.294  2.751 ± 1.591  1.977 ± 1.495  3.664 ± 2.559  0.733 ± 0.624  3.487 ± 2.980  1.067 ± 1.580  4.391 ± 3.143 | |
| Fixed effects (type III) | P value | F (DFn, DFd) | |
| Genotype | 0.2528 | F (1, 266) = 1.313 | |
| Section | <0.0001 | F (2.110, 187.1) = 21.23 | |
| Genotype x Section | 0.0054 | F (2.110, 187.1) = 5.214 | |
| Tukey's multiple comparisons test | Mean diff. | 95.00% CI of diff. | Adjusted P Value |
| WT |  |  |  |
| Saline vs. PNU | -0.2465 | -1.816 to 1.323 | 0.9708 |
| Saline vs. RJR | 0.6232 | -0.3278 to 1.574 | 0.2873 |
| Saline vs. PNU+RJR | -1.085 | -2.757 to 0.5883 | 0.2959 |
| PNU vs. RJR | 1.167 | 0.2022 to 2.131 | 0.0138 |
| PNU vs. PNU+RJR | -0.7766 | -2.477 to 0.9238 | 0.6000 |
| RJR vs. PNU+RJR | -1.779 | -3.343 to -0.2149 | 0.0218 |
| 5XFAD |  |  |  |
| Saline vs. PNU | -3.241 | -5.156 to -1.326 | 0.0005 |
| Saline vs. RJR | -0.2224 | -1.021 to 0.5759 | 0.8721 |
| Saline vs. PNU+RJR | -4.131 | -5.909 to -2.352 | <0.0001 |
| PNU vs. RJR | 2.772 | 1.454 to 4.090 | <0.0001 |
| PNU vs. PNU+RJR | -0.5985 | -2.772 to 1.575 | 0.8783 |
| RJR vs. PNU+RJR | -3.420 | -5.025 to -1.816 | <0.0001 |
| Saline |  |  |  |
| WT vs. 5XFAD | 1.779 | 1.187 to 2.372 | <0.0001 |
| PNU |  |  |  |
| WT vs. 5XFAD | -0.7365 | -1.864 to 0.3913 | 0.1965 |
| RJR |  |  |  |
| WT vs. 5XFAD | 0.9094 | 0.2000 to 1.619 | 0.0128 |
| PNU+RJR |  |  |  |
| WT vs. 5XFAD | -0.7273 | -2.079 to 0.6245 | 0.2866 |

**Supplementary Table 9. Statistical analysis of c-Fos counting in hippocampal SST+ interneurons in Fig. 3d.**

| **Female** | | | | |
| --- | --- | --- | --- | --- |
|  | WT + Saline  WT + PNU  WT + RJR  WT + PNU + RJR  5XFAD + Saline  5XFAD + PNU  5XFAD + RJR  5XFAD + PNU + RJR | 1.868 ± 1.527  1.418 ± 0.687  1.725 ± 0.939  2.124 ± 1.333  0.781 ± 0.854  0.911 ± 0.934  2.876 ± 2.408  2.219 ± 1.498 | | |
| Fixed effects (type III) | P value | F (DFn, DFd) | | |
| Genotype | 0.6570 | F (1, 89) = 0.1986 | | |
| Section | <0.0001 | F (2.625, 121.6) = 10.77 | | |
| Genotype x Section | 0.0002 | F (2.625, 121.6) = 7.943 | | |
| Tukey's multiple comparisons test | Mean diff. | 95.00% CI of diff. | Adjusted P Value | |
| WT |  |  |  | |
| Saline vs. PNU | 0.08190 | -0.6761 to 0.8399 | 0.9880 | |
| Saline vs. RJR | 0.2914 | -0.7407 to 1.324 | 0.8544 | |
| Saline vs. PNU+RJR | -0.4847 | -1.888 to 0.9181 | 0.7495 | |
| PNU vs. RJR | -0.4226 | -1.194 to 0.3487 | 0.4188 | |
| PNU vs. PNU+RJR | -1.021 | -2.428 to 0.3855 | 0.1911 | |
| RJR vs. PNU+RJR | -0.6521 | -1.962 to 0.6577 | 0.5031 | |
| 5XFAD |  |  |  | |
| Saline vs. PNU | -0.2318 | -0.7893 to 0.3257 | 0.6740 | |
| Saline vs. RJR | -2.364 | -3.771 to -0.9579 | 0.0006 | |
| Saline vs. PNU+RJR | -1.486 | -2.281 to -0.6912 | 0.0002 | |
| PNU vs. RJR | -1.491 | -2.485 to -0.4966 | 0.0022 | |
| PNU vs. PNU+RJR | -1.297 | -2.238 to -0.3554 | 0.0042 | |
| RJR vs. PNU+RJR | 0.8887 | -0.6802 to 2.458 | 0.3994 | |
| Saline |  |  |  | |
| WT vs. 5XFAD | 1.087 | 0.3811 to 1.794 | 0.0037 | |
| PNU |  |  |  | |
| WT vs. 5XFAD | 0.5070 | 0.07205 to 0.9420 | 0.0234 | |
| RJR |  |  |  | |
| WT vs. 5XFAD | -1.151 | -2.139 to -0.1632 | 0.0237 | |
| PNU+RJR |  |  |  | |
| WT vs. 5XFAD | -0.09475 | -0.8809 to 0.6914 | 0.8094 | |
| **Male** | | | | |
|  | WT + Saline  WT + PNU  WT + RJR  WT + PNU + RJR  5XFAD + Saline  5XFAD + PNU  5XFAD + RJR  5XFAD + PNU + RJR | 1.699 ± 1.238  2.593 ± 1.741  1.776 ± 1.143  2.687 ± 1.386  0.511 ± 0.669  1.001 ± 0.903  2.650 ± 2.216  1.967 ± 1.413 | | |
| Fixed effects (type III) | P value | F (DFn, DFd) | | |
| Genotype | 0.0012 | F (1, 86) = 11.28 | | |
| Section | <0.0001 | F (2.480, 131.4) = 10.15 | | |
| Genotype x Section | <0.0001 | F (2.480, 131.4) = 9.251 | | |
| Tukey's multiple comparisons test | Mean diff. | 95.00% CI of diff. | | Adjusted *P* Value |
| WT |  |  | |  |
| Saline vs. PNU | -0.6092 | -1.853 to 0.6344 | | 0.5206 |
| Saline vs. RJR | 0.01845 | -0.9760 to 1.013 | | >0.9999 |
| Saline vs. PNU+RJR | -0.9301 | -1.901 to 0.04050 | | 0.0634 |
| PNU vs. RJR | 0.9116 | -0.7084 to 2.532 | | 0.3859 |
| PNU vs. PNU+RJR | 0.04961 | -1.265 to 1.364 | | 0.9996 |
| RJR vs. PNU+RJR | -0.9559 | -2.158 to 0.2457 | | 0.1454 |
| 5XFAD |  |  | |  |
| Saline vs. PNU | -0.3339 | -0.7896 to 0.1217 | | 0.2145 |
| Saline vs. RJR | -2.518 | -3.814 to -1.222 | | <0.0001 |
| Saline vs. PNU+RJR | -1.420 | -2.237 to -0.6040 | | 0.0004 |
| PNU vs. RJR | -1.704 | -2.770 to -0.6375 | | 0.0008 |
| PNU vs. PNU+RJR | -0.8153 | -1.531 to -0.09946 | | 0.0206 |
| RJR vs. PNU+RJR | 0.1945 | -1.036 to 1.425 | | 0.9714 |
| Saline |  |  | |  |
| WT vs. 5XFAD | 1.188 | 0.6284 to 1.748 | | 0.0001 |
| PNU |  |  | |  |
| WT vs. 5XFAD | 1.592 | 0.8010 to 2.383 | | 0.0003 |
| RJR |  |  | |  |
| WT vs. 5XFAD | -0.8740 | -1.770 to 0.02214 | | 0.0557 |
| PNU+RJR |  |  | |  |
| WT vs. 5XFAD | 0.7206 | 0.007995 to 1.433 | | 0.0476 |

**Supplementary Table 10. Statistical analysis of LFP data in Fig. 4e-g.**

| **Female** | | | | | |
| --- | --- | --- | --- | --- | --- |
| Theta power | | Slow gamma power | | Fast gamma power | |
| WT PRE  WT POST | 1.000 ± 1.199  1.696 ± 1.552 | WT PRE  WT POST | 1.000 ± 1.223  1.624 ± 1.420 | WT PRE  WT POST | 1.000 ± 1.086 1.601 ± 1.372 |
| Paired t test | | Paired t test | | Paired t test | |
| P value | 0.0029 | P value | 0.0031 | P value | 0.0009 |
| t, df | t=3.084, df=73 | t, df | t=3.057, df=73 | t, df | t=3.467, df=73 |
| 5XFAD PRE  5XFAD POST | 2.195 ± 6.108  0.225 ± 0.210 | 5XFAD PRE  5XFAD POST | 3.145 ± 7.767  0.480 ± 0.342 | 5XFAD PRE  5XFAD POST | 3.059 ± 7.262  0.493 ± 0.284 |
| Paired t test | | Paired t test | | Paired t test | |
| P value | 0.0015 | P value | 0.0007 | P value | 0.0005 |
| t, df | t=3.260, df=100 | t, df | t=3.494, df=100 | t, df | t=3.609, df=100 |
| WT PRE vs 5XFAD PRE | | WT PRE vs 5XFAD PRE | | WT PRE vs 5XFAD PRE | |
| Šídák's multiple comparisons test | | Šídák's multiple comparisons test | | Šídák's multiple comparisons test | |
| p=0.0445 (t=2.292, df=346) | | p=0.0022 (t=3.285, df=346) | | p=0.0017 (t=3.373, df=346) | |
| **Male** | | | | | |
| Theta power | | Slow gamma power | | Fast gamma power | |
| WT PRE  WT POST | 1.000 ± 1.270  1.926 ± 2.849 | WT PRE  WT POST | 1.000 ± 1.124 2.049 ± 2.413 | WT PRE  WT POST | 1.000 ± 1.051 2.467 ± 2.486 |
| Paired t test | | Paired t test | | Paired t test | |
| P value | 0.0001 | P value | < 0.0001 | P value | < 0.0001 |
| t, df | t=3.926, df=124 | t, df | t=5.816, df=124 | t, df | t=8.187, df=124 |
| 5XFAD PRE  5XFAD POST | 2.168 ± 3.446  0.539 ± 1.019 | 5XFAD PRE  5XFAD POST | 2.095 ± 3.080  0.690 ± 0.940 | 5XFAD PRE  5XFAD POST | 2.055 ± 2.724  0.770 ± 0.855 |
| Paired t test | | Paired t test | | Paired t test | |
| P value | < 0.0001 | P value | < 0.0001 | P value | < 0.0001 |
| t, df | t=4.906, df=105 | t, df | t=4.684, df=105 | t, df | t=4.857, df=105 |
| WT PRE vs 5XFAD PRE | | WT PRE vs 5XFAD PRE | | WT PRE vs 5XFAD PRE | |
| Šídák's multiple comparisons test | | Šídák's multiple comparisons test | | Šídák's multiple comparisons test | |
| p=0.0004 (t=3.739, df=458) | | p<0.0001 (t=4.002, df=458) | | P=0.0001 (t=4.078, df=458) | |

**Supplementary Table 11. Statistical analysis of LFP data in Fig. 4h-i.**

| **Female** | | | |
| --- | --- | --- | --- |
| Theta-slow gamma PAC | | Theta-fast gamma PAC | |
| WT PRE  WT POST | 0.036 ± 0.027  0.067 ± 0.044 | WT PRE  WT POST | 0.059 ± 0.087 0.076 ± 0.070 |
| Paired t test | | Paired t test | |
| P value | < 0.0001 | P value | 0.0004 |
| t, df | t=6.667, df=73 | t, df | t=4.128, df=73 |
| 5XFAD PRE  5XFAD POST | 0.055 ± 0.021 0.042 ± 0.018 | 5XFAD PRE  5XFAD POST | 0.028 ± 0.015 0.028 ± 0.022 |
| Paired t test | | Paired t test | |
| P value | *p* < 0.0001 | P value | 0.9556 |
| t, df | t=6.032, df=100 | t, df | t=0.056, df=100 |
| **Male** | | | |
| Theta-slow gamma PAC | | Theta-fast gamma PAC | |
| WT PRE  WT POST | 0.041 ± 0.032 0.070 ± 0.053 | WT PRE  WT POST | 0.036 ± 0.042 0.061 ± 0.061 |
| Paired t test | | Paired t test | |
| P value | < 0.0001 | P value | < 0.0001 |
| t, df | t=7.153, df=124 | t, df | t=9.003, df=124 |
| 5XFAD PRE  5XFAD POST | 0.060 ± 0.067 0.043 ± 0.047 | 5XFAD PRE  5XFAD POST | 0.041 ± 0.046 0.030 ± 0.039 |
| Paired t test | | Paired t test | |
| P value | 0.0157 | P value | 0.0180 |
| t, df | t=2.455, df=104 | t, df | t=2.404, df=104 |

**Supplementary Table 12. Statistical analysis of LFP theta power data in Fig. 5a.**

| **Female** | | | |
| --- | --- | --- | --- |
|  | WT + Saline  WT + PNU  WT + RJR  WT + PNU + RJR  5XFAD + Saline  5XFAD + PNU  5XFAD + RJR  5XFAD + PNU + RJR | 1.000 ± 0.915  0.490 ± 0.950  0.566 ± 0.600  0.457 ± 1.209  0.151 ± 0.148  0.138 ± 0.234  0.127 ± 0.174  0.518 ± 0.834 | |
| Fixed effects (type III) | P value | F (DFn, DFd) | |
| Genotype | <0.0001 | F (1, 532) = 30.14 | |
| Electrode | 0.0323 | F (3, 532) = 2.949 | |
| Genotype x Electrode | <0.0001 | F (3, 532) = 10.40 | |
| Tukey's multiple comparisons test | Mean diff. | 95.00% CI of diff. | Adjusted P Value |
| WT |  |  |  |
| Saline vs. PNU | 0.5098 | 0.1003 to 0.9193 | 0.0077 |
| Saline vs. RJR | 0.4344 | 0.06440 to 0.8043 | 0.0138 |
| Saline vs. PNU+RJR | 0.5429 | 0.2347 to 0.8510 | <0.0001 |
| PNU vs. RJR | -0.07542 | -0.5191 to 0.3683 | 0.9719 |
| PNU vs. PNU+RJR | 0.03306 | -0.3606 to 0.4267 | 0.9964 |
| RJR vs. PNU+RJR | 0.1085 | -0.2438 to 0.4608 | 0.8574 |
| 5XFAD |  |  |  |
| Saline vs. PNU | 0.01254 | -0.3304 to 0.3555 | 0.9997 |
| Saline vs. RJR | 0.02343 | -0.3462 to 0.3931 | 0.9984 |
| Saline vs. PNU+RJR | -0.3674 | -0.6543 to -0.08049 | 0.0057 |
| PNU vs. RJR | 0.01089 | -0.4079 to 0.4296 | 0.9999 |
| PNU vs. PNU+RJR | -0.3799 | -0.7278 to -0.03208 | 0.0260 |
| RJR vs. PNU+RJR | -0.3908 | -0.7650 to -0.01657 | 0.0368 |
| Saline |  |  |  |
| WT vs. 5XFAD | 0.8494 | 0.6165 to 1.082 | <0.0001 |
| PNU |  |  |  |
| WT vs. 5XFAD | 0.3521 | 0.01810 to 0.6861 | 0.0388 |
| RJR |  |  |  |
| WT vs. 5XFAD | 0.4384 | 0.1148 to 0.7620 | 0.0080 |
| PNU+RJR |  |  |  |
| WT vs. 5XFAD | -0.06087 | -0.2817 to 0.1600 | 0.5884 |
| **Male** | | | |
|  | WT + Saline  WT + PNU  WT + RJR  WT + PNU + RJR  5XFAD + Saline  5XFAD + PNU  5XFAD + RJR  5XFAD + PNU + RJR | 1.000 ± 1.479  0.422 ± 0.365  0.489 ± 0.256  0.609 ± 1.648  0.280 ± 0.529  0.123 ± 0.137  0.090 ± 0.107  0.684 ± 0.986 | |
| Fixed effects (type III) | P value | F (DFn, DFd) | |
| Genotype | 0.0004 | F (1, 570) = 12.50 | |
| Electrode | 0.0024 | F (3, 570) = 4.872 | |
| Genotype x Electrode | 0.0030 | F (3, 570) = 4.696 | |
| Tukey's multiple comparisons test | Mean diff. | 95.00% CI of diff. | Adjusted *P* Value |
| WT |  |  |  |
| Saline vs. PNU | 0.5778 | 0.1486 to 1.007 | 0.0031 |
| Saline vs. RJR | 0.5111 | 0.06496 to 0.9573 | 0.0173 |
| Saline vs. PNU+RJR | 0.3909 | 0.01470 to 0.7671 | 0.0382 |
| PNU vs. RJR | -0.06664 | -0.5842 to 0.4509 | 0.9874 |
| PNU vs. PNU+RJR | -0.1869 | -0.6455 to 0.2718 | 0.7201 |
| RJR vs. PNU+RJR | -0.1202 | -0.5948 to 0.3544 | 0.9146 |
| 5XFAD |  |  |  |
| Saline vs. PNU | 0.1567 | -0.3613 to 0.6747 | 0.8638 |
| Saline vs. RJR | 0.1893 | -0.3185 to 0.6970 | 0.7720 |
| Saline vs. PNU+RJR | -0.4040 | -0.8031 to -0.004831 | 0.0460 |
| PNU vs. RJR | 0.03258 | -0.5920 to 0.6571 | 0.9991 |
| PNU vs. PNU+RJR | -0.5606 | -1.101 to -0.02066 | 0.0384 |
| RJR vs. PNU+RJR | -0.5932 | -1.123 to -0.06308 | 0.0212 |
| Saline |  |  |  |
| WT vs. 5XFAD | 0.7203 | 0.4500 to 0.9906 | <0.0001 |
| PNU |  |  |  |
| WT vs. 5XFAD | 0.2992 | -0.1366 to 0.7350 | 0.1780 |
| RJR |  |  |  |
| WT vs. 5XFAD | 0.3984 | -0.04022 to 0.8371 | 0.0750 |
| PNU+RJR |  |  |  |
| WT vs. 5XFAD | -0.07455 | -0.3935 to 0.2444 | 0.6464 |

**Supplementary Table 13. Statistical analysis of LFP slow gamma power data in Fig. 5b.**

| **Female** | | | |
| --- | --- | --- | --- |
|  | WT + Saline  WT + PNU  WT + RJR  WT + PNU + RJR  5XFAD + Saline  5XFAD + PNU  5XFAD + RJR  5XFAD + PNU + RJR | 1.000 ± 0.874  0.601 ± 0.963  1.452 ± 1.705  0.998 ± 2.758  0.268 ± 0.167  0.305 ± 0.419  0.214 ± 0.297  1.058 ± 1.633 | |
| Fixed effects (type III) | P value | F (DFn, DFd) | |
| Genotype | <0.0001 | F (1, 268) = 18.24 | |
| Electrode | 0.0240 | F (3, 264) = 3.197 | |
| Genotype x Electrode | <0.0001 | F (3, 264) = 10.11 | |
| Tukey's multiple comparisons test | Mean diff. | 95.00% CI of diff. | Adjusted P Value |
| WT |  |  |  |
| Saline vs. PNU | 0.1377 | -0.5836 to 0.8590 | 0.9605 |
| Saline vs. RJR | -0.7994 | -1.467 to -0.1323 | 0.0115 |
| Saline vs. PNU+RJR | 0.08354 | -0.4661 to 0.6332 | 0.9794 |
| PNU vs. RJR | -0.9371 | -1.668 to -0.2060 | 0.0057 |
| PNU vs. PNU+RJR | -0.05416 | -0.7321 to 0.6237 | 0.9969 |
| RJR vs. PNU+RJR | 0.8830 | 0.2784 to 1.488 | 0.0011 |
| 5XFAD |  |  |  |
| Saline vs. PNU | 0.03497 | -0.5617 to 0.6317 | 0.9988 |
| Saline vs. RJR | 0.2035 | -0.4467 to 0.8536 | 0.8501 |
| Saline vs. PNU+RJR | -0.7432 | -1.231 to -0.2554 | 0.0006 |
| PNU vs. RJR | 0.1685 | -0.5122 to 0.8492 | 0.9190 |
| PNU vs. PNU+RJR | -0.7782 | -1.365 to -0.1916 | 0.0039 |
| RJR vs. PNU+RJR | -0.9467 | -1.583 to -0.3099 | 0.0009 |
| Saline |  |  |  |
| WT vs. 5XFAD | 0.7555 | 0.2897 to 1.221 | 0.0015 |
| PNU |  |  |  |
| WT vs. 5XFAD | 0.6528 | 0.01591 to 1.290 | 0.0446 |
| RJR |  |  |  |
| WT vs. 5XFAD | 1.758 | 1.137 to 2.380 | <0.0001 |
| PNU+RJR |  |  |  |
| WT vs. 5XFAD | -0.07124 | -0.5170 to 0.3745 | 0.7537 |
| **Male** | | | |
|  | WT + Saline  WT + PNU  WT + RJR  WT + PNU + RJR  5XFAD + Saline  5XFAD + PNU  5XFAD + RJR  5XFAD + PNU + RJR | 1.000 ± 1.079  0.879 ± 0.716  0.996 ± 0.461  1.149 ± 2.906  0.304 ± 0.414  0.272 ± 0.157  0.142 ± 0.122  1.300 ± 2.375 | |
| Fixed effects (type III) | P value | F (DFn, DFd) | |
| Genotype | 0.0004 | F (1, 260) = 12.70 | |
| Electrode | 0.0005 | F (3, 310) = 6.133 | |
| Genotype x Electrode | 0.0225 | F (3, 310) = 3.237 | |
| Tukey's multiple comparisons test | Mean diff. | 95.00% CI of diff. | Adjusted *P* Value |
| WT |  |  |  |
| Saline vs. PNU | 0.1095 | -0.5254 to 0.7445 | 0.9704 |
| Saline vs. RJR | -0.005204 | -0.6656 to 0.6552 | >0.9999 |
| Saline vs. PNU+RJR | -0.1470 | -0.7031 to 0.4092 | 0.9037 |
| PNU vs. RJR | -0.1147 | -0.8790 to 0.6496 | 0.9802 |
| PNU vs. PNU+RJR | -0.2565 | -0.9344 to 0.4215 | 0.7626 |
| RJR vs. PNU+RJR | -0.1418 | -0.8437 to 0.5602 | 0.9539 |
| 5XFAD |  |  |  |
| Saline vs. PNU | 0.03240 | -0.7349 to 0.7997 | 0.9995 |
| Saline vs. RJR | 0.1664 | -0.5856 to 0.9184 | 0.9405 |
| Saline vs. PNU+RJR | -0.9977 | -1.588 to -0.4077 | 0.0001 |
| PNU vs. RJR | 0.1340 | -0.7904 to 1.058 | 0.9821 |
| PNU vs. PNU+RJR | -1.030 | -1.829 to -0.2311 | 0.0054 |
| RJR vs. PNU+RJR | -1.164 | -1.948 to -0.3797 | 0.0009 |
| Saline |  |  |  |
| WT vs. 5XFAD | 0.6978 | 0.2971 to 1.099 | 0.0007 |
| PNU |  |  |  |
| WT vs. 5XFAD | 0.6207 | -0.02533 to 1.267 | 0.0597 |
| RJR |  |  |  |
| WT vs. 5XFAD | 0.8694 | 0.2191 to 1.520 | 0.0089 |
| PNU+RJR |  |  |  |
| WT vs. 5XFAD | -0.1529 | -0.6258 to 0.3200 | 0.5257 |

**Supplementary Table 14. Statistical analysis of LFP fast gamma power data in Fig. 5c.**

| **Female** | | | |
| --- | --- | --- | --- |
|  | WT + Saline  WT + PNU  WT + RJR  WT + PNU + RJR  5XFAD + Saline  5XFAD + PNU  5XFAD + RJR  5XFAD + PNU + RJR | 1.000 ± 0.853  0.532 ± 0.746  1.455 ± 1.660  1.052 ± 2.620  0.297 ± 0.162  0.269 ± 0.392  0.198 ± 0.240  1.253 ± 1.919 | |
| Fixed effects (type III) | P value | F (DFn, DFd) | |
| Genotype | 0.0003 | F (1, 532) = 13.01 | |
| Electrode | 0.0005 | F (3, 532) = 5.948 | |
| Genotype x Electrode | 0.0010 | F (3, 532) = 5.522 | |
| Tukey's multiple comparisons test | Mean diff. | 95.00% CI of diff. | Adjusted P Value |
| WT |  |  |  |
| Saline vs. PNU | 0.4679 | -0.3300 to 1.266 | 0.4314 |
| Saline vs. RJR | -0.4552 | -1.176 to 0.2656 | 0.3640 |
| Saline vs. PNU+RJR | -0.05159 | -0.6520 to 0.5488 | 0.9962 |
| PNU vs. RJR | -0.9232 | -1.788 to -0.05858 | 0.0311 |
| PNU vs. PNU+RJR | -0.5195 | -1.287 to 0.2475 | 0.3012 |
| RJR vs. PNU+RJR | 0.4037 | -0.2828 to 1.090 | 0.4289 |
| 5XFAD |  |  |  |
| Saline vs. PNU | 0.02842 | -0.6398 to 0.6966 | 0.9995 |
| Saline vs. RJR | 0.09868 | -0.6216 to 0.8190 | 0.9849 |
| Saline vs. PNU+RJR | -0.9564 | -1.515 to -0.3974 | <0.0001 |
| PNU vs. RJR | 0.07027 | -0.7456 to 0.8862 | 0.9961 |
| PNU vs. PNU+RJR | -0.9848 | -1.663 to -0.3071 | 0.0011 |
| RJR vs. PNU+RJR | -1.055 | -1.784 to -0.3259 | 0.0012 |
| Saline |  |  |  |
| WT vs. 5XFAD | 0.7031 | 0.2494 to 1.157 | 0.0024 |
| PNU |  |  |  |
| WT vs. 5XFAD | 0.2635 | -0.3873 to 0.9143 | 0.4267 |
| RJR |  |  |  |
| WT vs. 5XFAD | 1.257 | 0.6265 to 1.887 | 0.0001 |
| PNU+RJR |  |  |  |
| WT vs. 5XFAD | -0.2018 | -0.6321 to 0.2285 | 0.3574 |
| **Male** | | | |
|  | WT + Saline  WT + PNU  WT + RJR  WT + PNU + RJR  5XFAD + Saline  5XFAD + PNU  5XFAD + RJR  5XFAD + PNU + RJR | 1.000 ± 1.008  0.667 ± 0.595  0.794 ± 0.380  0.904 ± 2.040  0.312 ± 0.347  0.263 ± 0.195  0.142 ± 0.139  1.261 ± 1.763 | |
| Fixed effects (type III) | P value | F (DFn, DFd) | |
| Genotype | 0.0011 | F (1, 570) = 10.80 | |
| Electrode | <0.0001 | F (3, 570) = 8.428 | |
| Genotype x Electrode | <0.0001 | F (3, 570) = 7.248 | |
| Tukey's multiple comparisons test | Mean diff. | 95.00% CI of diff. | Adjusted *P* Value |
| WT |  |  |  |
| Saline vs. PNU | 0.3329 | -0.1438 to 0.8095 | 0.2747 |
| Saline vs. RJR | 0.2064 | -0.2892 to 0.7019 | 0.7062 |
| Saline vs. PNU+RJR | 0.09575 | -0.3221 to 0.5136 | 0.9350 |
| PNU vs. RJR | -0.1265 | -0.7014 to 0.4483 | 0.9418 |
| PNU vs. PNU+RJR | -0.2371 | -0.7465 to 0.2723 | 0.6274 |
| RJR vs. PNU+RJR | -0.1106 | -0.6377 to 0.4165 | 0.9490 |
| 5XFAD |  |  |  |
| Saline vs. PNU | 0.04936 | -0.5260 to 0.6247 | 0.9962 |
| Saline vs. RJR | 0.1708 | -0.3931 to 0.7347 | 0.8634 |
| Saline vs. PNU+RJR | -0.9487 | -1.392 to -0.5054 | <0.0001 |
| PNU vs. RJR | 0.1214 | -0.5722 to 0.8151 | 0.9694 |
| PNU vs. PNU+RJR | -0.9980 | -1.598 to -0.3983 | 0.0001 |
| RJR vs. PNU+RJR | -1.119 | -1.708 to -0.5307 | <0.0001 |
| Saline |  |  |  |
| WT vs. 5XFAD | 0.6877 | 0.3875 to 0.9879 | <0.0001 |
| PNU |  |  |  |
| WT vs. 5XFAD | 0.4041 | -0.07985 to 0.8881 | 0.1015 |
| RJR |  |  |  |
| WT vs. 5XFAD | 0.6521 | 0.1649 to 1.139 | 0.0088 |
| PNU+RJR |  |  |  |
| WT vs. 5XFAD | -0.3567 | -0.7110 to -0.002441 | 0.0484 |

**Supplementary Table 15. Statistical analysis of LFP theta-slow gamma PAC data in Fig. 5d.**

| **Female** | | | |
| --- | --- | --- | --- |
|  | WT + Saline  WT + PNU  WT + RJR  WT + PNU + RJR  5XFAD + Saline  5XFAD + PNU  5XFAD + RJR  5XFAD + PNU + RJR | 0.067 ± 0.044  0.125 ± 0.042  0.072 ± 0.034  0.100 ± 0.061  0.042 ± 0.018  0.072 ± 0.053  0.100 ± 0.062  0.120 ± 0.040 | |
| Fixed effects (type III) | P value | F (DFn, DFd) | |
| Genotype | 0.0789 | F (1, 546) = 3.100 | |
| Electrode | <0.0001 | F (3, 546) = 48.48 | |
| Genotype x Electrode | <0.0001 | F (3, 546) = 19.41 | |
| Tukey's multiple comparisons test | Mean diff. | 95.00% CI of diff. | Adjusted P Value |
| WT |  |  |  |
| Saline vs. PNU | -0.05811 | -0.08203 to -0.03419 | <0.0001 |
| Saline vs. RJR | -0.005962 | -0.02757 to 0.01565 | 0.8927 |
| Saline vs. PNU+RJR | -0.03313 | -0.05112 to -0.01513 | <0.0001 |
| PNU vs. RJR | 0.05215 | 0.02624 to 0.07807 | <0.0001 |
| PNU vs. PNU+RJR | 0.02499 | 0.001997 to 0.04798 | 0.0270 |
| RJR vs. PNU+RJR | -0.02716 | -0.04774 to -0.006587 | 0.0040 |
| 5XFAD |  |  |  |
| Saline vs. PNU | -0.03056 | -0.05046 to -0.01066 | 0.0005 |
| Saline vs. RJR | -0.05830 | -0.07920 to -0.03741 | <0.0001 |
| Saline vs. PNU+RJR | -0.07795 | -0.09432 to -0.06158 | <0.0001 |
| PNU vs. RJR | -0.02774 | -0.05148 to -0.004002 | 0.0144 |
| PNU vs. PNU+RJR | -0.04739 | -0.06726 to -0.02752 | <0.0001 |
| RJR vs. PNU+RJR | -0.01965 | -0.04051 to 0.001218 | 0.0733 |
| Saline |  |  |  |
| WT vs. 5XFAD | 0.02470 | 0.01110 to 0.03830 | 0.0004 |
| PNU |  |  |  |
| WT vs. 5XFAD | 0.05226 | 0.03282 to 0.07169 | <0.0001 |
| RJR |  |  |  |
| WT vs. 5XFAD | -0.02764 | -0.04608 to -0.009195 | 0.0034 |
| PNU+RJR |  |  |  |
| WT vs. 5XFAD | -0.02012 | -0.03273 to -0.007517 | 0.0018 |
| **Male** | | | |
|  | WT + Saline  WT + PNU  WT + RJR  WT + PNU + RJR  5XFAD + Saline  5XFAD + PNU  5XFAD + RJR  5XFAD + PNU + RJR | 0.070 ± 0.053  0.126 ± 0.055  0.097 ± 0.042  0.108 ± 0.082  0.043 ± 0.047  0.122 ± 0.054  0.107 ± 0.032  0.071 ± 0.053 | |
| Fixed effects (type III) | P value | F (DFn, DFd) | |
| Genotype | 0.0045 | F (1, 574) = 8.128 | |
| Electrode | <0.0001 | F (3, 574) = 38.17 | |
| Genotype x Electrode | 0.0059 | F (3, 574) = 4.209 | |
| Tukey's multiple comparisons test | Mean diff. | 95.00% CI of diff. | Adjusted *P* Value |
| WT |  |  |  |
| Saline vs. PNU | -0.05610 | -0.07907 to -0.03314 | <0.0001 |
| Saline vs. RJR | -0.02726 | -0.05113 to -0.003381 | 0.0178 |
| Saline vs. PNU+RJR | -0.03771 | -0.05784 to -0.01757 | <0.0001 |
| PNU vs. RJR | 0.02885 | 0.001149 to 0.05654 | 0.0375 |
| PNU vs. PNU+RJR | 0.01840 | -0.006146 to 0.04294 | 0.2161 |
| RJR vs. PNU+RJR | -0.01045 | -0.03585 to 0.01495 | 0.7140 |
| 5XFAD |  |  |  |
| Saline vs. PNU | -0.07905 | -0.1057 to -0.05238 | <0.0001 |
| Saline vs. RJR | -0.06323 | -0.09040 to -0.03606 | <0.0001 |
| Saline vs. PNU+RJR | -0.02743 | -0.04879 to -0.006075 | 0.0055 |
| PNU vs. RJR | 0.01582 | -0.01673 to 0.04837 | 0.5939 |
| PNU vs. PNU+RJR | 0.05162 | 0.02373 to 0.07950 | <0.0001 |
| RJR vs. PNU+RJR | 0.03580 | 0.007430 to 0.06417 | 0.0067 |
| Saline |  |  |  |
| WT vs. 5XFAD | 0.02650 | 0.01204 to 0.04097 | 0.0003 |
| PNU |  |  |  |
| WT vs. 5XFAD | 0.003558 | -0.01904 to 0.02615 | 0.7572 |
| RJR |  |  |  |
| WT vs. 5XFAD | -0.009470 | -0.03294 to 0.01400 | 0.4285 |
| PNU+RJR |  |  |  |
| WT vs. 5XFAD | 0.03678 | 0.01971 to 0.05385 | <0.0001 |

**Supplementary Table 16. Statistical analysis of LFP theta-fast gamma PAC data in Fig. 5e.**

| **Female** | | | | |
| --- | --- | --- | --- | --- |
|  | WT + Saline  WT + PNU  WT + RJR  WT + PNU + RJR  5XFAD + Saline  5XFAD + PNU  5XFAD + RJR  5XFAD + PNU + RJR | | 0.076 ± 0.070  0.090 ± 0.048  0.053 ± 0.033  0.065 ± 0.051  0.028 ± 0.022  0.055 ± 0.039  0.086 ± 0.036  0.078 ± 0.041 | |
| Fixed effects (type III) | P value | | F (DFn, DFd) | |
| Genotype | 0.0107 | | F (1, 267) = 6.604 | |
| Section | <0.0001 | | F (3, 279) = 10.02 | |
| Genotype x Section | <0.0001 | | F (3, 279) = 26.71 | |
| Tukey's multiple comparisons test | Mean diff. | 95.00% CI of diff. | | Adjusted P Value |
| WT |  |  | |  |
| Saline vs. PNU | -0.02545 | -0.04790 to -0.003008 | | 0.0191 |
| Saline vs. RJR | 0.01421 | -0.005747 to 0.03417 | | 0.2567 |
| Saline vs. PNU+RJR | 0.01091 | -0.005441 to 0.02726 | | 0.3129 |
| PNU vs. RJR | 0.03967 | 0.01695 to 0.06239 | | <0.0001 |
| PNU vs. PNU+RJR | 0.03636 | 0.01550 to 0.05722 | | <0.0001 |
| RJR vs. PNU+RJR | -0.003305 | -0.02182 to 0.01521 | | 0.9673 |
| 5XFAD |  |  | |  |
| Saline vs. PNU | -0.02663 | -0.04447 to -0.008802 | | 0.0008 |
| Saline vs. RJR | -0.05564 | -0.07446 to -0.03682 | | <0.0001 |
| Saline vs. PNU+RJR | -0.04664 | -0.06140 to -0.03188 | | <0.0001 |
| PNU vs. RJR | -0.02900 | -0.04958 to -0.008429 | | 0.0018 |
| PNU vs. PNU+RJR | -0.02001 | -0.03811 to -0.001901 | | 0.0237 |
| RJR vs. PNU+RJR | 0.008998 | -0.009927 to 0.02792 | | 0.6090 |
| Saline |  |  | |  |
| WT vs. 5XFAD | 0.04454 | 0.03117 to 0.05792 | | <0.0001 |
| PNU |  |  | |  |
| WT vs. 5XFAD | 0.04336 | 0.02460 to 0.06212 | | <0.0001 |
| RJR |  |  | |  |
| WT vs. 5XFAD | -0.02531 | -0.04313 to -0.007486 | | 0.0055 |
| PNU+RJR |  |  | |  |
| WT vs. 5XFAD | -0.01301 | -0.02546 to -0.0005567 | | 0.0406 |
| **Male** | | | | |
|  | WT + Saline  WT + PNU  WT + RJR  WT + PNU + RJR  5XFAD + Saline  5XFAD + PNU  5XFAD + RJR  5XFAD + PNU + RJR | | 0.061 ± 0.061  0.107 ± 0.053  0.088 ± 0.076  0.076 ± 0.064  0.030 ± 0.039  0.097 ± 0.047  0.077 ± 0.057  0.056 ± 0.032 | |
| Fixed effects (type III) | P value | | F (DFn, DFd) | |
| Genotype | 0.0002 | | F (1, 574) = 14.12 | |
| Section | <0.0001 | | F (3, 574) = 29.44 | |
| Genotype x Section | 0.2437 | | F (3, 574) = 1.394 | |
| Tukey's multiple comparisons test | Mean diff. | | 95.00% CI of diff. | Adjusted *P* Value |
| WT |  | |  |  |
| Saline vs. PNU | -0.04548 | | -0.06696 to -0.02401 | <0.0001 |
| Saline vs. RJR | -0.02654 | | -0.04887 to -0.004218 | 0.0122 |
| Saline vs. PNU+RJR | -0.01467 | | -0.03350 to 0.004156 | 0.1862 |
| PNU vs. RJR | 0.01894 | | -0.006960 to 0.04484 | 0.2360 |
| PNU vs. PNU+RJR | 0.03081 | | 0.007864 to 0.05376 | 0.0033 |
| RJR vs. PNU+RJR | 0.01187 | | -0.01187 to 0.03562 | 0.5708 |
| 5XFAD |  | |  |  |
| Saline vs. PNU | -0.06721 | | -0.09214 to -0.04228 | <0.0001 |
| Saline vs. RJR | -0.04695 | | -0.07236 to -0.02155 | <0.0001 |
| Saline vs. PNU+RJR | -0.02577 | | -0.04574 to -0.005795 | 0.0052 |
| PNU vs. RJR | 0.02026 | | -0.01018 to 0.05069 | 0.3170 |
| PNU vs. PNU+RJR | 0.04144 | | 0.01537 to 0.06752 | 0.0003 |
| RJR vs. PNU+RJR | 0.02119 | | -0.005340 to 0.04771 | 0.1683 |
| Saline |  | |  |  |
| WT vs. 5XFAD | 0.03098 | | 0.01746 to 0.04451 | <0.0001 |
| PNU |  | |  |  |
| WT vs. 5XFAD | 0.009258 | | -0.01187 to 0.03038 | 0.3898 |
| RJR |  | |  |  |
| WT vs. 5XFAD | 0.01057 | | -0.01138 to 0.03252 | 0.3445 |
| PNU+RJR |  | |  |  |
| WT vs. 5XFAD | 0.01989 | | 0.003924 to 0.03585 | 0.0147 |

**Supplementary Table 17. Statistical analysis of fear conditioning in Fig. 6a.**

| **Female** | | | | | | | |
| --- | --- | --- | --- | --- | --- | --- | --- |
| Training | | | | Test | | | |
| WT + Saline  WT + PNU  WT + RJR  WT + PNU + RJR  5XFAD + Saline  5XFAD + PNU  5XFAD + RJR  5XFAD + PNU + RJR | | 0.349 ± 0.198  0.393 ± 0.098  0.433 ± 0.051  0.294 ± 0.078  0.585 ± 0.131  0.567 ± 0.108  0.523 ± 0.132  0.393 ± 0.142 | | WT + Saline  WT + PNU  WT + RJR  WT + PNU + RJR  5XFAD + Saline  5XFAD + PNU  5XFAD + RJR  5XFAD + PNU + RJR | | 0.530 ± 0.149  0.584 ± 0.102  0.667 ± 0.107  0.552 ± 0.191  0.492 ± 0.147  0.546 ± 0.223  0.577 ± 0.188  0.698 ± 0.137 | |
|  | P value | | Difference | SE of difference | t ratio | | df |
| WT + Saline  WT + PNU  WT + RJR  WT + PNU + RJR  5XFAD + Saline  5XFAD + PNU  5XFAD + RJR  5XFAD + PNU + RJR | 0.0072  0.0017  0.0153  0.0466  0.0823  0.7310  0.5159  0.0022 | | -0.1810  -0.1917  -0.2342  -0.2576  0.09294  0.02070  -0.05360  -0.3052 | 0.05236  0.02551  0.05757  0.09796  0.04862  0.05784  0.07887  0.06463 | 3.456  7.513  4.068  2.629  1.912  0.3579  0.6796  4.723 | | 9.000  4.000  4.000  5.000  11.00  7.000  8.000  7.000 |
| **Male** | | | | | | | |
| Training | | | | Test | | | |
| WT + Saline  WT + PNU  WT + RJR  WT + PNU + RJR  5XFAD + Saline  5XFAD + PNU  5XFAD + RJR  5XFAD + PNU + RJR | | 0.404 ± 0.184  0.448 ± 0.083  0.387 ± 0.096  0.359 ± 0.127  0.506 ± 0.115  0.438 ± 0.245  0.552 ± 0.114  0.517 ± 0.109 | | WT + Saline  WT + PNU  WT + RJR  WT + PNU + RJR  5XFAD + Saline  5XFAD + PNU  5XFAD + RJR  5XFAD + PNU + RJR | | 0.575 ± 0.256  0.637 ± 0.173  0.617 ± 0.111  0.520 ± 0.150  0.589 ± 0.191  0.283 ± 0.232  0.582 ± 0.171  0.731 ± 0.108 | |
|  | P value | | Difference | SE of difference | t ratio | | df |
| WT + Saline  WT + PNU  WT + RJR  WT + PNU + RJR  5XFAD + Saline  5XFAD + PNU  5XFAD + RJR  5XFAD + PNU + RJR | 0.0114  0.0206  0.0059  0.0043  0.4290  0.1186  0.7547  0.0093 | | -0.1712  -0.1892  -0.2304  -0.1612  -0.08235  0.1551  -0.03047  -0.2134 | 0.05643  0.06356  0.05526  0.03251  0.09712  0.08242  0.09103  0.04540 | 3.035  2.976  4.170  4.958  0.8479  1.882  0.3347  4.700 | | 11.000  7.000  6.000  5.000  6.000  5.000  4.000  4.000 |

**Supplementary Table 18. Statistical analysis of fear conditioning in Fig. 6b.**

| **Female** | | | | | | | |
| --- | --- | --- | --- | --- | --- | --- | --- |
|  | | WT + Saline  WT + PNU  WT + RJR  WT + PNU + RJR  5XFAD + Saline  5XFAD + PNU  5XFAD + RJR  5XFAD + PNU + RJR | | | | 36.785 ± 10.866%  41.245 ± 19.035%  32.724 ± 12.744%  37.833 ± 10.913%  14.188 ± 12.202%  15.751 ± 10.332%  14.836 ± 10.400%  43.664 ± 13.580% | |
| ANOVA table | SS (Type III) | DF | | MS | F (DFn, DFd) | | P value |
| Interaction | 3321 | 3 | | 1107 | F (3, 117) = 6.834 | | P=0.0003 |
| Row Factor | 6665 | 1 | | 6665 | F (1, 117) = 41.15 | | P<0.0001 |
| Column Factor | 6665 | 3 | | 2222 | F (3, 117) = 13.72 | | P<0.0001 |
| Residual | 18951 | 117 | | 162.0 |  | | |
| Tukey's multiple comparisons test | | Mean diff. | | | | 95.00% CI of diff. | Adjusted *P* Value |
| WT + Saline vs. WT + PNU + RJR | | | -4.460 | | | -18.01 to 9.094 | 0.9712 |
| WT + Saline vs. WT + PNU | | | 4.061 | | | -9.801 to 17.92 | 0.9851 |
| WT + Saline vs. WT + RJR | | | -1.048 | | | -15.67 to 13.57 | >0.9999 |
| WT + Saline vs. 5XFAD + Saline | | | 22.60 | | | 10.74 to 34.45 | <0.0001 |
| WT + Saline vs. 5XFAD + PNU + RJR | | | -6.879 | | | -19.91 to 6.156 | 0.7320 |
| WT + Saline vs. 5XFAD + PNU | | | 21.03 | | | 6.819 to 35.25 | 0.0003 |
| WT + Saline vs. 5XFAD + RJR | | | 21.95 | | | 8.670 to 35.23 | <0.0001 |
| WT + PNU + RJR vs. WT + PNU | | | 8.521 | | | -6.609 to 23.65 | 0.6624 |
| WT + PNU + RJR vs. WT + RJR | | | 3.412 | | | -12.42 to 19.24 | 0.9977 |
| WT + PNU + RJR vs. 5XFAD + Saline | | | 27.06 | | | 13.74 to 40.37 | <0.0001 |
| WT + PNU + RJR vs. 5XFAD + PNU + RJR | | | -2.419 | | | -16.79 to 11.96 | 0.9995 |
| WT + PNU + RJR vs. 5XFAD + PNU | | | 25.49 | | | 10.04 to 40.95 | <0.0001 |
| WT + PNU + RJR vs. 5XFAD + RJR | | | 26.41 | | | 11.81 to 41.01 | <0.0001 |
| WT + PNU vs. WT + RJR | | | -5.109 | | | -21.20 to 10.98 | 0.9764 |
| WT + PNU vs. 5XFAD + Saline | | | 18.54 | | | 4.906 to 32.17 | 0.0013 |
| WT + PNU vs. 5XFAD + PNU + RJR | | | -10.94 | | | -25.61 to 3.727 | 0.3014 |
| WT + PNU vs. 5XFAD + PNU | | | 16.97 | | | 1.248 to 32.70 | 0.0248 |
| WT + PNU vs. 5XFAD + RJR | | | 17.89 | | | 3.003 to 32.77 | 0.0075 |
| WT + RJR vs. 5XFAD + Saline | | | 23.65 | | | 9.245 to 38.05 | <0.0001 |
| WT + RJR vs. 5XFAD + PNU + RJR | | | -5.831 | | | -21.22 to 9.555 | 0.9389 |
| WT + RJR vs. 5XFAD + PNU | | | 22.08 | | | 5.685 to 38.48 | 0.0016 |
| WT + RJR vs. 5XFAD + RJR | | | 23.00 | | | 7.404 to 38.59 | 0.0003 |
| 5XFAD + Saline vs. 5XFAD + PNU + RJR | | | -29.48 | | | -42.26 to -16.69 | <0.0001 |
| 5XFAD + Saline vs. 5XFAD + PNU | | | -1.563 | | | -15.55 to 12.43 | >0.9999 |
| 5XFAD + Saline vs. 5XFAD + RJR | | | -0.6480 | | | -13.68 to 12.39 | >0.9999 |
| 5XFAD + PNU + RJR vs. 5XFAD + PNU | | | 27.91 | | | 12.91 to 42.91 | <0.0001 |
| 5XFAD + PNU + RJR vs. 5XFAD + RJR | | | 28.83 | | | 14.71 to 42.95 | <0.0001 |
| 5XFAD + PNU vs. 5XFAD + RJR | | | 0.9153 | | | -14.30 to 16.13 | >0.9999 |
| **Male** | | | | | | | |
|  | | WT + Saline  WT + PNU  WT + RJR  WT + PNU + RJR  5XFAD + Saline  5XFAD + PNU  5XFAD + RJR  5XFAD + PNU + RJR | | | | 38.938 ± 13.066%  32.414 ± 11.302%  38.651 ± 15.155%  38.346 ± 11.517%  16.177 ± 8.761%  15.558 ± 6.577%  14.648 ± 6.018%  43.434 ± 21.002% | |
| ANOVA table | SS (Type III) | DF | | MS | | F (DFn, DFd) | P value |
| Interaction | 5022 | 3 | | 1674 | | F (3, 107) = 10.82 | P<0.0001 |
| Row Factor | 5304 | 1 | | 5304 | | F (1, 107) = 34.29 | P<0.0001 |
| Column Factor | 1997 | 3 | | 665.8 | | F (3, 107) = 4.305 | P=0.0066 |
| Residual | 16549 | 107 | | 154.7 | |  | |
| Tukey's multiple comparisons test | | | Mean diff. | | | 95.00% CI of diff. | Adjusted *P* Value |
| WT + Saline vs. WT + PNU + RJR | | | 6.524 | | | -8.505 to 21.55 | 0.8804 |
| WT + Saline vs. WT + PNU | | | 0.2876 | | | -13.31 to 13.88 | >0.9999 |
| WT + Saline vs. WT + RJR | | | 0.5926 | | | -14.44 to 15.62 | >0.9999 |
| WT + Saline vs. 5XFAD + Saline | | | 22.76 | | | 11.88 to 33.65 | <0.0001 |
| WT + Saline vs. 5XFAD + PNU + RJR | | | -4.496 | | | -17.74 to 8.745 | 0.9654 |
| WT + Saline vs. 5XFAD + PNU | | | 23.38 | | | 9.786 to 36.97 | <0.0001 |
| WT + Saline vs. 5XFAD + RJR | | | 24.29 | | | 9.818 to 38.76 | <0.0001 |
| WT + PNU + RJR vs. WT + PNU | | | -6.237 | | | -23.19 to 10.72 | 0.9470 |
| WT + PNU + RJR vs. WT + RJR | | | -5.932 | | | -24.06 to 12.19 | 0.9717 |
| WT + PNU + RJR vs. 5XFAD + Saline | | | 16.24 | | | 1.366 to 31.11 | 0.0221 |
| WT + PNU + RJR vs. 5XFAD + PNU + RJR | | | -11.02 | | | -27.69 to 5.653 | 0.4581 |
| WT + PNU + RJR vs. 5XFAD + PNU | | | 16.86 | | | -0.09964 to 33.81 | 0.0525 |
| WT + PNU + RJR vs. 5XFAD + RJR | | | 17.77 | | | 0.09869 to 35.43 | 0.0477 |
| WT + PNU vs. WT + RJR | | | 0.3050 | | | -16.65 to 17.26 | >0.9999 |
| WT + PNU vs. 5XFAD + Saline | | | 22.47 | | | 9.055 to 35.89 | <0.0001 |
| WT + PNU vs. 5XFAD + PNU + RJR | | | -4.784 | | | -20.18 to 10.61 | 0.9788 |
| WT + PNU vs. 5XFAD + PNU | | | 23.09 | | | 7.395 to 38.79 | 0.0004 |
| WT + PNU vs. 5XFAD + RJR | | | 24.00 | | | 7.539 to 40.47 | 0.0004 |
| WT + RJR vs. 5XFAD + Saline | | | 22.17 | | | 7.298 to 37.04 | 0.0003 |
| WT + RJR vs. 5XFAD + PNU + RJR | | | -5.089 | | | -21.76 to 11.58 | 0.9809 |
| WT + RJR vs. 5XFAD + PNU | | | 22.79 | | | 5.832 to 39.74 | 0.0016 |
| WT + RJR vs. 5XFAD + RJR | | | 23.70 | | | 6.031 to 41.36 | 0.0017 |
| 5XFAD + Saline vs. 5XFAD + PNU + RJR | | | -27.26 | | | -40.32 to -14.20 | <0.0001 |
| 5XFAD + Saline vs. 5XFAD + PNU | | | 0.6189 | | | -12.80 to 14.04 | >0.9999 |
| 5XFAD + Saline vs. 5XFAD + RJR | | | 1.529 | | | -12.78 to 15.84 | >0.9999 |
| 5XFAD + PNU + RJR vs. 5XFAD + PNU | | | 27.88 | | | 12.48 to 43.27 | <0.0001 |
| 5XFAD + PNU + RJR vs. 5XFAD + RJR | | | 28.79 | | | 12.61 to 44.96 | <0.0001 |
| 5XFAD + PNU vs. 5XFAD + RJR | | | 0.9101 | | | -15.55 to 17.37 | >0.9999 |

**Supplementary Table 19. Statistical analysis of plaques in Fig. 7.**

| **Female** | | | |
| --- | --- | --- | --- |
| **Average number of plaques / mm^2^** | | | |
|  | Saline  PNU  RJR  PNU + RJR | 50.62 ± 13.21  49.71 ± 11.17  47.71 ± 10.28  32.19 ± 7.19 | |
| ANOVA summary |  |  | |
| F | 7.497 |  |  |
| P value | 0.0006 |  |  |
| P value summary | *** |  |  |
| Significant diff. among means (P < 0.05)? | Yes |  |  |
| R squared | 0.3981 |  | |
| Tukey's multiple comparisons test | Mean diff. | 95.00% CI of diff. | Adjusted P Value |
| Saline vs. PNU | 0.9117 | -12.55 to 14.37 | 0.9978 |
| Saline vs. RJR | 2.916 | -10.54 to 16.37 | 0.9359 |
| Saline vs. PNU + RJR | 18.43 | 6.280 to 30.58 | 0.0013 |
| PNU vs. RJR | 2.004 | -12.18 to 16.19 | 0.9808 |
| PNU vs. PNU + RJR | 17.52 | 4.566 to 30.47 | 0.0046 |
| RJR vs. PNU + RJR | 15.51 | 2.562 to 28.46 | 0.0137 |
| **Male** | | | |
| **Average number of plaques / mm^2^** | | | |
|  | Saline  PNU  RJR  PNU + RJR | 38.95 ± 13.53  38.66 ± 8.85  47.88 ± 5.97  24.82 ± 9.89 | |
| ANOVA summary |  |  | |
| F | 10.57 |  |  |
| P value | <0.0001 |  |  |
| P value summary | **** |  |  |
| Significant diff. among means (P < 0.05)? | Yes |  |  |
| R squared | 0.4360 |  | |
| Tukey's multiple comparisons test | Mean diff. | 95.00% CI of diff. | Adjusted P Value |
| Saline vs. PNU | 0.2931 | -12.33 to 12.92 | >0.9999 |
| Saline vs. RJR | -8.926 | -21.55 to 3.700 | 0.2469 |
| Saline vs. PNU + RJR | 14.14 | 3.706 to 24.57 | 0.0042 |
| PNU vs. RJR | -9.220 | -23.05 to 4.612 | 0.2951 |
| PNU vs. PNU + RJR | 13.84 | 1.982 to 25.70 | 0.0165 |
| RJR vs. PNU + RJR | 23.06 | 11.20 to 34.92 | <0.0001 |
